# The Geometry and Dimensionality of Brain-wide Activity

**DOI:** 10.1101/2023.02.23.529673

**Authors:** Zezhen Wang, Weihao Mai, Yuming Chai, Kexin Qi, Hongtai Ren, Chen Shen, Shiwu Zhang, Guodong Tan, Yu Hu, Quan Wen

## Abstract

Understanding neural activity organization is vital for deciphering brain function. By recording whole-brain calcium activity in larval zebrafish during hunting and spontaneous behaviors, we find that the shape of the neural activity space, described by the neural covariance spectrum, is scale-invariant: a smaller, *randomly sampled* cell assembly resembles the entire brain. This phenomenon can be explained by Euclidean Random Matrix theory, where neurons are reorganized from anatomical to functional positions based on their correlations. Three factors contribute to the observed scale invariance: slow neural correlation decay, higher functional space dimension, and neural activity heterogeneity. In addition to matching data from zebrafish and mice, our theory and analysis demonstrate how the geometry of neural activity space evolves with population sizes and sampling methods, thus revealing an organizing principle of brain-wide activity.

## 1 Introduction

Geometric analysis of neuronal population activity has revealed the fundamental structures of neural representations and brain dynamics (1–4). Dimensionality reduction methods, which identify collective or latent variables in neural populations, simplify our view of high-dimensional neural data (5). Their applications to optical and multi-electrode recordings have begun to reveal important mechanisms by which neural cell assemblies process sensory information (6, 7), make decisions (8, 9), maintain working memory (10) and generate motor behaviors (1, 11–13).

In the past decade, the number of neurons that can be simultaneously recorded *in vivo* has grown exponentially (11, 14–21). This increase spans various brain regions (6, 16, 22) and the entire mammalian brain (23, 24). As more neurons are recorded, the multidimensional neural activity space, with each axis representing a neuron’s activity level (Fig. 1A), becomes more complex. The changing size of observed cell assemblies raises a number of basic questions. How does this space’s geometry evolve and what structures remain invariant with increasing number of neurons recorded? A key measure, the *effective dimension* or *participation ratio* (denoted as *D*_PR_, Fig. 1B), captures a major part of variability in neural activity (25–29). How does *D*_PR_ vary with the number of sampled neurons (Fig. 1A)? Two scenarios are possible: *D*_PR_ grows continuously with more sampled neurons; *D*_PR_ saturates as the sample size increases. Which scenario fits the brain? Furthermore, even if two cell assemblies have the same *D*_PR_, they can have different shapes (the geometric configuration of the neural activity space, as dictated by the eigenspectrum of the covariance matrix, Fig. 1C). How does the shape vary with the number of neurons sampled? Lastly, are we going to observe the same picture of neural activity space when using different recording methods such as two-photon microscopy, which records all neurons in a brain region, versus Neuropixels (16), which conducts a broad random sampling of neurons?

**Figure 1.**
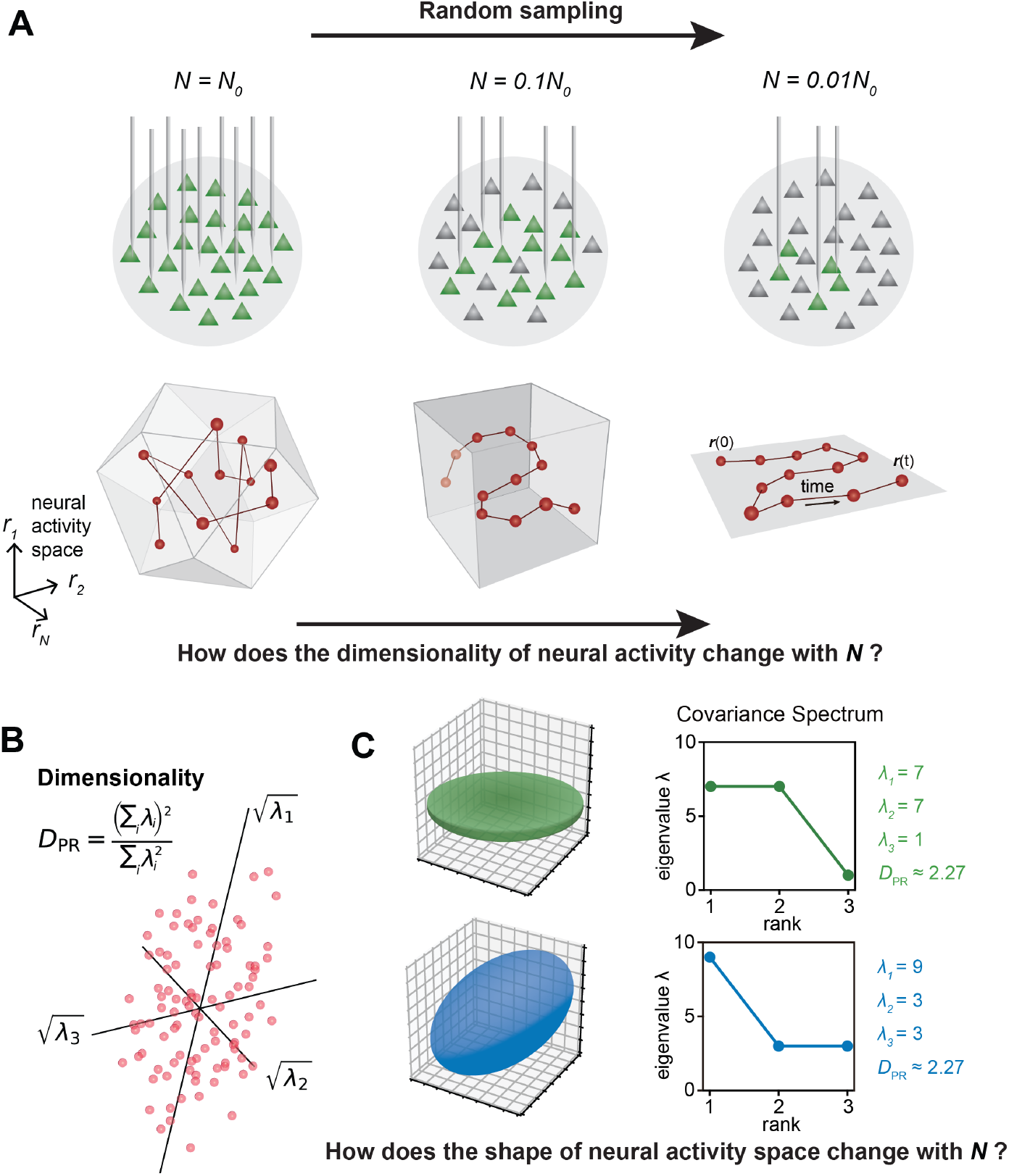
The relationship between the geometric properties of the neural activity space and the size of neural assemblies. **A.** Illustration of how dimensionality of neural activity (*D*_PR_) changes with the number of recorded neurons. **B**. The eigenvalues of the neural covariance matrix dictate the geometrical configuration of the neural activity space with 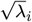 being the distribution width along a principal axis. **C**. Examples of two neural populations with identical dimensionality (*D*_PR_ = 25/11 ≈ 2.27) but different spatial configurations, as revealed by the eigenvalue spectrum (green: {λ_i_} = {7, 7, 1}, blue: {λ_i_} = {9, 3, 3}).

Here, we aim to address these questions by analyzing brain-wide Ca^2+^ activity in larval zebrafish during hunting or spontaneous behavior (Fig. 2A) recorded by Fourier light-field microscopy (30). The small size of this vertebrate brain, together with the volumetric imaging method, enables us to capture a significant amount of neural activity across the entire brain simultaneously. To characterize the geometry of neural activity beyond its dimensionality *D*_PR_, we examine the eigenvalues or spectrum of neural covariance (31) (Fig. 1C). The covariance spectrum has been instrumental in offering mechanistic insights into neural circuit structure and function, such as the effective strength of local recurrent interactions and the depiction of network motifs (29, 31, 32). Intriguingly, we find that both the dimensionality and covariance spectrum remain invariant for cell assemblies that are randomly selected from various regions of the zebrafish brain. We also verify this observation in datasets recorded by different experimental methods, including light-sheet imaging of larval zebrafish (33), two-photon imaging of mouse visual cortex (23), and multi-area Neuropixels recording in the mouse (23). To explain the observed phenomenon, we model the covariance matrix of brain-wide activity by generalizing the Euclidean Random Matrix (ERM) (34) such that neurons correspond to points distributed in a *d*-dimensional functional or feature space, with pairwise correlation decaying with distance. The ERM theory, studied in theoretical physics (34, 35), provides extensive analytical tools for a deep understanding of the neural covariance matrix model, allowing us to unequivocally identify three crucial factors for the observed scale invariance.

**Figure 2.**
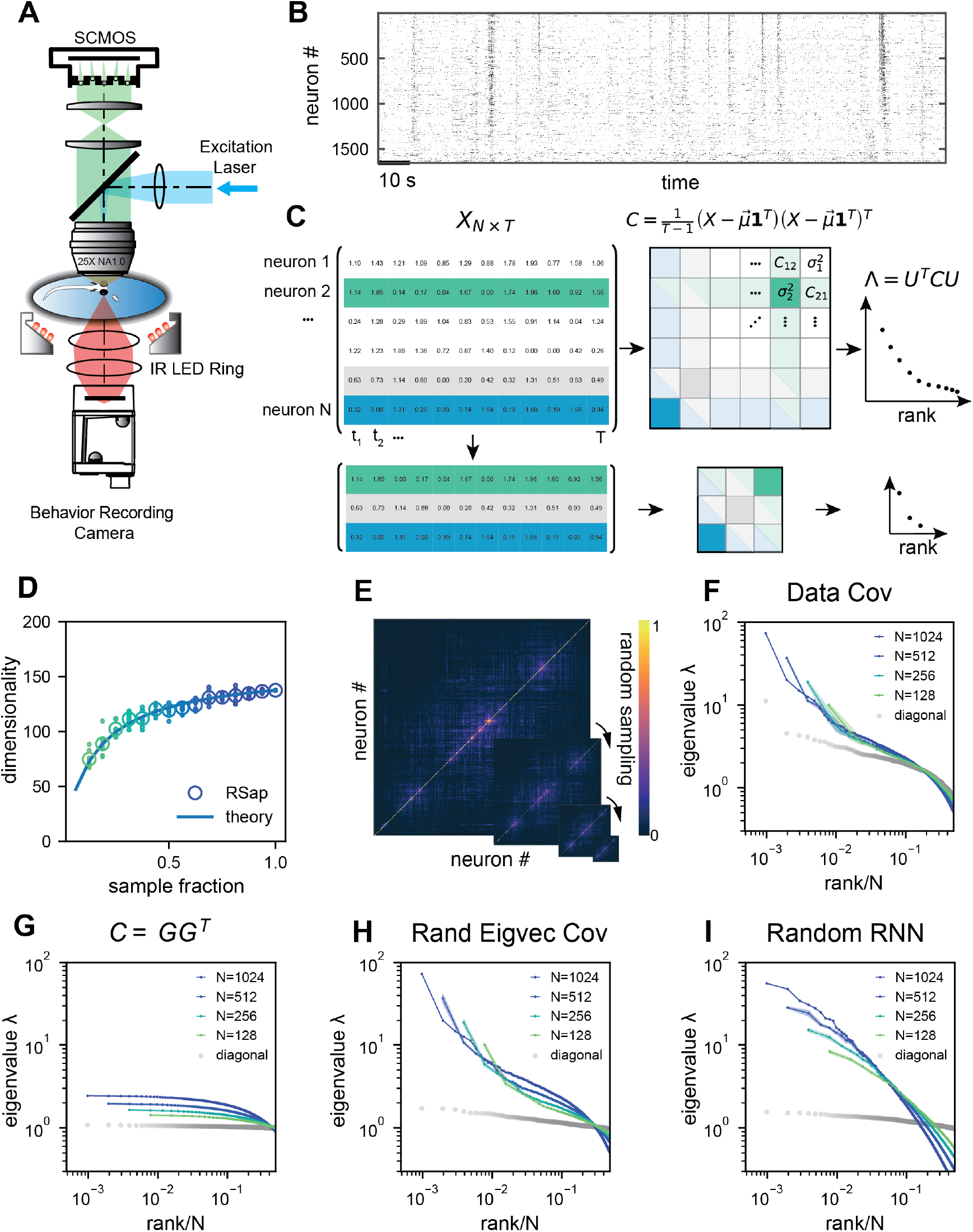
Whole-brain calcium imaging of zebrafish neural activity and the phenomenon of its scale-invariant covariance eigenspectrum. **A.** Rapid light-field Ca^2+^ imaging system for whole brain neural activity in larval zebrafish. **B**. Inferred firing rate activity from the brain-wide calcium imaging. The ROIs are sorted by their weights in the first principal component (23). **C**. Procedure of calculating the covariance spectrum on the full and sampled neural activity matrices. **D**. Dimensionality (circles, average across 8 samplings (dots)), as a function of the sampling fraction. The curve is the predicted dimensionality using Eq. (5). **E**. Iteratively sampled covariance matrices. Neurons are sorted in each matrix to maximize values near the diagonal. **F**. The covariance spectra, i.e., eigenvalue vs. rank/N, for randomly sampled neurons of different sizes (colors). The gray dots represent the sorted variances *C*_*ii*_ of all neurons. **G to I**. Same as **F** but from three models of covariance (see details in Methods): (G) a Wishart random matrix calculated from a random activity matrix of the same size as the experimental data; (H) replacing the eigenvectors by a random orthogonal set; (I) covariance generated from a randomly connected recurrent network.

Building upon our theoretical results, we further explore the connection between the spatial arrangement of neurons and their locations in functional space, which allows us to distinguish among three sampling approaches: random sampling, anatomical sampling (akin to optical recording of all neurons within a specific region of the brain) and functional sampling (20). Our ERM theory makes distinct predictions regarding the scaling relationship between dimensionality and the size of cell assembly, as well as the shape of covariance eigenspetrum under various sampling methods. Taken together, our results offer a new perspective for interpreting brain-wide activity and unambiguously show its organizing principles, with unexplored consequences for neural computation.

## 2 Results

### 2.1 Geometry of neural activity across random cell assemblies in zebrafish brain

We recorded brain-wide Ca^2+^ activity at a volume rate of 10 Hz in head-fixed larval zebrafish (Fig. 2A) during hunting attempts (Methods) and spontaneous behavior using a Fourier light field microscopy (30). Approximately 2000 ROIs (1977.3 ± 677.1, mean ± SD) with a diameter of 16.84 ± 8.51 µm were analyzed per fish based on voxel activity (Methods). These ROIs likely correspond to multiple nearby neurons with correlated activity. Henceforth, we refer to the ROIs as “neurons” for simplicity.

We first investigate the dimensionality of neural activity *D*_PR_ (Fig. 1B) in a randomly chosen cell assembly in zebrafish, similar to multi-area Neuropixels recording in a mammalian brain. We focus on how *D*_PR_ changes with a large sample size *N*. We find that if the mean squared covariance remains finite instead of vanishing with *N*, the dimensionality *D*_PR_ (Fig. 1B) becomes sample size independent and depends only on the variance 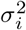 and the covariance *C*_*ij*_ between neurons *i* and *j*:

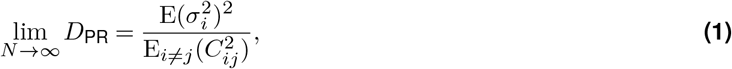

where E(…) denotes average across neurons (Methods and (29)). The finite mean squared covariance condition is supported by the observation that the neural activity covariance *C*_*ij*_ is positively biased and widely distributed with a long tail (Fig. S2A). As predicted, the data dimensionality grows with sample size and reaches the maximum value specified by Eq. (1) (Fig. 2D).

Next, we investigate the shape of the neural activity space described by the eigenspectrum of the covariance matrix derived from the activity of *N* randomly selected neurons (Fig. 2C). When the eigenvalues are arranged in descending order and plotted against the normalized rank *r*/*N*, where *r* = 1, …, *N*, (we refer to it as the *rank plot*), this curve shows an approximate power law that spans 10 folds. Interestingly, as the size of the covariance matrices decreases (*N* decreases), the eigenspectrum curves nearly collapse over a wide range of eigenvalues. This pattern holds across diverse datasets and experimental techniques (Fig. 2F, Fig. S2E-L). The similarity of the covariance matrices of randomly sampled neural populations can be intuitively visualized (Fig. 2E), after properly sorting the neurons (Methods).

The scale invariance in the neural covariance matrix – the collapse of the covariance eigenspectrum under random sampling – is non-trivial. The spectrum is not scale-invariant in a common covariance matrix model based on independent noise (Fig. 2G). It is absent when replacing the neural covariance matrix eigenvectors with random ones, keeping the eigenvalues identical (Fig. 2H). A recurrent neural network with random connectivity (31) does not yield a scale-invariant covariance spectrum (Fig. 2I). A recently developed latent variable model (36) (Fig. S23), which is able to reproduce avalanche criticality, also fails to generate the scale-invariant covariance spectrum. Thus, a new model is needed for the covariance matrix of neural activity.

### 2.2 Modeling covariance by organizing neurons in functional space

Dimension reduction methods simplify and visualize complex neuron interactions by embedding them into a low-dimensional map, within which nearby neurons have similar activities. Inspired by these ideas, we use the Euclidean Random Matrix (ERM (34)) to model neural covariance. Imagine sprinkling neurons uniformly distributed on a d-dimensional *functional space* of size *L* (Fig. 3A), where the distance between neurons *i* and *j* affects their correlation. Let 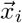 represent the functional coordinate of the neuron *i*. The distance-correlation dependency is described by *kernel function* 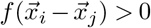 with *f* (0) = 1, indicating closer neurons have stronger correlations, and decreases as distance 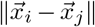 increases (Fig. 3A and Methods). To model the covariance, we extend the ERM by incorporating heterogeneity of neuron activity levels (shown as the size of the neuron in the functional space in Fig. 3A)

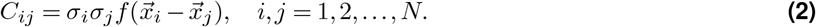

The variance of neural activity 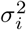 is drawn i.i.d. from a given distribution and is independent of neurons’ position.

**Figure 3.**
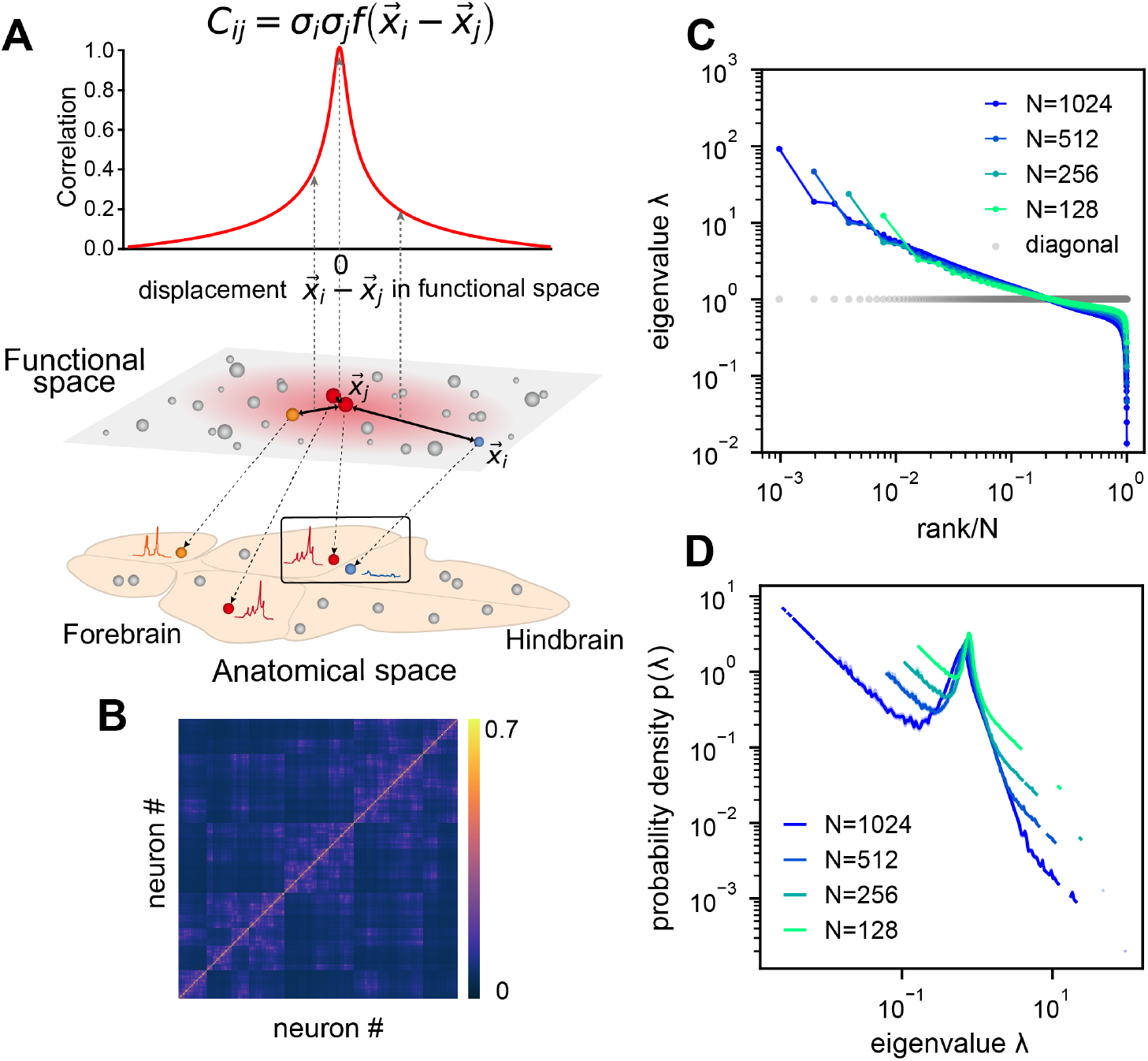
ERM model of covariance and its eigenspectrum. **A.** Schematic of the Euclidean Random Matrix (ERM) model, which reorganizes neurons (circles) from the anatomical space to the functional space (here *d* = 2 is a two-dimensional box). The correlation between a pair of neurons decreases with their distance in the functional space according to a kernel function 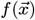. This correlation is then scaled by neurons’ variance 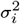 (circle size) to obtain the covariance *C*_*ij*_. **B**. An example ERM correlation matrix (i.e., when 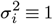). **C**. Spectrum (same as Fig. 2F) for the ERM correlation matrix in B. The gray dots represent the sorted variances C_ii_ of all neurons (same as in Fig. 2F). **D**. Visualizing the distribution of the same ERM eigenvalues in **C** by plotting the probability density function (pdf).

This multidimensional functional space may represent attributes to which neurons are tuned, such as sensory features (e.g., visual orientation (37), auditory frequency) and movement characteristics (e.g., direction, speed (38, 39)). In sensory systems, it represents stimuli as neural activity patterns, with proximity indicating similarity in features. For motor control, it encodes movement parameters and trajectories. In the hippocampus, it represents the place field of a place cell, acting as a cognitive map of physical space (40–42).

We first explore the ERM with various forms of 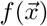 and find that fast-decaying functions like Gaussian and exponential functions do not produce eigenspectra similar to the data and no scale invariance over random sampling (Fig. S4A-H and Supp. Note). Thus, we turn to slow-decaying functions including the power law, which produce spectra similar to the data (Fig. 3C,D; see also Fig. S5). We adopt a particular kernel function because of its closed-form and analytical properties: 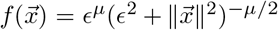 (Methods). For large distance 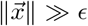, it approximates a power law 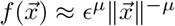 and smoothly transitions at small distance to satisfy the correlation requirement *f* (0) = 1 (Fig. S7I, J).

### 2.3 Analytical theory on the conditions of scale invariance in ERM

To determine the conditions for scale invariance in ERM, we analytically calculate the eigenspectrum of covariance matrix *C* (Eq. (2)) for large *N, L* using the replica method (34). A key order parameter emerging from this calculation is the neuron density *ρ*:= *N*/*L*^d^. In the high-density regime 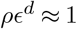, the covariance spectrum can be approximated in a closed form (Methods). For the slow-decaying kernel function 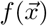 defined above, the spectrum for large eigenvalues follows a power law (Supp. Note):

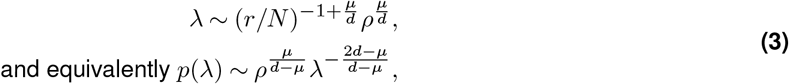

where *r* is the rank of the eigenvalues in descending order and *p*(λ) is their probability density function. Eq. (3) intuitively explains the scale invariance over random sampling. Sampling in the ERM reduces the neuron density *ρ*. The eigenspectrum is *ρ*-independent whenever *µ/d* ≈ 0. This indicates two factors contributing to the scale invariance of the eigenspectrum. First, a small exponent *µ* in the kernel function 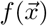 means that pairwise correlations slowly decay with functional distance and can be significantly positive across various functional modules and throughout the brain. For a given *µ*, an increase in dimension *d* improves the scale invariance. The dimension *d* could represent the number of independent features or latent variables describing neural activity or cognitive states.

We verify our theoretical predictions by comparing sampled eigenspectra in finite-size simulated ERMs across different *µ* and *d* (Fig. 4A). We first consider the case of homogeneous neurons (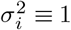 in Eq. (2), revisited later) in these simulations (Fig. 3C, D and Fig. 4A), making *C*’s entries correlation coefficients. To quantitatively assess the level of scale invariance, we introduce a *collapse index* (CI, see Methods for a detailed definition). Motivated by Eq. (3), the CI measures the shift of the eigenspectrum when the number of sampled neurons changes. The smaller CI values indicate higher scale invariance. Intuitively, it is defined as the area between spectrum curves from different sample sizes (Fig. 4A upper right). In the log-log scale rank plot, Eq. (3) shows the spectrum shifts vertically with ρ. Thus, we define CI as this average displacement (Fig. 4A upper right, Methods), and a smaller CI means more scale-invariant. Using CI, Fig. 4A shows that scale invariance improves with slower correlation decay as *µ* decreases and the functional dimension d increases. Conversely, with large *µ* and small d, the covariance eigenspectrum varies significantly with scale (Fig. 4A).

**Figure 4.**
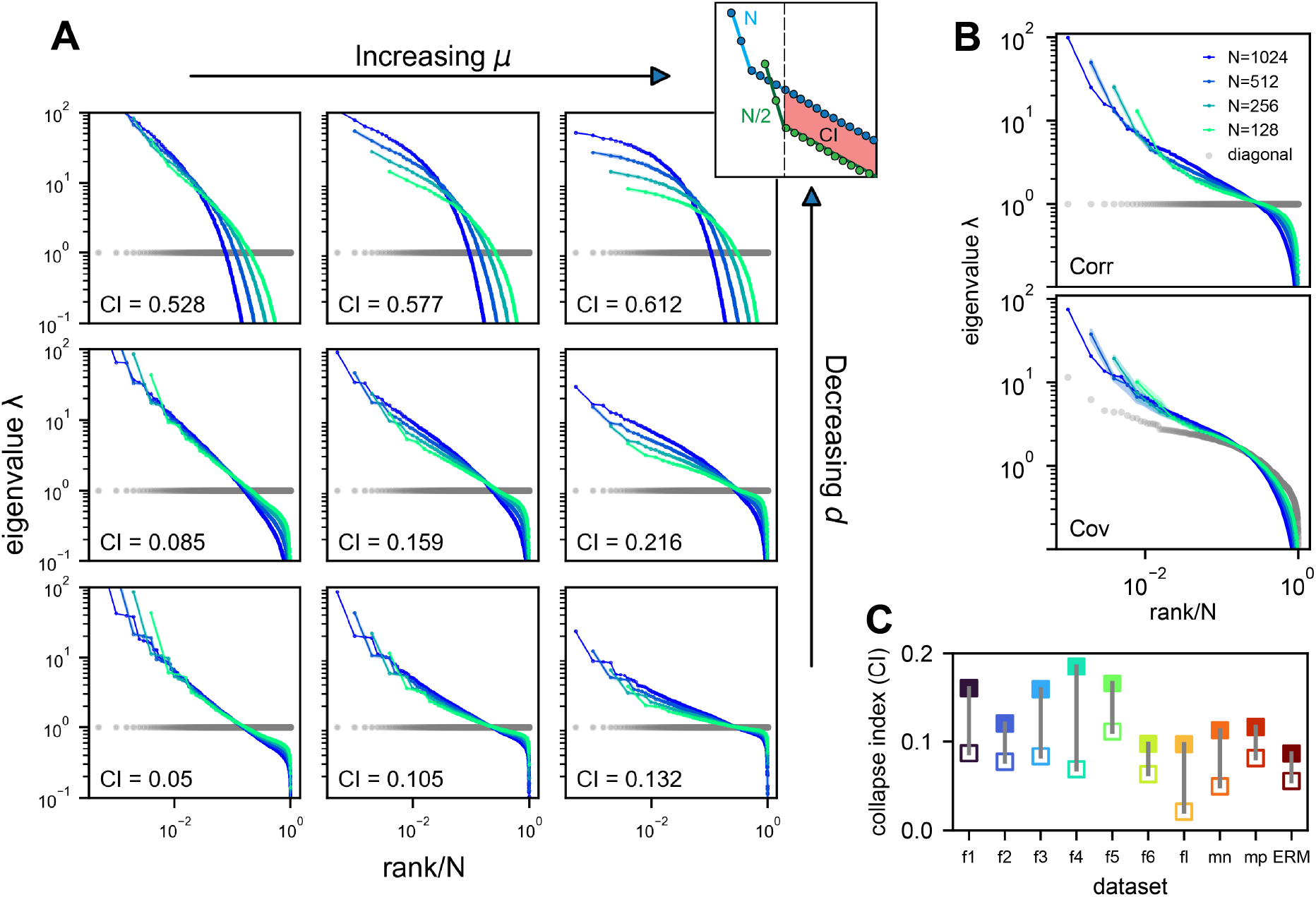
Three factors contributing to scale invariance. **A.** Impact of µ and *d* (see text) on the scale invariance of ERM spectrum (same plots as Fig. 3C) with 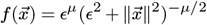. The degree of scale invariance is quantified by the collapse index (CI), which essentially measures the area between different spectrum curves (upper right inset). For comparison, we fix the same coordinate range across panels hence some plots are cropped. The gray dots represent the sorted variances C_*ii*_ of all neurons (same as in Fig. 2F). **B**. Top: sampled correlation matrix spectrum in an example animal (fish 1). Bottom: Same as top but for the covariance matrix that incorporates heterogeneous variances. The gray dots represent the sorted variances *C*_*ii*_ of all neurons (same as in Fig. 2F). **C**. The CI of the correlation matrix (filled squares) is found to be larger than that for the covariance matrix (opened squares) across different datasets: f1 to f6: six light-field zebrafish data (10 Hz per volume, this paper); fl: light-sheet zebrafish data (2 Hz per volume, (33)); mn: mouse Neuropixels data (downsampled to 10 Hz per volume); mp: mouse two-photon data, (3 Hz per volume, (23)).

Next, we consider the general case of unequal neural activity levels 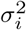 and check for differences between the correlation (equivalent to 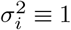) and covariance matrix spectra. Using the collapsed index (CI), we compare the scale invariance of the two spectra in the experimental data. Intriguingly, the CI of the covariance matrix is consistently smaller (more scale-invariant) across all datasets (Fig. 4C, Fig. S6C, open vs. closed squares), indicating that the heterogeneity of neuronal activity variances significantly affects the eigenspectrum and the geometry of neural activity space (43). By extending our spectrum calculation to the intermediate density regime 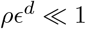 (Methods), we show that the ERM model can quantitatively explain the improved scale invariance in the covariance matrix compared to the correlation matrix (Fig. S6B).

Lastly, we examine factors that turn out to have minimal impact on the scale invariance of the covariance spectrum. First, the shape of the kernel function 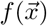 over a small distance (small distance means *f(x)* near *x* = 0 in the functional space, Fig. S7) does not affect the distribution of large eigenvalues (Fig. S7, table S3, Fig. S9A). This supports our use of a specific 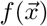 to represent a class of slow-decaying kernels. Second, altering the spatial distribution of neurons in the functional space, whether using a Gaussian, uniform, or clustered distribution, does not affect large covariance eigenvalues, except possibly the leading ones (Fig. S9B, Methods). Third, different geometries of the functional space, such as a flat square, a sphere, or a hemisphere, result in eigenspectra similar to the original ERM model (Fig. S9C). These findings indicate that our theory for the covariance spectrum’s scale invariance is robust to various modeling details.

### 2.4 Connection among random sampling, functional sampling, and anatomical sampling

So far, we have focused on random sampling of neurons, but how does the neural activity space change with different sampling methods? To this end, we consider three methods (Fig. 5A1): random sampling (RSap), anatomical sampling (ASap) where neurons in a brain region are captured by optical imaging (6, 44, 45), and functional sampling (FSap) where neurons are selected based on activity similarity (20). In ASap or FSap, sampling involves expanding regions of interest in anatomical space or functional space while measuring all neural activity within those regions (Methods). The difference among sampling methods depends on the neuron organization throughout the brain. If anatomically localized neurons also cluster functionally (Fig. 5A4), ASap ≈ FSap; if they are spread in the functional space (Fig. 5A2), ASap ≈ RSap. Generally, the anatomical-functional relationship is in-between and can be quantified using the Canonical Correlation Analysis (CCA). This technique finds axes (CCA vectors 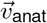 and 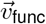) in anatomical and functional spaces such that the neurons’ projection along these axes has the maximum correlation, *R*_CCA_. The extreme scenarios described above correspond to *R*_CCA_ = 1 and *R*_CCA_ = 0.

**Figure 5.**
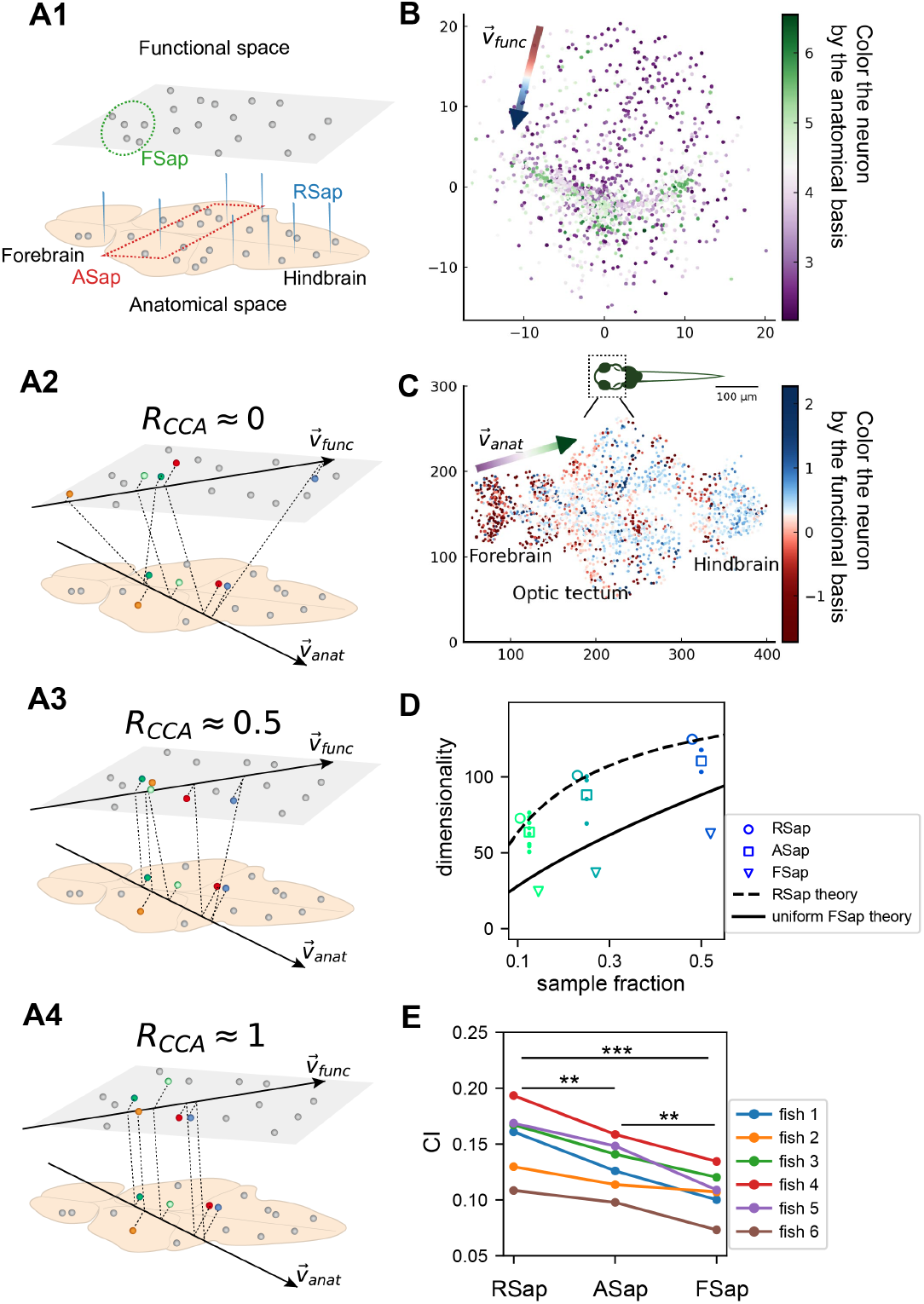
The relationship between the functional and anatomical space and theoretical predictions. **A.** Three sampling methods (A1) and *R*_CCA_ (see text). When *R*_CCA_ ≈ 0 (A2), the anatomical sampling (ASap) resembles the random sampling (RSap), and while when *R*_CCA_ ≈ 1 (A4), ASap is similar to the functional sampling (FSap). **B**. Distribution of neurons in the functional space inferred by MDS. Each neuron is color-coded by its projection along the first canonical direction 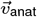 in the anatomical space (see text). Data based on fish 6, same for C to E. **C**. Similar to B. but plotting neurons in the anatomical space with color based on their projection along 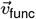 in the functional space (see text). **D**. Dimensionality (*D*_PR_) across sampling methods: average D_PR_ under RSap (circles), average and individual brain region *D*_PR_ under ASap (squares and dots), and *D*_PR_ under FSap for the most correlated neuron cluster (triangles; Methods). Dashed and solid lines are theoretical predictions for *D*_PR_ under RSap and FSap, respectively (Methods). **E**. The CI of correlation matrices under three sampling methods in 6 animals (colors). ^**^p<0.01; ^***^p<0.001; one-sided paired t tests: RSap vs. ASap, p = 0.0010; RSap vs. FSap, p = 0.0004; ASap vs. FSap, p = 0.0014.

To determine the anatomical-functional relationship in neural data, we infer the functional coordinates 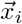 of each neuron by fitting the ERM using multidimensional scaling (MDS) (46) (Methods). For simplicity and better visualization, we use a low-dimensional functional space where *d* = 2. The fitted functional coordinates confirm the slow decay kernel function in ERM except for a small distance (Fig. S12). The ERM with inferred coordinates 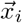 also reproduces the experimental covariance matrix, including cluster structures (Fig. S11) and its sampling eigenspectra (Fig. S10).

Equipped with the functional and anatomical coordinates, we next use CCA to determine which scenarios illustrated in Fig. 5A align better with the neural data. Fig. 5B,C shows a representative fish with a significant *R*_CCA_ = 0.327 (p-value=0.0042, Anderson–Darling test). Notably, the CCA vector in the anatomical space, 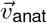, aligns with the rostrocaudal axis. Coloring each neuron in the functional space by its projection along 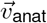 shows a correspondence between clustering and anatomical coordinates (Fig. 5B). Similarly, coloring neurons in the anatomical space (Fig. 5C) by their projection along 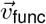 reveals distinct localizations in regions like the forebrain and optic tectum. Across animals, functionally clustered neurons show anatomical segregation (33), with an average *R*_CCA_ of 0.335±0.054 (mean±SD).

Next, we investigate the effects of different sampling methods (Fig. 5A1) on the geometry of the neural activity space when there is a significant but moderate anatomical-functional correlation as in the experimental data. Interestingly, dimensionality 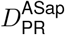 in data under anatomical sampling consistently falls between random and functional sampling values (Fig. 5D). This phenomenon can be intuitively explained by the ERM theory. Recall that for large *N*, the key term in Eq. (1) is 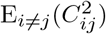. For a fixed number of sampled neurons, this average squared covariance is maximized when neurons are selected closely in the functional space (FSap) and minimized when distributed randomly (RSap). Thus, RSap and FSap D_PR_ set the upper and lower bounds of dimensionality, with ASap expected to fall in between. This reasoning can be precisely formulated to obtain quantitative predictions of the bounds (Methods). We predict the ASap dimension at large *N* as

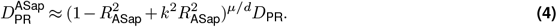

Here *D*_PR_ is the dimensionality under RSap (Eq. (1)), k represents the fraction of sampled neurons. R_ASap_ is the correlation between anatomical and functional coordinates along the direction where the anatomical subregions are divided (Methods), and it is bounded by the canonical correlation *R*_ASap_ ≤ *R*_CCA_. When *R*_ASap_ = 0, we get the upper bound 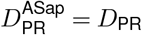 (Fig. 5D dashed line). The lower bound is reached when *R*_ASap_ = *R*_CCA_ = 1 (5A4), where Eq. (4) shows a scaling relationship 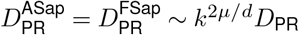 that depends on the sampling fraction *κ* (Fig. 5D solid line). This contrasts with the *κ*-independent dimensionality of RSap in Eq. (1). Furthermore, if *R*_ASap_ and its upper bound is not close to 1 (precisely *R*_ASap_ ≤ 0.84 for the ERM model in Fig. 5D), 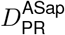 align closer to the upper bound of RSap. This prediction agrees well with our observations in data across animals (Fig. 5D, Fig. S20 and Fig. S21).

Beyond dimensionality, our theory predicts the difference in the covariance spectrum between sampling methods based on the neuronal density *ρ* in the functional space (Eq. (3)). This density *ρ* remains constant during FSap (Fig. 5A1) and decreases under RSap; the average density across anatomical regions ⟨*ρ*⟩ in ASap lies between those of FSap and RSap. Analogous to Eq. (4), the relationship in *ρ* orders the spectra: ASap’s spectrum lies between those of FSap and RSap (Methods). This further implies that the level of scale invariance under ASap should fall between that of RSap and FSap, which is confirmed by our experimental data (Fig. 5E).

## 3 Discussion

### Impact of hunting behavior on scale invariance and functional space organization

How does task-related neural activity shape the covariance spectrum and brain-wide functional organization? We examine the hunting behavior in larval zebrafish, marked by eye convergence (both eyes move inward to focus on the central visual field) (47). We find that scale invariance of the eigenspectra persists and is enhanced even after removing the hunting frames from the Ca^2+^ imaging data (Fig. 4C, Fig. S15AB, Methods). This is consistent with the scale-invariant spectrum found in other data sets during spontaneous behaviors (Fig. S10F, Fig. S2GH), suggesting scale invariance is a general phenomenon.

Interestingly, in the inferred functional space, we observe reorganizations of neurons after removing hunting behavior (Fig. S15C, D). Neurons in one cluster disperse from their center of mass (Fig. S15D) and decreases the local neuronal density ρ (Methods and Fig. S15E). The neurons in this dispersed cluster have a consistent anatomical distribution from the midbrain to the hindbrain in 4 out of 5 fish (Fig. S17). During hunting, the cluster has robust activations that are widespread in the anatomical space but localized in the functional space(Movie. S1).

Our findings suggest that the functional space could be defined by latent variables that represent cognitive factors such as decision-making, memory, and attention. These variables set the space’s dimensions, with neural activity patterns reflecting cognitive state dynamics. Functionally related neurons – through sensory tuning, movement parameters, internal conditions, or cognitive factors – become closer in this space, leading to stronger activity correlations.

### Criticality and power law

What drives brain dynamics with a slow-decaying distance-correlation function 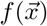 in functional space? Long-range connections and a slow decline in projection strength over distance (48) may cause extensive correlations, enhancing global activity patterns. This behavior is also reminiscent of phase transitions in statistical mechanics (49), where local interactions lead to expansive correlated behaviors. Studies suggest that critical brains optimize information processing (50, 51). The link between neural correlation structures and neuronal connectivity topology is an exciting area for future exploration.

In the high-density regime of the ERM model, the rank plot (Eq. (3)) for large eigenvalues (λ > 1) follows a power law λ ∼ *r*^−*α*^, with *α* = 1 − *µ*/*d* < 1. The scale invariant spectrum occurs when *α* is close to 1. Experimental data, however, align more closely with the model in the intermediate-density regime, where the power-law spectrum is an approximation and the decay is slower (for ERM model Fig. S3BC, and for data *α* = 0.47 ± 0.08, mean ± SD, *n* = 6 fish). Stringer et al. (6) found an *α* ≳ 1 decay in the mouse visual cortex’s stimulus trial averaged covariance spectrum, and they argued that this decay optimizes visual code efficiency and smoothness. Our study differs in two fundamental ways. First, we recorded brain-wide activity during spontaneous or hunting behavior, calculating neural covariance from single-trial activity. Much of the neural activity was not driven by sensory stimulus and unrelated to specific tasks, requiring a different interpretation of the neural covariance spectrum. Second, without loss of generality, we normalized the mean variance of neural activity E(*σ*^2^) by scaling the covariance matrix so that its eigenvalues sum up to *N*. This normalization imposes a constraint on the spectrum. In particular, large and small eigenvalues may have different behaviors and do not need to obey a single power law *λ* ∼ *r*^−*α*^ for all *N* eigenvalues (Methods). Stringer et al. (6) did not take this possibility into account, making their theory less applicable to our analysis.

We draw inspiration from the renormalization group (RG) approach to navigate neural covariance across scales, which has also been explored in the recent literature. Following Kadanoff’s block spin transformation (49), Meshulam et al. (20) formed size-dependent neuron clusters and their covariance matrices by iteratively pairing the most correlated neurons and placing them adjacent on a lattice. The groups expanded until the largest reached the system size. The RG process, akin to spatial sampling in functional space (FSap), maintains constant neuron density *ρ*. Thus, for any kernel function 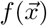, including the power law and exponential, the covariance eigenspectrum remains invariant across scales (Fig. S19A,B,D,E).

Morrell et al. (36, 52) proposed a simple model in which a few time-varying latent factors impact the whole neural population. We evaluated if this model could account for the scale invariance seen in our data. Simulations showed that the resulting eigenspectra differed considerably from our findings (Fig. S23). Although the Morrell model demonstrated a degree of scale invariance under functional sampling (or RG), it did not align with the scale-invariant features under random sampling, suggesting that this simple model might not capture all crucial features in our observations.

We emphasize that the covariance spectrum being a power law is distinct from the scale invariance we define in this study, namely the collapse of spectrum curves under random neuron sampling. The random RNN model in Fig. 2I shows a power-law behavior, but lacks true scale invariance as spectrum curves for different sizes do not collapse. When connection strength *g* approaches 1, the system exhibits a power law spectrum of 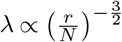. Subsampling causes the spectrum to shift by 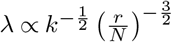, where *k* = *N*_*s*_/*N* is the sampling fraction (derived from Eq. 24 in(31)).

### Bounded dimensionality under random sampling

The saturation of the dimensionality D_PR_ at large sample sizes indicates a limit to neural assembly complexity, evidenced by the finite mean square covariance. This is in contrast with neural dynamics models such as the balanced excitatory-inhibitory (E-I) neural network (53), where 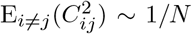 resulting in an unbounded dimensionality (see Supp. Note). Our results suggest that the brain encodes experiences, sensations, and thoughts using a finite set of dimensions instead of an infinitely complex neural activity space.

We found that the relationship between dimensionality and the number of recorded neurons depends on the sampling method. For functional sampling, the dimensionality scales with the sampling fraction 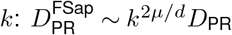. This suggests that if anatomically sampled neurons are functionally clustered, as with cortical neurons forming functional maps, the increase in dimensionality with neuron number may seem unbounded. This offers new insights for interpreting large-scale neural activity data recorded under various techniques.

Manley et al. (54) found that, unlike in our study, neural activity dimensionality in head-fixed, spontaneously behaving mice did not saturate. They used shared variance component analysis (SVCA) and noted that PCA-based estimates often show dimensionality saturation, which is consistent with our findings. We intentionally chose PCA in our study for several reasons. First, PCA is a trusted and widely used method in neuroscience, proven to uncover meaningful patterns in neural data. Second, its mathematical properties are well understood, making it particularly suitable for our theoretical analysis. Although newer methods such as SVCA might offer valuable insights, we believe PCA remains the most appropriate method for our research questions.

It’s important to note that the scale invariance of dimensionality and covariance spectrum are distinct phenomena with different underlying requirements. Dimensionality invariance relies on finite mean square covariance, causing saturation at large sample sizes. In contrast, spectral invariance requires a slow-decaying correlation kernel (small *µ*) and/or a high-dimensional functional space (large *d*). Although both features appear in our data, they result from distinct mechanisms. A neural system could show saturating dimensionality without spectral invariance if it has finite mean square covariance but rapidly decaying correlations with functional distance. Understanding these requirements clarifies how neural organization affects different scale-invariant properties.

### Computational benefits of a scale-invariant covariance spectrum

Our findings are validated across multiple datasets obtained through various recording techniques and animal models, ranging from single-neuron calcium imaging in larval zebrafish to single-neuron multi-electrode recordings in the mouse brain (see Fig. S2). The conclusion remains robust when the multi-electrode recording data are reanalyzed under different sampling rates (6 Hz - 24 Hz, Fig. S24). We also confirm that substituting a few negative covariances with zero retains the spectrum of the data covariance matrix (Fig. S18 and Methods).

The scale invariance of neural activity across different neuron assembly sizes could support efficient multiscale information encoding and processing. This indicates that the neural code is robust and requires minimal adjustments despite changes in population size. One recent study shows that randomly sampled and coarse-grained macrovoxels can predict population neural activity (55), reinforcing that a random neuron subset may capture overall activity patterns. This enables downstream circuits to readout and process information through random projections (27). A recent study demonstrates that a scale-invariant noise covariance spectrum with a specific slope *α* < 1 enables neurons to convey unlimited stimulus information as the population size increases (56). The linear Fisher information, in this context, grows at least as *N* ^1−*α*^.

Understanding how dimensionality and spectrum change with sample size also suggests the possibility of extrapolating from small samples to overcome experimental limitations. This is particularly feasible when *µ*/*d* → 0, where the dimensionality and spectrum under anatomical, random, and functional sampling coincide (Equations (3) and (4)). Developing extrapolation methods and exploring the benefits of scale-invariant neural code are promising future research directions.

## 4 Materials and Methods

### 4.1 Experimental methods

The handling and care of the zebrafish complied with the guidelines and regulations of the Animal Resources Center of the University of Science and Technology of China (USTC). All larval zebrafish (huc:h2b -GCaMP6f) were raised in E2 embryo medium (comprising 7.5 mM NaCl, 0.25 mM KCl, 0.5 mM MgSO_4_, 0.075 mM KH_2_PO_4_, 0.025 mM Na_2_HPO_4_, 0.5 mM CaCl_2_, and 0.35 mM NaHCO_3_; containing 0.5 mg/L methylene blue) at 28.5 °C and with a 14-h light and 10-h dark cycle.

To induce hunting behavior (composed of motor sequences like eye convergence and J turn) in larval zebrafish, we fed them a large amount of paramecia over a period of 4-5 days post-fertilization (dpf). The animals were then subjected to a 24-hour starvation period, after which they were transferred to a specialized experimental chamber. The experimental chamber was 20mm in diameter and 1mm in depth, and the head of each zebrafish was immobilized by applying 2% low melting point agarose. The careful removal of the agarose from the eyes and tail of the fish ensured that these body regions remained free to move during hunting behavior. Thus, characteristic behavioral features such as J-turns and eye convergence could be observed and analyzed. Subsequently, the zebrafish were transferred to an incubator and stayed overnight. At 7 dpf, several paramecia were introduced in front of the previously immobilized animals, each of which was monitored by a stereomicroscope. Those displaying binocular convergence were selected for subsequent Ca^2+^ imaging experiments.

We developed a novel optomagnetic system that allows (1) precise control of the trajectory of the paramecium and (2) imaging brain-wide Ca^2+^ activity during the hunting behavior of zebrafish. To control the movement of the paramecium, we treated these microorganisms with a suspension of ferric tetroxide for 30 minutes and selected those that responded to its magnetic attraction. A magnetic paramecium was then placed in front of a selected larva, and its movement was controlled by changing the magnetic field generated by Helmholtz coils that were integrated into the imaging system. The real-time position of the paramecium, captured by an infrared camera, was identified by online image processing. The positional vector relative to a predetermined target position was calculated. The magnitude and direction of the current in the Helmholtz coils were adjusted accordingly, allowing for precise control of the magnetic field and hence the movement of the paramecium. Multiple target positions could be set to drive the paramecium back and forth between multiple locations.

The experimental setup consisted of head-fixed larval zebrafish undergoing two different types of behavior: induced hunting behavior by a moving paramecium in front of a fish (fish 1-5), and spontaneous behavior without any visual stimulus as a control (fish 6). Experiments were carried out at ambient temperature (ranging from 23°C to 25°C). The behavior of the zebrafish was monitored by a high-speed infrared camera (Basler acA2000-165umNIR, 0.66x) behind a 4F optical system and recorded at 50 Hz. Brain-wide Ca^2+^ imaging was achieved using XLFM. Light-field images were acquired at 10 Hz, using customized LabVIEW software (National Instruments, USA) or Solis software (Oxford Instruments, UK), with the help of a high-speed data acquisition card (PCIe-6321, National Instruments, USA) to synchronize the fluorescence with behavioral imaging.

#### 4.1.1 Behavior analysis

The background of each behavior video was removed using the clone stamp tool in Adobe Photoshop CS6. Individual images were then processed by an adaptive thresholding algorithm, and fish head and yolk were selected manually to determine the head orientation. The entire body centerline, extending from head to tail, was divided into 20 segments. The amplitude of a bending segment was defined as the angle between the segment and the head orientation. To identify the paramecium in a noisy environment, we subtracted a background image, averaged over a time window of 100 s, from all the frames. The major axis of the left or right eye was identified using DeepLabCut (57). The eye orientation was defined as the angle between the rostrocaudal axis and the major axis of an eye; The convergence angle was defined as the angle between the major axes of the left and right eyes. An eye-convergence event was defined as a period of time where the angle between the long axis of the eyes stayed above 50 degrees (47).

**Table S1.** Table of notations.

| Notation | Description |
| --- | --- |
| $C$ | covariance matrix, Eq. (2) |
| $C_{ij}$ | pairwise covariance between neuron $i, j$; entries of $C$, section 4.3 |
| $D_{PR}$ | participation ratio dimension, Eq. (5) |
| $D_{PR}^{ASap}$ | anatomical sampling dimension, Eq. (4) |
| $\lambda$ | eigenvalue of a covariance matrix $C$ |
| $p(\lambda)$ | probability density function of covariance eigenvalues, Eq. (8) |
| $r$ | rank of an eigenvalue in descending order, Eq. (3) |
| $q$ | fraction of eigenvalues up to $\lambda$ and $q = r/N$, Eq. (13) |
| $f(\vec{x}) = f(\ \vec{x}_i - \vec{x}_j\ )$ | kernel function or distance-correlation function, Eq. (11) |
| $\tilde{f}(\vec{k})$ | Fourier transform of $f(\vec{x})$, $\tilde{f}(\vec{k}) = \int_{\mathbb{R}^d} f(\vec{x}) e^{-i\vec{x} \cdot \vec{k}} d^d \vec{x}$ |
| $\mu$ | power-law exponent in $f(\vec{x})$, Eq. (11) |
| $\epsilon$ | resolution parameter in $f(\vec{x})$ to smooth the singularity near 0, Eq. (11) |
| $N$ | number of neurons |
| $N_0$ | the total number of neurons prior to sampling |
| $k$ | $N/N_0$ the fraction of sampled neurons |
| $L$ | linear box size of the functional space |
| $\rho$ | density of neurons in the functional space |
| $d$ | dimension of the functional space |
| $a_i(t)$ | neural activity of neuron $i$ at time $t$ |
| $\sigma_i^2$ | temporal variance of neural activity, Eq. (2) |
| CI | collapse index for measuring scale invariance Eq. (13) |
| $\alpha$ | power-law coefficient of eigenspectrum in the rank plot, section 3 |
| $\vec{x}_i, \vec{y}_i$ | neuron $i$'s coordinate in the functional and anatomical space, respectively |
| $\vec{v}_{func}, \vec{v}_{anat}$ | the first canonical directions in the functional and anatomical space, respectively, section 2.4 |
| $R_{CCA}$ | the first canonical correlation, section 2.4 |
| $R_{ASap}$ | correlation between anatomical and functional coordinates along ASap direction |

#### 4.1.2 Imaging data acquisition and processing

We used a fast eXtended light field microscope (XLFM, with a volume rate of 10 Hz) to record Ca^2+^ activity throughout the brain of head-fixed larval zebrafish. Fish were ordered by the dates of experiments. As previously described (30), we adopted the Richardson-Lucy deconvolution method to iteratively reconstruct 3D fluorescence stacks (600 × 600 × 250) from the acquired 2D images (2048 × 2048). This algorithm requires an experimentally measured point spread function (PSF) of the XLFM system. The entire recording for each fish is 15.3±4.3 min (mean±SD).

To perform image registration and segmentation, we first cropped and resized the original image stack to 400 × 308 × 210, which corresponded to the size of a standard zebrafish brain (zbb) atlas (58). This step aimed to reduce substantial memory requirements and computational costs in subsequent operations. Next, we picked a typical volume frame and aligned it with the zbb atlas using a basic 3D affine transformation. This transformed frame was used as a template. We aligned each volume with the template using rigid 3D intensity-based registration (59) and non-rigid pairwise registration (60) in the Computational Morphometry Toolkit (CMTK) (https://www.nitrc.org/projects/cmtk/). After voxel registration, we computed the pairwise correlation between nearby voxel intensities and performed the watershed algorithm on the correlation map to cluster and segment voxels into consistent ROIs across all volumes. We defined the diameter of each ROI using the maximum Feret diameter (the longest distance between any two voxels within a single ROI).

Finally, we adopted the “OASIS” deconvolution method to denoise and infer neural activity from the fluorescence time sequence (61). The deconvolved Δ*F*/*F* of each ROI was used to infer firing rates for subsequent analysis.

**Table S2.**
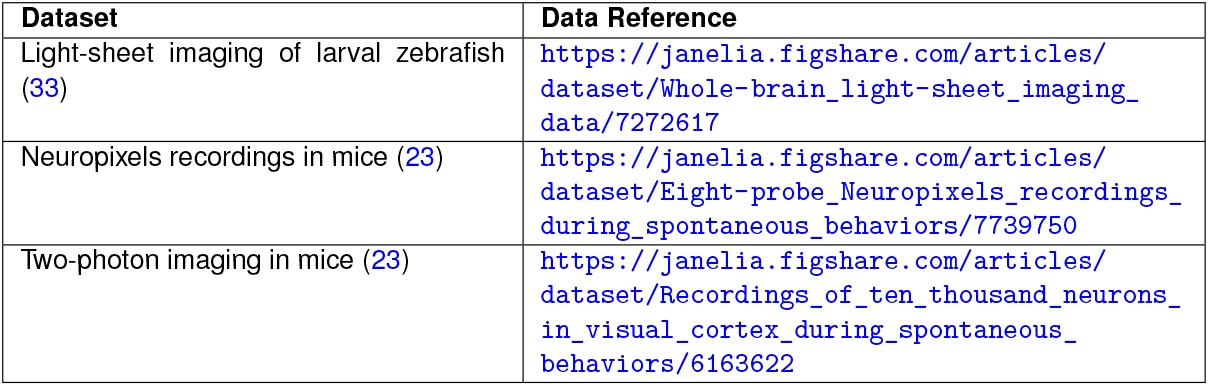
Resources for additional experimental datasets.

| Dataset | Data Reference |
| --- | --- |
| Light-sheet imaging of larval zebrafish (33) | https://janelia.figshare.com/articles/dataset/Whole-brain_light-sheet_imaging_data/7272617 |
| Neuropixels recordings in mice (23) | https://janelia.figshare.com/articles/dataset/Eight-probe_Neuropixels_recordings_during_spontaneous_behaviors/7739750 |
| Two-photon imaging in mice (23) | https://janelia.figshare.com/articles/dataset/Recordings_of_ten_thousand_neurons_in_visual_cortex_during_spontaneous_behaviors/6163622 |

### 4.2 Other experimental datasets analyzed

To validate our findings across different recording methods and animal models, we also analyzed three additional datasets. We include a brief description below for completeness. Further details can be found in the respective reference. The first dataset includes whole-brain light-sheet Ca^2+^ imaging of immobilized larval zebrafish in the presence of visual stimuli as well as in a spontaneous state (33). Each volume of the brain was scanned through 2.11 ± 0.21 planes per sec, providing a near-simultaneous readout of neuronal Ca^2+^ signals. We analyzed fish 8 (69,207 neurons × 7,890 frames), 9 (79,704 neurons × 7,720 frames) and 11 (101,729 neurons × 8,528 frames), which are the first three fish data with more than 7,200 frames. For simplicity, we labeled them l2, l3, and l1(fl). The second dataset consists of Neuropixels recordings from approximately ten different brain areas in mice during spontaneous behavior (23). Data from the three mice, *Kerbs, Robbins*, and *Waksman*, include the firing rate matrices of 1,462 neurons × 39,053 frames, 2,296 neurons × 66,409 frames, and 2,688 neurons × 74,368 frames, respectively. The last dataset comprises two-photon Ca^2+^ imaging data (2-3 Hz) obtained from the visual cortex of mice during spontaneous behavior. While this dataset includes numerous animals, we focused on the first three animals that exhibited spontaneous behavior: `spont_M150824_MP019_2016-04-05` (11,983 neurons × 21,055 frames), `spont_M160825_MP027_2016-12-12` (11,624 neurons × 23,259 frames), and `spont_M160907_MP028_2016-09-26` (9,392 neurons × 10,301 frames) (23).

### 4.3 Covariance matrix, eigenspectrum and sampling procedures

To begin, we multiplied the inferred firing rate of each neuron (see section 4.1.2) by a constant such that in the resulting activity trace a_*i*_, the mean of *a*_*i*_*(t) over the nonzero time frames* equaled one (20). Consistent with the literature (20), this step aimed to eliminate possible confounding factors in the raw activity traces, such as the heterogeneous expression level of the fluorescence protein within neurons and the non-linear conversion of the electrical signal to Ca^2+^ concentration. Note that after this scaling, neurons could still have different activity levels characterized by the variance of *a*_*i*_*(t)* over time, due to differences in the sparsity of activity (proportion of nonzero frames) and the distribution of nonzero *a*_*i*_*(t)* values. Without normalization, the covariance matrix becomes nearly diagonal, causing significant underestimation of the covariance structures.

The three models of covariance in Fig. 2G-I were constructed as follows. For model in Fig. 2G, the entries of matrix *G* (with dimensions *N* × *T*) were sampled from an i.i.d. Gaussian distribution with zero mean and standard deviation *σ* = 1. In Fig. 2H, we constructed the composite covariance matrix for fish 1 achieved by maintaining the eigenvalues from the fish 1 data covariance matrix and replacing the eigenvectors *U* with a set of random orthonormal basis. Lastly, the covariance matrix in Fig. 2I was generated from a randomly connected recurrent network of linear rate neurons. The entries in the synaptic weight matrix are normally distributed with *J*_*ij*_ ∼ 𝒩 (0, g^2^/*N*), with a coupling strength *g* = 0.95 (31, 32). For consistency, we used the same number of time frames *T* = 7, 200 when comparing CI across all the datasets (Fig. 4BC, Fig. 5DE, Fig. S6C). For other cases, we analyzed the full length of the data (number of time frames: fish 1 - 7495, fish 2 - 9774, fish 3 - 13904, fish 4 - 7318, fish 5 - 7200, fish 6 - 9388). Next, the covariance matrix was calculated as 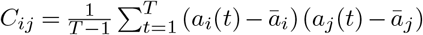, where 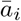 is the mean of *a*_*i*_*(t)* over time. Finally, to visualize covariance matrices on a common scale, we multiplied matrix C by a constant such that the average of its diagonal entries equaled one, that is, Tr(C)/*N* = 1. This scaling did not alter the shape of covariance eigenvalue distribution, but set the mean at 1 (see also Eq. (8)).

To maintain consistency across data sets, we fixed the same initial number of neurons at *N*_0_ = 1, 024. These *N*_0_ neurons were randomly chosen once for each zebrafish dataset and then used throughout the subsequent analyses. We adopted this setting for all analyses except in two particular instances: (1) for comparisons among the three sampling methods (RSap, ASap, and FSap), we specifically chose 1,024 neurons centered along the anterior-posterior axis, mainly from the midbrain to the anterior hindbrain regions (Fig. 5DE, Fig. S20). (2) When investigating the impact of hunting behavior on scale invariance, we included the entire neuronal population (section 4.11).

We used an iterative procedure to sample the covariance matrix *C* (calculated from data or as simulated ERMs). For RSap, in the first iteration, we randomly selected half of the neurons. The covariance matrix for these selected neurons was a *N*/2 × *N*/2 diagonal block of *C*. Similarly, the covariance matrix of the unselected neurons was another diagonal block of the same size. In the next iteration, we similarly created two new sampled blocks with half the number of neurons for each of the blocks we had. Repeating this process for n iterations resulted in 2^n^ blocks, each containing *N* := *N*_0_/2^n^ neurons. At each iteration, the eigenvalues of each block were calculated and averaged across the blocks after being sorted in descending order. Finally, the averaged eigenvalues were plotted against rank/*N* on a log-log scale.

In the case of ASap and FSap, the process of selecting neurons was different, although the remaining procedures followed the RSap protocol. In ASap, the selection of neurons was based on a spatial criterion: neurons close to the anterior end on the anterior-posterior axis were grouped to create a diagonal block of size 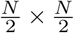 with the remaining neurons forming a separate block. FSap, on the other hand, used the Renormalization Group (RG) framework (20) to define the blocks (details in section 4.12). In each iteration, the cluster of neurons within a block that showed the highest average correlation 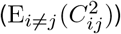 was identified and labeled as the most correlated cluster (refer to Fig. 5D, Figures S20 and S21).

In the ERM model, as part of implementing ASap, we generated anatomical and functional coordinates for neurons with a specified CCA properties as described in section 4.9. Mirroring the approach taken with our data, ASap segmented neurons into groups based on the first dimension of their anatomical coordinates, akin to the anterier-posterior axis. FSap employed the same RG procedures outlined earlier (section 4.12).

To determine the overall power-law coefficient of the eigenspectra, *α*, throughout sampling, we fitted a straight line in the log-log rank plot to the large eigenvalues that combined the original and three iterations of sampled covariance matrices (selecting the top 10% eigenvalues for each matrix and excluding the first four largest ones for each matrix). We averaged the estimated *α* over 10 repetitions of the entire sampling procedure. *R*^2^ of the power-law fit was computed in a similar way. To visualize the statistical structures of the original and sampled covariance matrices, the orders of the neurons (i.e. columns and rows) are determined by the following algorithm. We first construct a symmetric Toeplitz matrix 𝒯, with entries 𝒯_*i,j*_ = *t*_*i*−*j*_ and *t*_*i*−*j*_ ≡ *t*_*j*−*i*_. The vector 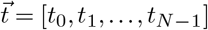 is equal to the mean covariance vector of each neuron calculated below. Let 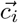 be a row vector of the data covariance matrix; we identify 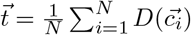, where *D*(·) denotes a numerical ordering operator, namely rearranging the elements in a vector 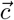 such that *c*_0_ ≥ *c*_1_ … ≥ *c*_N−1_. The second step is to find a permutation matrix *P* such that ║𝒯 − PCP^T^ ║_F_ is minimized, where ║ ║_F_ denotes the Frobenius norm. This quadratic assignment problem is solved by simulated annealing. Note that after sampling, the smaller matrix will appear different from the larger one. We need to perform the above reordering algorithm for every sampled matrix so that matrices of different sizes become similar in Fig. 2E.

The composite covariance matrix with substituted eigenvectors in (Fig. 2H) was created as described in the following steps. First, we generated a random orthogonal matrix *U*_*r*_ (based on the Haar measure) for the new eigenvectors. This was achieved by QR decomposition *A* = *U*_*r*_*R* of a random matrix *A* with i.i.d. entries *A*_*ij*_ ∼ 𝒩 (0, 1/*N*). The composite covariance matrix *C*_*r*_ was then defined as 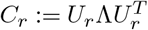, where Λ is a diagonal matrix that contains the eigenvalues of *C*. Note that since all the eigenvalues are real and *U*_*r*_ is orthogonal, the resulting *C*_*r*_ is a real and symmetric matrix. By construction, *C*_*r*_ and *C* have the same eigenvalues, but their *sampled* eigenspectra can differ.

### 4.4 Dimensionality

In this section, we introduce the Participation Ratio (*D*_PR_) as a metric for effective dimensionality of a system, based on (25–29, 62). *D*_PR_ is defined as:

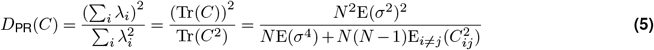

Here, *λ*_*i*_ are the eigenvalues of the covariance matrix *C*, representing variances of neural activities. Tr(*·*) denotes the trace of the matrix. The term 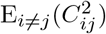 denotes the expected value of the squared elements that lie off the main diagonal of *C*. This represents the average squared covariance between the activities of distinct pairs of neurons.

With these definitions, we explore the asymptotic behavior of *D*_PR_ as the number of neurons *N* approaches infinity:

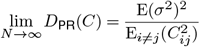

This limit highlights the relationship between the PR dimension and the average squared covariance among different pairs of neurons. To predict how *D*_PR_ scales with the number of neurons (Fig. 2D), we first estimated these statistical quantities 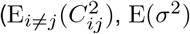, and E(*σ*^4^)) using all available neurons, then applied Eq. (5) for different values of *N*. It is worth mentioning that a similar theoretical finding is established by Dahmen et. al. (29). The transition from increasing *D*_PR_ with *N* to approaching the saturation point occurs when *N* is significantly larger than *D*_PR_.

### 4.5 ERM model

We consider the eigenvalue distribution or spectrum of the matrix *C* at the limit of *N* ≫ 1 and *L* ≫ 1. This spectrum can be analytically calculated in both high-density and intermediate-density scenarios using the replica method (34). The following sketch shows our approach, and detailed derivations can be found in Supp. Note. To calculate the probability density function of the eigenvalues (or eigendensity), we first compute the resolvent or Stieltjes transform 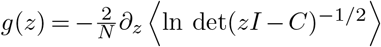. Here ⟨…⟩ is the average across the realizations of *C* (that is, random 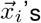 and 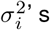). The relationship between the resolvent and the eigendensity is given by the Sokhotski-Plemelj formula:

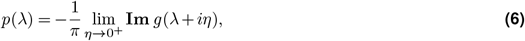

where Im means imaginary part.

Here we follow the field-theoretic approach (34), which turns the problem of calculating the resolvent to a calculation of the partition function in statistical physics by using the replica method. In the limit *N* → ∞, *L*^*d*^ → ∞, *ρ* being finite, by performing a leading order expansion of the canonical partition function at large *z* (Supp. Note), we find the resolvent is given by

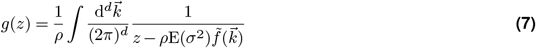

In the *high-density* regime, the probability density function (pdf) of the covariance eigenvalues can be approximated and expressed from Equations (6) and (7) using the Fourier transform of the kernel function 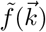:

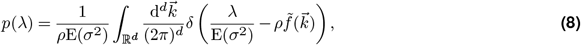

where *δ*(*x*) is the Dirac delta function and E(*σ*^2^) is the expected value of the variances of neural activity. Intuitively, Eq. (8) means that *λ/ρ* are distributed with a density proportional to the area of 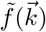’ level sets (i.e., isosurfaces).

In section 2.3, we found that the covariance matrix consistently shows greater scale invariance compared to the correlation matrix across all datasets. This suggests that the variability in neuronal activity significantly influences the eigenspectrum. This finding, however, cannot be explained by the high-density theory, which predicts that the eigenspectrum of the covariance matrix is simply a rescaling of the correlation eigenspectrum by 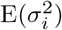, the expected value of the variances of neural activity. Without loss of generality, we can always standardize the fluctuation level of neural activity by setting E(*σ*^2^) = 1. This is equivalent to multiplying the covariance matrix *C* by a constant such that Tr(*C*)*/N* = 1, which in turn scales all the eigenvalues of *C* by the same factor. Consequently, the heterogeneity of 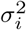 has no effect on the scale invariance of the eigenspectrum (see Eq. (8)). This theoretical prediction is indeed correct and is confirmed by direct numerical simulations and quantifying the scale invariance using the CI (Fig. S6A).

Fortunately, the inconsistency between theory and experimental results can be resolved by focusing the ERM within the intermediate density regime *ρϵ*^*d*^ ≪ 1, where neurons are positioned at a moderate distance from each other. As mentioned above, we set E(*σ*^2^) = 1 in our model and vary the diversity of activity fluctuations among neurons represented by E(*σ*^4^). Consistent with the experimental observations, we find that the CI decreases with E(*σ*^4^) (see Fig. S6B). This agreement indicates that the neural data are better explained by the ERM in the intermediate density regime.

To gain a deeper understanding of this behavior, we use the Gaussian variational method (34) to calculate the eigenspectrum. Unlike the high-density theory where the eigendensity has an explicit expression, in the intermediate density the resolvent *g*(*z*) no longer has an explicit expression and is given by the following equation

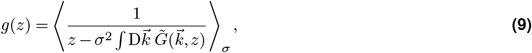

where ⟨…⟩_*σ*_ computes the expectation value of the term within the bracket with respect to *σ*, namely ⟨…⟩_*σ*_ ≡…*p*(*σ*)d*σ*. Here and in the following, we denote 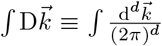. The function 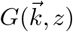 is determined by a self-consistent equation,

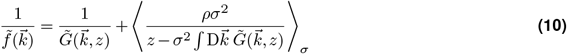

We can solve 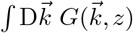 from Eq. (10) numerically and below is an outline, and the details are explained in Supp. Note. Let us define the integral 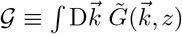 First, we substitute *z ≡ λ* + *iη* into Eq. (10) and write 𝒢 = **Re**𝒢 +*i***Im**𝒢. Eq. (10) can thus be decomposed into its real part and imaginary part, and a set of nonlinear and integral equations, each of which involves both **Re***𝒢* and **Im***𝒢*. We solve these equations at the limit *η* → 0 using a fixed-point iteration that alternates between updating **Re***𝒢* and **Im***𝒢* until convergence.

We find that the variational approximations exhibit excellent agreement with the numerical simulation for both large and intermediate *ρ* where the high-density theory starts to deviate significantly (for *ρ* = 256 and *ρ* = 10.24, *ϵ* = 0.03125, Fig. S3). Note that the departure of the leading eigenvalues in these plots is expected, since the power-law kernel function we use is not integrable (see section 4.6).

To elucidate the connection between the two different methods, we estimate the condition when the result of the high-density theory (Eq. (8)) matches that of the variational method (Equations (9) and (10)) (Supp. Note). The transition between these two density regimes can also be understood (see section 4.8.1 and Supp. Note).

Importantly, the scale invariance of the spectrum at *µ/d* → 0 previously derived using the high-density result (Eq. (3)) can be extended to the intermediate-density regime by proving the *ρ*-independence using the variational method (Supp. Note).

Finally, using the variational method and the integration limit estimated by simulation (see section 4.7.2), we show that the heterogeneity of the variance of neural activity, quantified by E(*σ*^4^), indeed improves the collapse of the eigenspectra for intermediate *ρ* (Supp. Note). Our theoretical results agree excellently with the ERM simulation (Fig. S6A, B).

### 4.6 Kernel function

Throughout the paper, we have mainly considered a particular approximate power-law kernel function inspired by the Student’s t distribution (section 2.2)

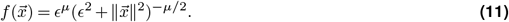

To understand how to choose *ϵ* and *µ*, see section 4.8.1. Variations of Eq. (11) near *x* = 0 have also been explored; see a summary in table S3.

**Table S3.** Modifications of the shape of 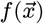 near 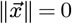 used in Fig. S7, Fig. S8 and Fig. S9. Flat: when 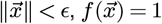,. Tangent: when 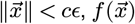 follows a tangent line of the exact power law (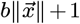 and 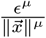 have a same first-order derivative when 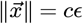). b and c are constants. Tent: when 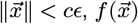 follows a straight line while the slope is not the same as the tangent case. Parabola: when 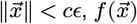 follows a quadratic function (ax^2^ + 1 and 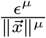 have same first-order derivative). t pdf: mimic the smoothing treatment like the t distribution. All the constant parameters are set such that f (0) = 1.

| $f(\vec{x})$ | Definition |
| --- | --- |
| Flat | $f(\vec{x}) = \begin{cases} 1, & \ \vec{x}\ < \epsilon \\ \frac{\epsilon^\mu}{\ \vec{x}\ ^\mu}, & \ \vec{x}\ \geq \epsilon \end{cases}$ |
| Tangent | $f(\vec{x}) = \begin{cases} b\ \vec{x}\ + 1, & \ \vec{x}\ < c\epsilon, f'(c\epsilon) = b \\ \frac{\epsilon^\mu}{\ \vec{x}\ ^\mu}, & \ \vec{x}\ \geq c\epsilon \end{cases}$ |
| Tent | $f(\vec{x}) = \begin{cases} b\ \vec{x}\ + 1, & \ \vec{x}\ < c\epsilon, f'(c\epsilon) \neq b \\ \frac{\epsilon^\mu}{\ \vec{x}\ ^\mu}, & \ \vec{x}\ \geq c\epsilon \end{cases}$ |
| Parabola | $f(\vec{x}) = \begin{cases} b\ \vec{x}\ ^2 + 1, & \ \vec{x}\ < c\epsilon, f'(c\epsilon) = 2bc\epsilon \\ \frac{\epsilon^\mu}{\ \vec{x}\ ^\mu}, & \ \vec{x}\ \geq c\epsilon \end{cases}$ |
| t pdf | $f(\vec{x}) = \epsilon^\mu (\epsilon^2 + \ \vec{x}\ ^2)^{-\mu/2}$ |

It is worth mentioning that a power law is not the only slow decaying function that can produce a scale-invariant covariance spectrum (Fig. S5). We choose it for its analytical tractability in calculating the eigenspectrum. Importantly, we find numerically that the two contributing factors to scale invariance – namely, slow spatial decay and higher functional space – can be generalized to other *nonpower-law* functions. An example is the stretched exponential function 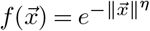 with 0 *< η <* 1. When *η* is small and *d* is large, the covariance eigenspectra also display a similar collapse upon random sampling (Fig. S5).

This approximate power-law 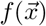 has the advantage of having an analytical expression for its Fourier transform, which is crucial for the high-density theory (Eq. (8)),

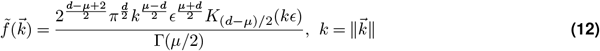

Here *K*_*α*_(*x*) is the modified Bessel function of the second kind, and Γ(*x*) is the Gamma function. We calculated the above formulas analytically for *d* = 1, 2, 3 with the assistance of Mathematica and conjectured the case for general dimension *d*, which we confirmed numerically for *d* ≤ 10.

We want to explain two technical points relevant to the interpretation of our numerical results and the choice of 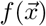. Unlike the case in the usual ERM, here we allow 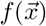 to be non-integrable (over ℝ^*d*^), which is crucial to allow power law 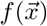. The nonintegrability violates a condition in the classical convergence results of the ERM spectrum (63) as *N → ∞*. We believe that this is exactly the reason for the departure of the first few eigenvalues from our theoretical spectrum (e.g., in Fig. 3). Our hypothesis is also supported by ERM simulations with integrable 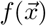 (Fig. S4), where the numerical eigenspectrum matches closely with our theoretical one, including the leading eigenvalues. For ERM to be a legitimate model for covariance matrices, we need to ensure that the resulting matrix *C* is positive semidefinite. According to the Bochner theorem (64), this is equivalent to the Fourier transform (FT) of the kernel function 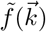 being nonnegative for all frequencies. For example, in 1D, a rectangle function 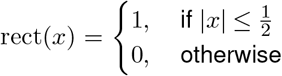 does not meet the condition (its FT is 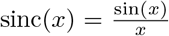, but a tent function tent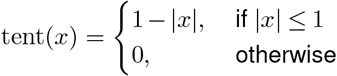 otherwise does (its FT is sinc^2^(*x*)). For the particular kernel function 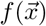 in Eq. (11), this condition can be easily verified using the analytical expressions of its Fourier transform (Eq. (12)). The integral expression for *K*_*α*_(*x*), given as 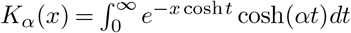 shows that *K*_*α*_(*x*) is positive for all *x >* 0. Likewise, the Gamma function Γ(*x*) *>* 0. Therefore, the Fourier transform of Eq. (11) is positive and the resulting matrix *C* (of any size and values of 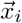) is guaranteed to be positive definite.

Building upon the theory outlined above, numerical simulations further validated the empirical robustness of our ERM model, as showcased in Fig. 3B-D and Fig. 4A. In Fig. 3B-D, the ERM was characterized by the parameters *N* = 1024, *d* = 2, *L* = 10, *ρ* = 10.24 and *µ* = 0.5 and *ϵ* = 0.03125 for 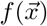. To numerically compute the eigenvalue probability density function, we generated the ERM 100 times, each sampled using the method described in section 4.3. The probability density function (pdf) was computed by calculating the pdf of each ERM realization and averaging these across the instances. The curves in Fig. 3D showed the average of over 100 ERM simulations. The shaded area (most of which is smaller than the marker size) represented the SEM. For Fig. 4A, the columns from left to right were corresponded to *µ* = 0.5, 0.9, 1.3, and the rows from top to bottom were corresponded to *d* = 1, 2, 3. Other ERM simulation parameters: *N* = 4096, *ρ* = 256, *L* = (*N/ρ*)^1*/d*^, *ϵ* = 0.03125 and 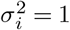. It should be noted that for Fig. 4A, the presented data pertain to a single ERM realization.

### 4.7 Collapse index (CI)

We quantify the extent of scale invariance using CI defined as the area between two spectrum curves (Fig. 4A upper right), providing an intuitive measure of the shift of the eigenspectrum when varying the number of sampled neurons. We chose the CI over other measures of distance between distributions for several reasons. First, it directly quantifies the shift of the eigenspectrum, providing a clear and interpretable measure of scale invariance. Second, unlike methods that rely on estimating the full distribution, the CI avoids potential inaccuracies in estimating the probability of the top leading eigenvalues. Finally, the use of CI is motivated by theoretical considerations, namely the ERM in the high-density regime, which provides an analytical expression for the covariance spectrum (Eq. (3)) valid for large eigenvalues.

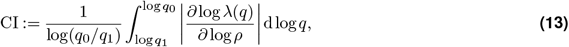

we set *q*_1_ such that *λ*(*q*_1_) = 1, which is the mean of the eigenvalues of a normalized covariance matrix. The other integration limit *q*_0_ is set to 0.01 such that *λ*(*q*_0_) is the 1% largest eigenvalue.

Here we provide numerical details on calculating CI for the ERM simulations and experimental data.

#### 4.7.1 A calculation of collapse index for experimental datasets/ERM model

To calculate CI for a covariance matrix *C* of size *N*_0_, we first computed its eigenvalues 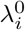 and those of the sampled block *C*_*s*_ of size *N*_*s*_ = *N*_0_*/*2, denoted as 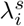 (averaged over 20 times for the ERM simulation and 2000 times in experimental data). Next, we estimated log*λ*(*q*) using the eigenvalues of *C*_0_ and *C*_*s*_ at *q* = *i/N*_*s*_, *i* = 1, 2,…, *N*_*s*_. For the sampled *C*_*s*_, we simply had 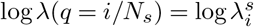, its *i*-th largest eigenvalue. For the original *C*_0_, log*λ*(*q* = *i/N*_*s*_) was estimated by a linear interpolation, *on the* log*λ-*log*q scale*, using the value of log*λ*(*q*) in the nearest neighboring *q* = *i/N*_0_’s (which again are simply 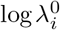). Finally, the integral (Eq. (13)) was computed using the trapezoidal rule, discretized at *q* = *i/N*_*s*_’ s, using the finite difference 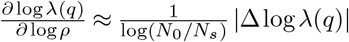, where Δ denotes the difference between the original eigenvalues of *C*_0_ and those of sampled *C*_*s*_.

#### 4.7.2 Estimating CI using the variational method

In the definition of CI (Eq. (13), calculating *λ*(*q*) and 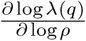 directly using the variational method is difficult, but we can make use of an implicit differentiation

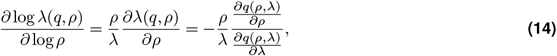

where 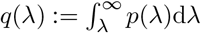 is the complementary cdf (the inverse function of *λ*(*q*) in section 4.7.1). Using this, the integral in CI (Eq. (13)) can be rewritten as

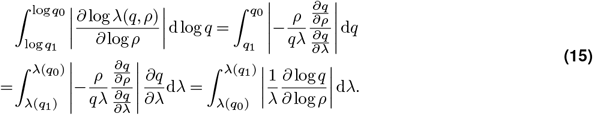

Since 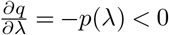, we switch the order of the integration interval in the final expression of Eq. (15).

First, we explain how to compute the complementary cdf *q*(*λ*) numerically using the variational method. The key is to integrate the probability density function *p*(*λ*) from *λ* to a finite *λ*(*q*_*s*_) rather than to infinity,

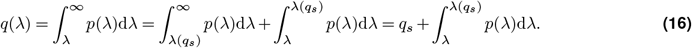

The integration limit *λ*(*q*_*s*_) cannot be calculated directly using the variational method. We thus used the value of *λ*^*s*^(*q*_*s*_ ≈ *q*_0_) (section 4.7) from simulations of the ERM with a large *N* = 1024 as an approximation. Furthermore, we employed a smoothing technique to reduce bias in the estimation of *λ*^*s*^(*q*_*s*_) due to the leading zigzag eigenvalues (i.e., the largest eigenvalues) of the eigenspectrum. Specifically, we determined the nearest rank *j* < *N*_*q*0_ and then smoothed the eigenvalue log λ^*s*^(*q*_*s*_) on the log-log scale using the formula 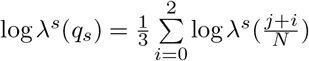, and 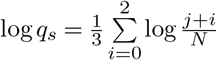 averaging over 100 ERM simulations.

Note that we can alternatively use the high-density theory (Supp. Note) to compute the integration limit *λ*(*q*_*s*_ = 1*/N*) instead of resorting to simulations. However, since the true value deviates from the *λ*^*h*^(*q*_*s*_ = 1*/N*) derived from high-density theory, this approach introduces a constant bias (Fig. S6) when computing the integral in Eq. (16). Therefore we used the simulation value *λ*^*s*^(*q*_*s*_ ≈ *q*_0_) when producing Fig. S6AB.

Next, we describe how each term within the integral of Eq. (15) was numerically estimated. First, we calculated 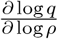 with a similar method described in section 4.7.1. Briefly, we calculated *q*_0_(*λ*) for density 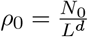 and *q*_*s*_(*λ*) for density 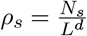 and then used the finite difference 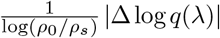. Second, 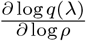 was evaluated at 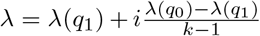, where *i* = 0, 1, 2,…, *k* − 1, and we used *k* = 20. Finally, we performed a cubic spline interpolation of the term 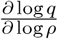 and obtained the theoretical CI by an integration of Eq. (15). Fig. S6A,B shows a comparison between theoretical CI and that obtained by numerical simulations of ERM (section 4.7.1).

### 4.8 Fitting ERM to data

#### 4.8.1 Estimating the ERM parameters

Our ERM model has 4 parameters: *µ* and *ϵ* dictate the kernel function 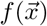, whereas the box size *L* and the embedding dimension *d* determine the neuronal density *ρ*. In the following, we describe an approximate method to estimate these parameters from pairwise correlations measured experimentally 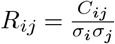. We proceed by deriving a relationship between the correlation probability density distribution *h*(*R*) and the pairwise distance probability density distribution 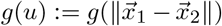 in the functional space, from which the parameters of the ERM can be estimated.

Consider a distribution of neurons in the functional space with a coordinate distribution 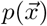. The pairwise distance density function *g*(*u*) is related to the spatial point density by the following formula:

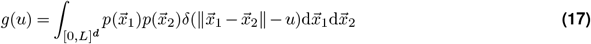

For ease of notation, we subsequently omit the region of integration, which is the same as here. In the case of a uniform distribution, 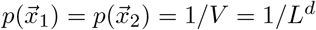. For other spatial distributions, Eq. (17) cannot be explicitly evaluated. We therefore make a similar approximation by focusing on a small pairwise distance (i.e., large correlation):

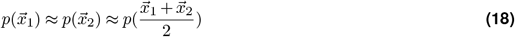

By a change of variables:

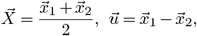

Eq. (17) can be rewritten as

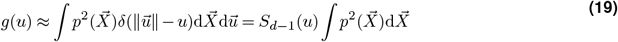

where *S*_*d*−1_(*u*) is the surface area of *d* − 1 sphere with radius *u*. Note that the approximation of *g*(*u*) is not normalized to 1, as Eq. (19) provides an approximation valid only for small pairwise distances (i.e., large correlation). Therefore, we believe this does not pose an issue.

With the approximate power-law kernel function 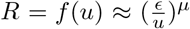 the probability density function of pairwise correlation *h*(*R*) is given by:

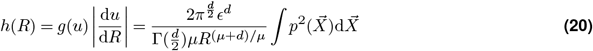

Taking the logarithm on both sides

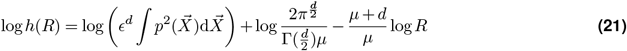

Eq. (21) is the key formula for ERM parameters estimation. In the case of a uniform spatial distribution, 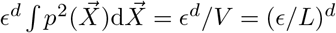For a given dimension *d*, we can therefore estimate *µ* and (*ϵ/L*)^*d*^ separately by fitting *h*(*R*) on the log-log scale using the linear least squares. Lastly, we fit the distribution of *σ*^2^ (the diagonal entries of the covariance matrix *C*) to a log-normal distribution by estimating the maximum likelihood.

There is a redundancy between the unit of the functional space (using a rescaled *ϵ*_*δ*_ ≡ *ϵ/δ*) and the unit of 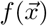 (using a rescaled 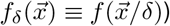, thus *ϵ* and *L* are a pair of redundant parameters: once *ϵ* is given, *L* is also determined. We set *ϵ* = 0.03125 throughout the article. In summary, for a given dimension *d* and *ϵ, µ* of 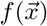 (Eq. (11)), the distribution of *σ*^2^ (section 2.2) and *ρ* (or equivalently *L*) (section 2.2) can be fitted by comparing the distribution of pairwise correlations in experimental data and ERM. Furthermore, knowing (*ϵ/L*)^*d*^ enables us to determine *a fundamental dimensionless parameter*

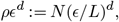

which tells us whether the experimental data are better described by the high-density theory or the Gaussian variational method (Supp. Note). Indeed, the fitted *ρϵ*^*d*^ ∽ 10^−3^ − 10^0^ is much smaller than 1, consistent with our earlier conclusion that neural data are better described by an ERM model in the intermediate-density regime.

Notably, we found that a smaller embedding dimension *d* ≤ 5 gave a better fit to the overall pairwise correlation distribution. The following is an empirical explanation. As *d* grows, to best fit the slope of log*h*(*R*) − log *R, µ* will also grow. However, for very high dimensions *d*, the y-intercept would become very negative, or equivalently, the fitted correlation would become extremely small. This can be verified by examining the leading order log *R* independent term in Eq. (21) which can be approximated as 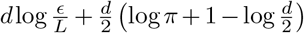. It becomes very negative for larg *d* since *ϵ* ≪*L* by construction. Throughout this article, we use *d* = 2 when fitting the experimental data with our ERM model.

The above calculation can be extended to the cases where the coordinate distribution 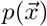 becomes dependent on other parameters. To estimate the parameters in coordinate distributions that can generate ERMs with a similar pairwise correlation distribution (Fig. S9), we fixed the integral value 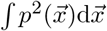. Consider, for example, a transformation of the uniform coordinate distribution to the normal distribution 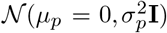 in ℝ^2^. We imposed 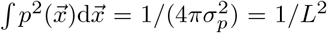. For the log-normal distribution, a similar calculation led to 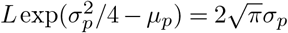 The numerical values for these parameters are shown in section 4.10. However, note that due to the approximation we used (Eq. (18)), our estimate of the ERM parameters becomes less accurate if the density function 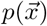 changes rapidly over a short distance in the functional space. More sophisticated methods, such as grid search, may be needed to tackle such a scenario.

After determining the parameters of the ERM, we first examine the spectrum of the ERM with uniformly distributed random functional coordinates 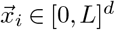 (Fig. S10M-R). Second, we use 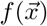 to translate experimental pairwise correlations into pairwise distances for all neurons in the functional space (Fig. S11, Fig. S10G-L). The embedding coordinates 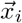 in the functional space can then be solved through Multidimensional Scaling (MDS) by minimizing the Sammon error (section 4.8.3). The similarity between the spectra of the uniformly distributed coordinates (Fig. S10M-R) and those of the embedding coordinates (Fig. S10G-L) is also consistent with the notion that specific coordinate distributions in the functional space have little impact on the shape of the eigenspectrum (Fig. S9).

#### 4.8.2 Nonnegativity of data covariance

To use ERM to model the covariance matrix, the pairwise correlation is given by a *non-negative* kernel function 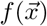 that monotonically decreases with the distance between neurons in the functional space. This nonnegativeness brings about a potential issue when applied to experimental data, where, in fact, a small fraction of pairwise correlations/covariances are negative. We have verified that the spectrum of the data covariance matrix (Fig. S18) remains virtually unchanged when replacing these negative covariances with zero (Fig. S18). This confirms that the ERM remains a good model when the neural dynamics is in a regime where pairwise covariances are mostly positive (51) (see also Fig. S2B, Fig. S2B-D).

#### 4.8.3 Multidimensional Scaling (MDS)

With the estimated ERM parameters (*µ* in 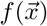 and the box size *L* for given *ϵ* and *d*, see section 4.8.1), we performed MDS to infer neuronal coordinates 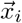 in functional space. First, we computed a pairwise correlation 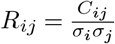 from the data covariances. Next, we calculated the pairwise distance, denoted by 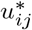 by computing the inverse function of 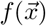with respect to the absolute value 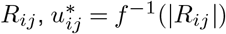 We used the absolute value | *R*_*ij*_ | instead of *R*_*ij*_ as a small percentage of *R*_*ij*_ are negative (Fig. S2A-D) where the distance is undefined. This substitution by the absolute value serves as a simple workaround for the issue and is only used here in the analysis to infer the neuronal coordinates by MDS. Finally, we estimated the embedding coordinates 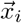 for each neuron by the SMACOF algorithm (Scaling by MAjorizing a COmplicated Function), which minimizes the Sammon error

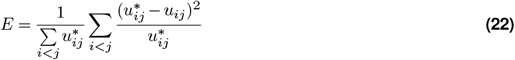

where 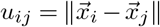 is the pairwise distance in the embedding space calculated above.

To reduce errors at large distances (i.e., small correlations with *R*_*ij*_ *< f*(*L*), where *L* is the estimated box size), we performed a soft cut-off at a large distance:

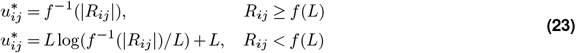

During the optimization process, we started at the embedding coordinates estimated by the classical MDS (46), with an initial sum of squares distance error that can be calculated directly, and ended with an error or its gradient smaller than 10^−4^.

The fitted ERM with the embedding coordinates 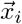 reproduced the experimental covariance matrix including the cluster structures (Fig. S11) and its sampling eigenspectra (Fig. S10).

### 4.9 Canonical-Correlation Analysis (CCA)

Here we briefly explain the CCA method (65) for completeness. The basis vectors 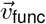 and 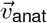, in functional and anatomical space, respectively, were found by maximizing the correlation 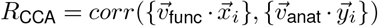. These basis vectors satisfy the condition that the projections of the neuron coordinates along them, 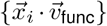 and 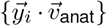, are maximally correlated among all possible choices of 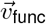 and 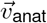. Here 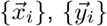 represent the coordinates in functional and anatomical spaces, respectively. The resulting maximum correlation is *R*_CCA_. To check the significance of the canonical correlation, we shuffled the functional space coordinates 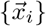 across neurons’ identity and re-calculated the canonical correlation with the anatomical coordinates, as shown in Fig. S13.

To study the effect of functional-anatomical relation described by *R*_CCA_ in the ERM model, we generated three dimensional anatomical coordinates 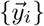 and two dimensional functional coordinates 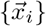 for each neuron which are jointly five-dimensional zero-mean multivariate Gaussian random variables. The coordinates are independent among each other, except for the first dimension 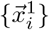 of the functional coordinates and the first dimension 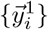 which are assigned to have a correlation coefficient equals to *R*_CCA_. The variances of the coordinates are 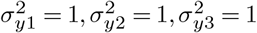 and 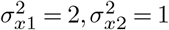 for the numerics in Fig. S21. Under this construction, the first canonical correlation between the anatomical and functional coordinates equals *R*_CCA_, and the first canonical direction 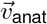 in the anatomical space is (1, 0, 0)^*T*^ and the first canonical direction 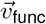 in the functional space is (1, 0)^*T*^.

### 4.10 Extensions of ERM and factors not affecting the scale invariance

In Fig. S9 we considered five additional types of spatial density distributions (coordinate distributions) in functional space and two additional functional space geometries. We examined the points distributed according to the uniform distribution 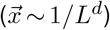 the normal distribution 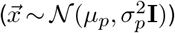 and the log-normal distribution 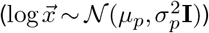. We used the method described in Methods section 4.8.1 to adjust the parameters of the coordinate distributions based on the uniform distribution case, so that they all generate similar pairwise correlation distributions. The relationships between these parameters are described in Methods section 4.8.1. In Fig. S9B, we used the following parameters: *d* = 2; *L* = 10 for the uniform distribution; *µ*_*p*_ = 0, *σ*_*p*_ = 2.82 for the normal distribution; and *µ*_*p*_ = 2, *σ*_*p*_ = 0.39 for the log-normal distribution.

Second, we introduced multiple clusters of neurons in the functional space, with each cluster uniformly distributed in a box. We considered three arrangements: (1) two closely situated clusters (with a box size of 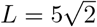the distance between two cluster centers being *L*_*c*_ = *L*), (2) two distantly situated clusters (with a box size of 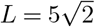 and the distance between clusters *L*_*c*_ = 4*L*), and three clusters arranged symmetrically in an equilateral triangle (with a box size of 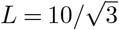 and the distance between clusters *L*_*c*_ = *L*).

Finally, we examined the scenario in which the points were uniformly distributed on the surface of a sphere (4*πl*^2^ = *L*^2^, *l* being the radius of the sphere) or a hemisphere (2*πl*^2^ = *L*^2^) embedded in ℝ ^3^ (the pairwise distance is that in ℝ ^3^). It should be noted that both cases have the same surface area as the 2D box.

### 4.11 Analyzing the effects of removing neural activity data during hunting

To identify and remove the time frames corresponding to putative hunting behaviors, the following procedure was used. The hunting interval was defined as 10 frames (1 sec) preceding the onset of an eye convergence (see Methods section 4.1.1) to 10 frames after the offset of this eye convergence. These frames were then excluded from the data before recalculating the covariance matrix (see Methods section 4.3) and subsequently the sampled eigenspectra (Fig. S15B, Fig. S16B,D,F,H). As a control to the removal of the hunting frame, an equal number of time frames that are not within those hunting intervals were randomly selected and then removed and analyzed (Fig. S15C, Fig. S16A,C,E,G). The number of hunting interval frames and total recording frames for five fish exhibiting hunting behaviors are as follows: fish 1 - 268/7495, fish 2 - 565/9774, fish 3 - 2734/13904, fish 4 - 843/7318 and fish 5 - 1066/7200. Fish 6 (number of time frames: 9388) was not exposed to a prey stimulus and, therefore, was excluded from the analysis.

To assess the impact of hunting removal on CI, we calculated the CI of the covariance matrix using all neurons recorded in each fish (without sampling to 1024 neurons). For the control case, we repeated the removal of the nonhunting frame 10 times to generate 10 covariance matrices and computed their CIs. We used a one-sample t-test to determine the level of statistical significance between the control CIs and the CI obtained after removal of the hunting frame.

Using fitted ERM parameters by full data, we performed a MDS on the control data and hunting-removed data to infer the functional coordinates. Note that the functional coordinates inferred by MDS are not unique: rotations and translations give equivalent solutions. For visualization purposes (not needed for analysis), we first used the Umeyama algorithm to optimally align the functional coordinates of control and hunting-removed data.

To identify distinct clusters within the functional coordinates, we fit Gaussian Mixture Models (GMMs) using the “GaussianMixtures” package in Julia. We chose the number of clusters *K* based on giving the smallest Bayesian Information Criterion (BIC) score. After fitting the GMMs, a list of probabilities *p*_*ik*_, *k* = 1, 2,…, *K* was given for each neuron *i* specifying the probability of the neuron belonging to the cluster *k*. The mean and covariance parameters were estimated for each Gaussian distributed cluster. For visualization (but not for analysis), a neuron was colored according to cluster *k*^*^ where *k*^*^ = arg max_1≤*k*≤*K*_ *p*_*ik*_.

We used the following method to measure the size of the cluster and its fold change. For a 2D (recall *d* = 2 in our ERM) Gaussian distributed cluster, let us consider an ellipse centered on its mean, and its axes are aligned with the eigenvectors of its covariance matrix *C*_2*×*2_. Let the eigenvalues of *C* be *λ*_1_, *λ*_2_. Then we set the length of the half-axis of the ellipse to be 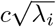, respectively. Here *c >* 0 is a constant determined below. Note that the ellipse axes correspond to linear combinations of 2D Gaussian random variables that are independent and *λ*_*i*_’s are the variance of these linear combinations. From this fact, it is straightforward to show that the probability that a sample from the Gaussian cluster lies in the above ellipse depends only on *c*, that is, 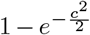, and not on the shape of the cluster. So, the ellipse represents a region that covers a fixed proportion of neurons for any cluster, and its area can be used as a measure for the size of the Gaussian cluster. Note that the area of the ellipse is 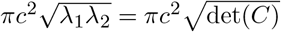 In Fig. S17, we plot the ellipses to help visualize the clusters and their changes. We choose *c* such that the ellipse covers 95% of the probability (that is, the fraction of neurons belonging to the cluster).

In the control functional map where we fit the GMMs, we directly calculated the size measure 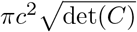 from the estimated covariance *C* for each Gaussian cluster. In the hunting-removed functional map, we needed to estimate the covariance *C*^*′*^ for neurons belonging to a cluster *k* under the new coordinates (we assume that the new distribution can still be approximated by a Gaussian distribution). We performed this estimation in a probabilistic manner to avoid issues of highly overlapping clusters where the cluster membership could be ambiguous for some neurons. First, we estimated the center/mean of the new Gaussian distribution by

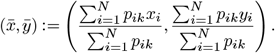

Here the summation goes over all the *N* neurons in the functional space and *p*_*ik*_ is the membership probability defined above, and (*x*_*i*_, *y*_*i*_) is the coordinate of neuron *i* in the hunting-removed map. Similarly, we can use a weighted average to estimate the entries in the covariance matrix 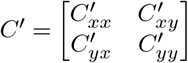. For example,

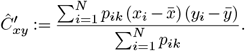

Then we calculated the size of the cluster on the new map as 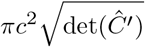. Finally, we computed the fold change in size as 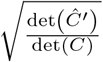.

### 4.12 Renormalization-Group (RG) Approach

Here we briefly summarize the RG approach used in (20) and elucidate the adjustments required when applying the RG approach to ERM. The method consists of two stages: (i) iterative agglomerate clustering of neurons, and (ii) computing the spectrum of a block of the *original* covariance matrix corresponding to a cluster of the desired size based on the previous clustering result.

#### 4.12.1Stage (i): Iterative Clustering

We begin with *N*_0_ neurons, where *N*_0_ is assumed to be a power of 2. In the first iteration, we compute Pearson’s correlation coefficients for all neuron pairs. We then search greedily for the most correlated pairs and group the half pairs with the highest correlation into the first cluster; the remaining neurons form the second cluster. For each pair (*a, b*), we define a coarse-grained variable according to:

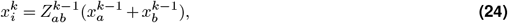

where 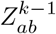 normalizes the average to ensure unit nonzero activity. This process reduces the number of neurons to *N*_1_ = *N*_0_*/*2. In subsequent iterations, we continue grouping the most correlated pairs of the coarse-grained neurons, iteratively reducing the number of neurons by half at each step. This process continues until the desired level of coarse-graining is achieved.

When applying the RG approach to ERM, instead of combining neural activity, we merge correlation matrices to traverse different scales. During the *k*th iteration, we compute the coarse-grained covariance as:

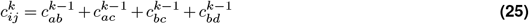

and the variance as:

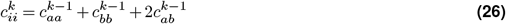

Following these calculations, we normalize the coarse-grained covariance matrix to ensure that all variances are equal to one. Note that these coarse-grained covariances are only used in stage (i) and not used to calculate the spectrum.

#### 4.12.2 Stage (ii): Eigenspectrum Calculation

The calculation of eigenspectra at different scales proceeds through three sequential steps. First, for each cluster identified in Stage (i), we compute the covariance matrix using the original firing rates of neurons within that cluster (not the coarse-grained activities). Second, we calculate the eigenspectrum for each cluster. Finally, we average these eigenspectra across all clusters at a given iteration level to obtain the representative eigenspectrum for that scale.

In stage (ii), we calculate the eigenspectra of the sub-covariance matrices across different cluster sizes as described in (20). Let *N*_0_ = 2^*n*^ be the original number of neurons. To reduce it to size *N* = *N*_0_*/*2^*k*^ = 2^*n*−*k*^, where *k* is the kth reduction step, consider the coarse-grained neurons in step *n − k* in stage (i). Each coarse-grained neuron is a cluster of 2^*n*−*k*^ neurons. We then calculate spectrum of the block of the original covariance matrix corresponding to neurons of each cluster (there are 2^*k*^ such blocks). Lastly, an average of these 2^*k*^ spectra is computed.

For example, when reducing from *N*_0_ = 2^3^ = 8 to *N* = 2^3−1^ = 4 neurons (*k* = 1), we would have two clusters of 4 neurons each. We calculate the eigenspectrum for each 4×4 block of the original covariance matrix, then average these two spectra together. To better understand this process through a concrete example, consider a hypothetical scenario where a set of eight neurons, labeled 1, 2, 3, …, 7, 8, are subjected to a two-step clustering procedure. In the first step, neurons are grouped based on their maximum correlation pairs, for example, resulting in the formation of four pairs: {1, 2}, {3, 4}, {5, 6}, and {7, 8} (see Fig. S22). Subsequently, the neurons are further grouped into two clusters based on the results of the RG step mentioned above. Specifically, if the correlation between the coarse-grained variables of the pair {1, 2} and the pair {3, 4} is found to be the largest among all other pairs of coarse-grained variables, the first group consists of neurons {1, 2, 3, 4}, while the second group contains neurons {5, 6, 7, 8}. Next, take the size of the cluster *N* = 4 for example. The eigenspectra of the covariance matrices of the four neurons within each cluster are computed. This results in two eigenspectra, one for each cluster. The correlation matrices used to compute the eigenspectra of different sizes do not involve coarse-grained neurons. It is the real neurons 1, 2, 3, …, 7, 8, but with expanding cluster sizes. Finally, the average of the eigenspectra of the two clusters is calculated.

### 4.13. Spectrum of three types of sampling procedures in ERM model

In section 2.4 we have considered three types of sampling procedures: random sampling (RSap), spatial sampling in the anatomical space (ASap, e.g., recording neurons in a brain region), and spatial sampling in the functional space (FSap), namely spatial sampling in functional space by subdividing the space into smaller regions, is equivalent to the previously reported renormalization group (RG) inspired process (66, 67). Here we consider the relationship between the spectrum of three types of sampling procedures.

We assume a uniform random distribution of neurons in a *d*-dimensional functional space, [0, *L*]^*d*^. For RSap procedures, the resulting neuronal density *ρ*_*R*_ is reduced to *ρ*_*R*_ = *kρ*_0_, with *k* representing the sampling ratio (*k* = *N/N*_0_) and *ρ*_0_ being the initial density. In contrast, FSap maintains the original density, *ρ*_*F*_ = *ρ*_0_. This constancy in neuronal density under FSap ensures that the covariance eigenspectrum remains invariant across scales for any spatial correlation functions 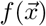, such as power law and exponential, as shown in Fig. S19A,B,D,E. In contrast, RSap reduces *ρ*, thus demanding more rigorous conditions to achieve a scale-invariant covariance spectrum (e.g., compare Fig. S19A and C).

Under ASap, sampled neurons are not spread out evenly in functional space, whereas our theoretical framework assumes a uniform distribution. To reconcile this discrepancy, we employ a uniform approximation of the neural distribution. This approach involves introducing an effective density, *ρ*^*′*^, defined as the spatial average of the density function 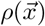. This adjustment allows our theoretical model to accommodate non-uniform distributions encountered in anatomically spatial sampling.

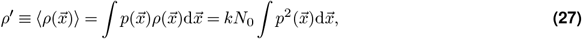

where 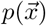 is the normalized density distribution (see Methods section 4.8.1). using the Cauchy-Schwarz inequality, we have

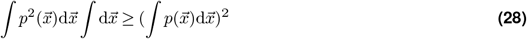

thus *ρ*^*′*^ ≥ *kρ*_0_.

According to the condition 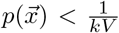, we have *ρ*^′^ *ρ*_0_, intuitively, sampling within a uniformly distributed neuron population does not increase the density.

So we have *ρ*_0_ ≥ *ρ*^*′*^_*A*_ ≥ *kρ*_0_, i.e., *ρ*_*F*_ ≥ *ρ*^*′*^_*A*_ ≥ *ρ*_*R*_. Thus the spectrum ASap should be between FSap and RSap.

### 4.14 Dimensions of three types of sampling procedures in ERM model

#### 4.14.1 Scaling of Dimensions through Random Samplings

Let us revisit the definition of the Participation Ratio (PR) dimension as defined in Equation Eq. (5):

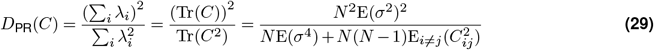

During the random sampling process, the expected values *E*(*σ*^2^), *E*(*σ*^4^), and 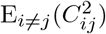 remain constant. These constants allow for the estimation of the PR dimension across various scales using:

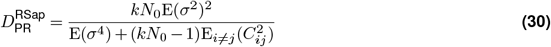

Here, *k* = *N/N*_0_ represents a scaling factor (fraction) associated with sampling. The key question is to understand how the dimensionality changes with *k*. Under random sampling, as *k* increases, the dimensionality will quickly approaches a saturating point defined by Eq. (1).

#### 4.14.2 Scaling of Dimensions through Functional Sampling

In this section, we leverage the uniform ERM model to estimate dimensions within the context of functional sampling, specifically focusing on the estimation of squared pairwise covariance 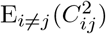 and dimensionality.

Adopting an approximation for a power-law kernel function *f*(*x*) ≈ *ϵ*^*µ*^∥*x*∥^−*µ*^ allows us to express the expected value of the squared covariance 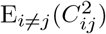 as follows:

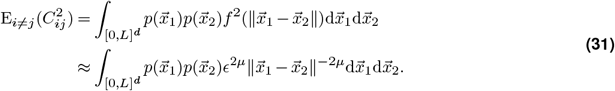

For a set subjected to functional sampling with a sampling fraction *k*, this procedure adjusts the size of the functional space in the ERM model by a factor of *k*^−1*/d*^. Consequently, the 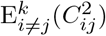 for the sampled fraction *k* is given by:

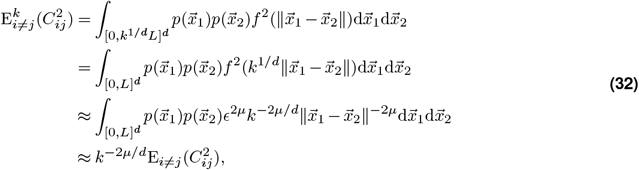

Here we assume that *E*[*σ*^2^] and *E*[*σ*^4^] are constant across the sampling process. This model enables the estimation of the ratio *µ/d* as detailed in the Methods section 4.8.1.

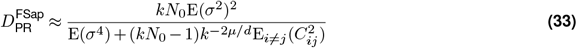

In the large *N* limit, we observe distinct behaviors in the evolution of dimensionality in both theory and data: it saturates in RSap (dashed line in Fig. 5D), namely 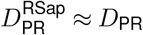 defined in Eq. (1), whereas it follows a different scaling relationship 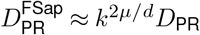 in FSap (solid line in Fig. 5D).

#### 4.14.3 Comparative Analysis of PR Dimension Across sampling Techniques

This section examines the behavior of the Participation Ratio (PR) dimension under three sampling techniques: anatomical sampling, random sampling, and functional sampling. We show that the average PR dimension following anatomical sampling occupies a middle ground between the extremes presented by random and functional sampling.

The PR dimension, denoted *D*_PR_, reflects the sampling impact and depends on the distribution 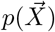 of the functional coordinates 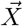. Defining the sampling fraction as *k* = 1*/q*, the mean *D*_PR_ is represented as:

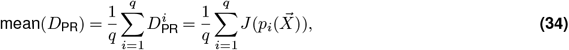

where the neuron set 1, 2, …, *N* is segmented into *q* clusters 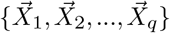, each comprising 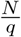 neurons. The probability distribution 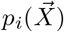 corresponds to each cluster 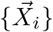. The probability distribution for each cluster, 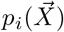, emerges naturally from the sampling process.

The equivalence of the mean probability density function across the sampled clusters to the original set’s probability density function leads us to the condition:

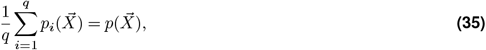

This condition is a direct consequence of the sampling process, ensuring that the aggregated probability density function of all sampled sets mirrors the overall density distribution of the neurons.

Applying the Lagrange multiplier method to optimize the mean *D*_PR_:

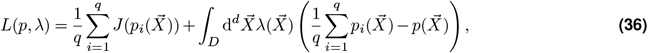

Here *L*(*p, λ*) is the Lagrangian, 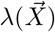 is the Lagrange multiplier, we derive the optimal condition:

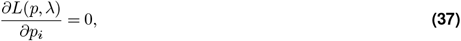

yielding:

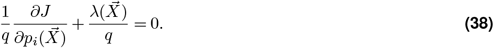

At the optimal mean *D*_PR_, each 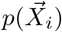 is equivalent, leading to 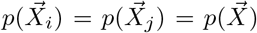 (representative of random sampling). Hence, the mean *D*_PR_ post-random sampling sets the upper limit for the mean *D*_PR_ after anatomical sampling.

Let us investigate the lower bound of the mean PR dimension with the ERM model. For the minimization of mean(*D*_PR_), a key requirement is the functional spatial proximity of neurons within the same cluster, in other words, the neuron set should be distinctly separated in functional space. Consequently, achieving the minimum mean PR dimension necessitates a functional sampling strategy.

#### 4.14.4 Derive upper bound of dimension from spectrum

To deduce *D*_*PR*_ from the spectrum, for simplicity, we focus on the high-density region, where we have an analytical expression for *λ* that is valid for large eigenvalues:

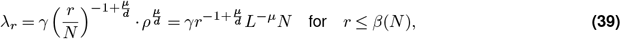

where *L* is the size of the functional space, *γ* is the coefficient in Eq. (3), which depends on *d, µ*, and E(*σ*^2^). Note that the eigenvalue *λ*_*r*_ decays rapidly after the threshold *r* = *β*(*N*). Since we did not discuss small eigenvalues in this article, we represent them here as an unknown function *η*(*r, N, L*):

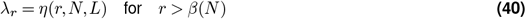

As discussed in section 4.5, without changing the properties of the spectrum, we can always impose E(*σ*^2^) = 1 such that

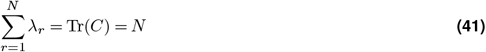

We emphasize that this constraint requires that large and small eigenvalues behave differently because otherwise 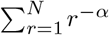 with *α <* 1 would scale as *N* ^1−*α*^, and 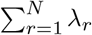_*r*_ is not proportional to *N*.

Using the Cauchy–Schwarz inequality, we have an upper bound of 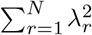:

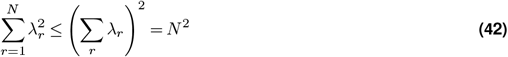

On the other hand, 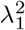 is a lower bound of 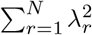:

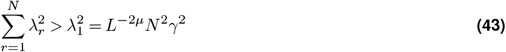

As a result, the dimensionality

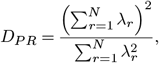

is bounded as

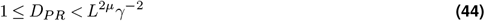

Under random sampling, *L* remains fixed. Thus, we must have a bounded dimensionality that is independent of *N* for our ERM model. A tighter lower bound of 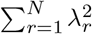 is

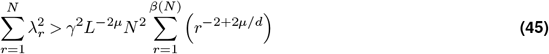

A tighter upper bound of participation ratio *D*_*P R*_ can be written as:

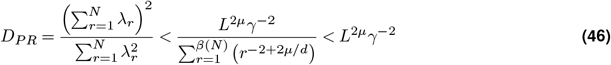

However, in functional sampling, enlarging the region size with constant density *ρ* results in *L ∼ N*^1*/d*^. Thus, the upper bound of *D*_*P R*_ should grow as *N* ^2*µ/d*^, consistent with the previously derived result (Eq. (33)) in section 4.14.2.

#### 4.14.5 Simulating CCA and anatomical sampling

In this section, we estimate the dimensions of the anatomically sampled neuron set. For simplicity, we assume that the functional coordinates of neurons, *X*_*i*_, and the anatomical coordinates of neurons, *Y*_*i*_, both follow a multivariate Gaussian distribution. We define anatomical sampling, which involves sampling on *Y*_*i*_, along a direction chosen arbitrarily and denote this direction as *Y* ^*A*^. Subsequently, we perform sampling on *X*_*i*_ in the direction denoted by *X*^*A*^, which is determined to have the highest correlation with *Y* ^*A*^ according to Canonical Correlation Analysis (CCA). This process effectively mimics the scenario of functional sampling.

The key to calculating the PR dimension involves computing the expected value 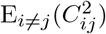. In the ERM model, the distribution of *C*_*ij*_ can be estimated by the distribution of points in the functional space. This allows for the calculation of the PR dimension across anatomical sampling by comparing the distribution of *X*_*i*_ after anatomical sampling with that after functional sampling. We can model the distribution of *X*^*A*^ and *Y* ^*A*^ as follows:

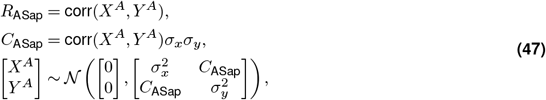

Here we consider only the projection of the functional coordinate onto the direction *X*^*A*^, which exhibits the highest correlation, denoted by *R*_ASap_, with *Y* ^*A*^. Specifically, when selecting the anatomical direction as the first CCA direction, the correlation between *X*^*A*^ and *Y* ^*A*^ reaches its maximum, such that *R*_ASap_ = *R*_CCA_. In this case, anatomical sampling results in the minimization of the dimensionality.

Now, let us perform anatomical sampling on the neurons. The 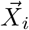 and 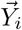 denote the functional and anatomical coordinates of the *i*^th^ neuron cluster after anatomical sampling, respectively.

To approximate, we need to calculate the functional coordinate probability distribution 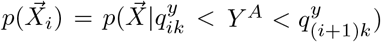, which is the distribution of the *i*^th^ neuron cluster after anatomical sampling. *Y* ^*A*^ represents the selected direction in anatomical space, and 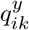 denotes the *ik*^th^ quantile of *Y* ^*A*^, where *k* is the sampled fraction. Note the following relationships and distributions:

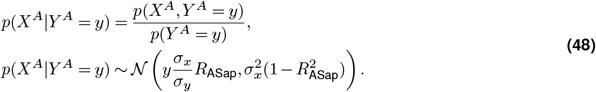

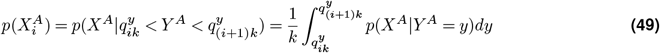

The conditional probability distribution 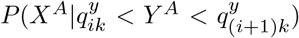 is equivalent to the distribution of the sum of 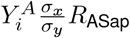 and *X*_0_, where 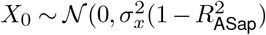):

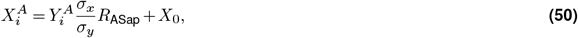

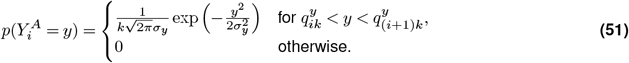

The computation of 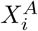 involves two technical challenges: 1. The distribution of 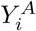 is represented by a non-elementary function (Eq. (51)), which complicates the direct calculation of 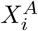, which is the sum of 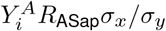 and *X*_0_. To facilitate approximation, we model 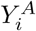 using a normal distribution with equivalent variance. 2. Calculating the variance of 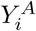 presents direct challenges, and the variance of 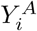 differs across different neuron clusters *i*. Using a uniform distribution for *Y* simplifies this task (this assumption is only used to calculate the variance of 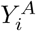). Under this assumption, the variance of 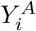 can be straightforwardly calculated as Var 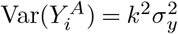. Consequently, we approximate 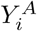 and 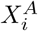 as follows:

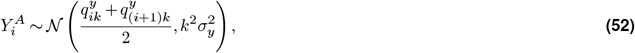

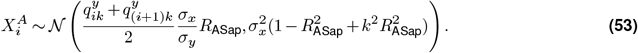

Calculating the PR dimension directly from the distribution of 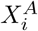 is difficult; thus, we approximate anatomical sampling with fraction *k* as functional sampling with fraction *k*_*f*_, leading to:

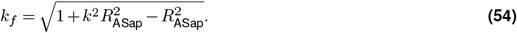

Using the equation for functional sampling 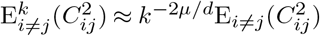 (Eq. (32)):

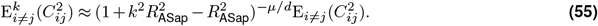

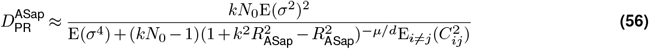

## Acknowledgments

We are grateful to Liqun Luo and Changsong Zhou for their helpful suggestions on our manuscript. QW thanks Hideaki Shimazaki for the suggestion that the functional space could be the feature space for sensory coding. QW thanks Jia Lou for improving the illustration in Figures 1 and 3. YH was supported by ECS-26303921 from the Research Grants Council of Hong Kong. QW was supported by NSFC-32071008 from the National Science Foundation of China and the STI2030-Major Projects 2022ZD0211900.

## Supplementary Materials

**Supplementary figures** comprises 24 supplementary figures.

**Supplementary note** includes additional details of the theoretical calculations and 3 supplementary figures.

**Supplementary video** includes 1 supplementary video.

## 5 Supplementary figures

**Figure S1.**
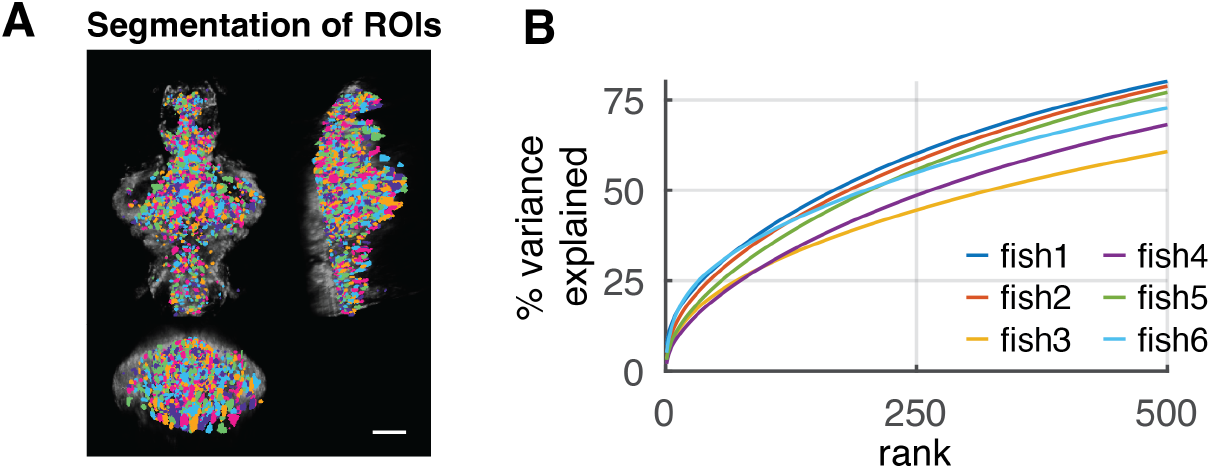
Related to Fig. 2. Experimental data description. **A**. Spatial distribution of segmented ROIs (shown in different colors). There are 1347 to 3086 ROIs in each animal. Scale bar, 100 µm. **B**. Explained variance of the activity data by PCs up to 500 rank. The different colored lines represent different fish data (n=6).

**Figure S2.**
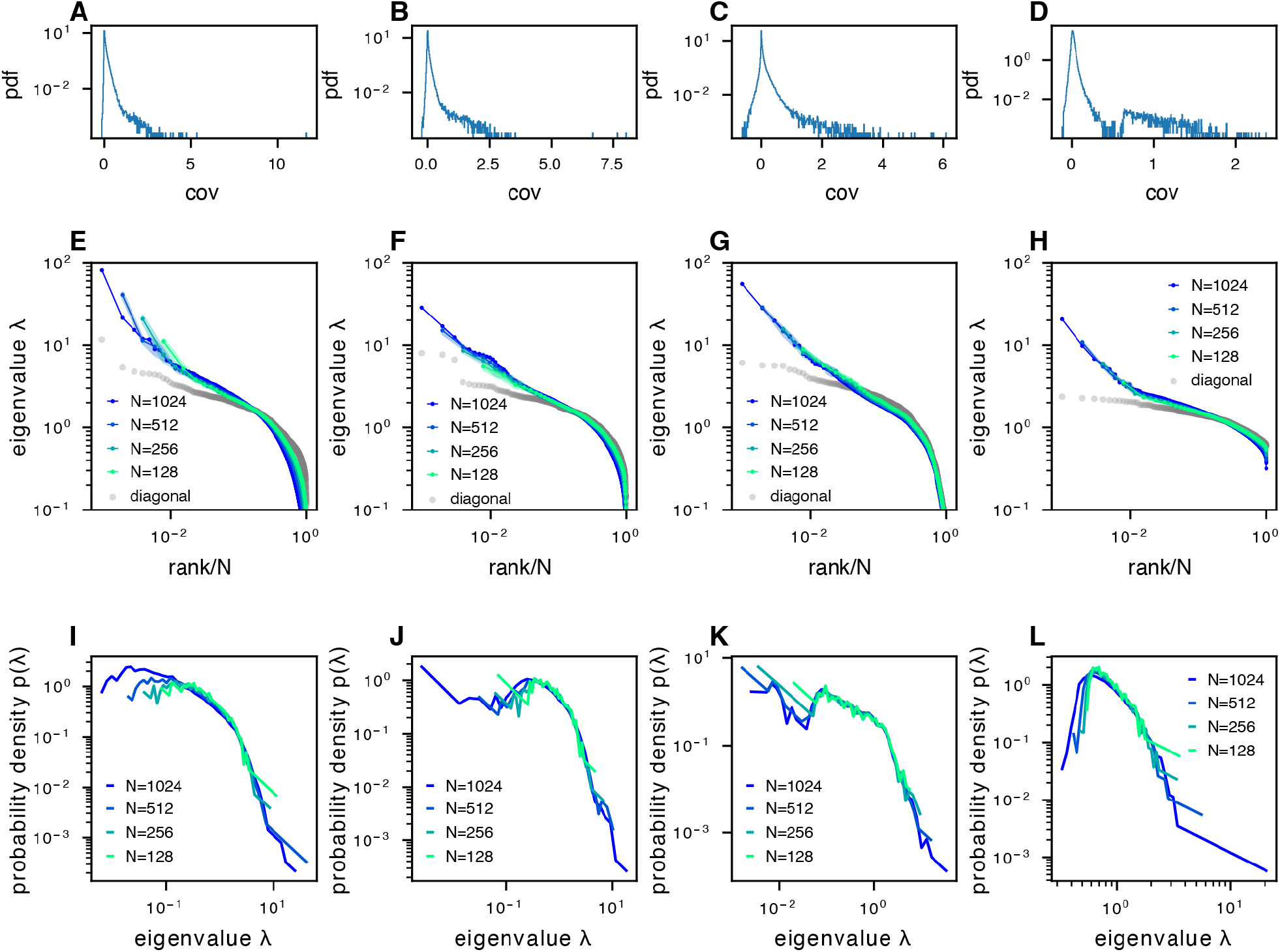
The phenomenon of scale-invariant eigenspectra across different datasets. **A-D**. Distribution of normalized pairwise covariances, where 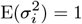 (Methods). **E-H**. Sampled covariance eigenspectra of different datasets. **I-L**. Pdfs of sampled covariance matrix eigenspectra of different datasets. The datasets correspond to the following examples: column 1: fish data (from fish 1, all fish data are shown in Fig. S10A-F) from whole brain light-field imaging; column 2: fish data from whole brain light-sheet imaging; column 3: mouse data from multi-area Neuropixels recording; column 4: mouse data from two-photon visual cortex recording.

**Figure S3.**
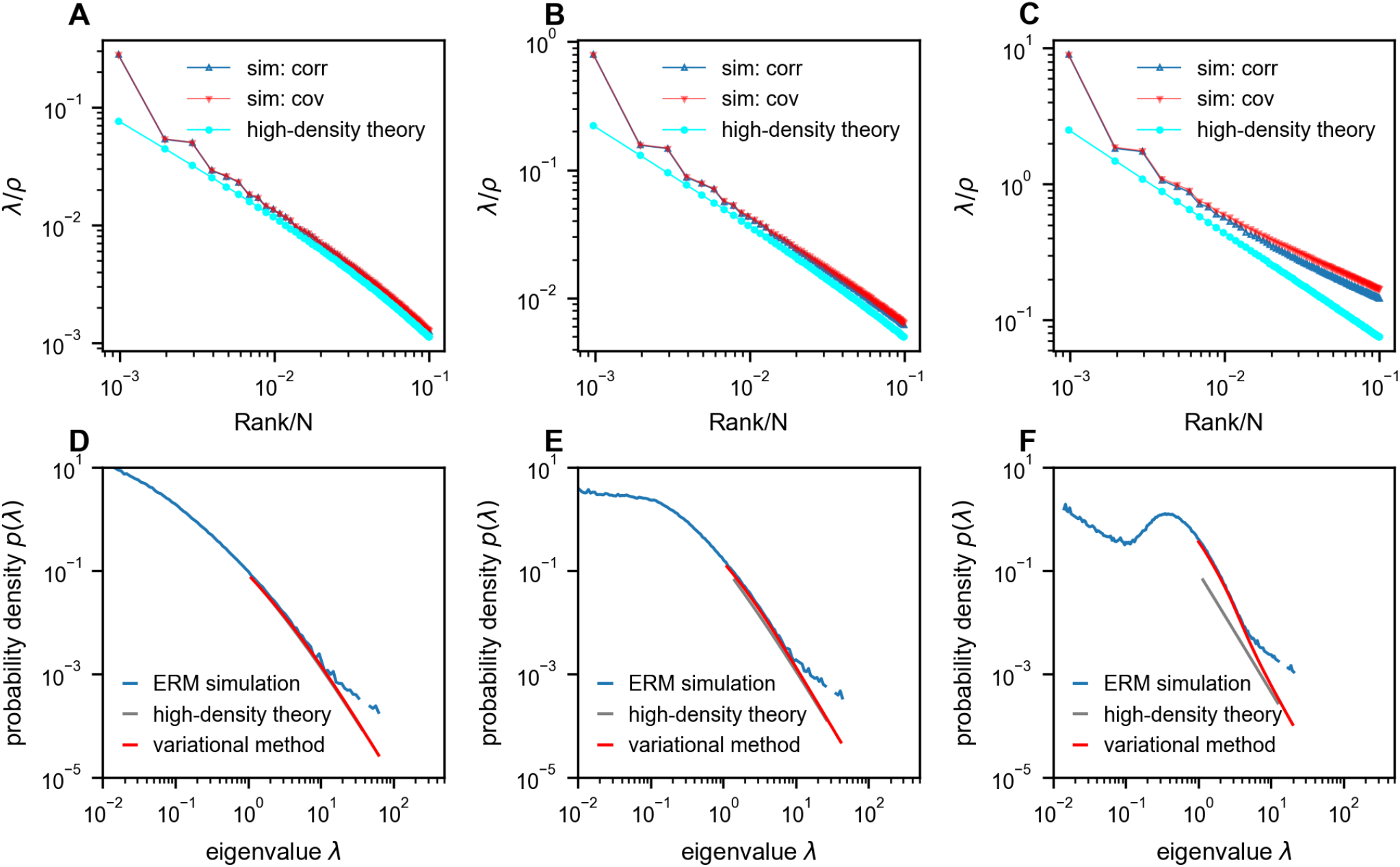
Comparison between ERM simulation and theory. **A-C**. Rank plots of the normalized eigenspectra (*λ*/ρ), with the simulations obtained using correlation matrix (sim: corr, 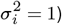 and covariance matrix (sim: cov, neuron’s activity variance 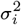 is i.i.d. sampled from a log-normal distribution with zero mean and a standard deviation of 0.5 in the natural logarithm of the 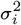 values; we also normalize 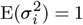 (Methods)). The curves between “sim: corr” and “sim: cov” are nearly identical in panels A and B. The theoretical predictions of normalized eigenvalues *λ*/*ρ* are obtained using the high-density theory (cyan, Eq. (12)). The density *ρ* decreases from panel A to panel C (*ρ* = 1024, 256, 10.24 respectively). **D-F**. Numerical validation of the theoretical spectrum by comparing probability density functions for increasing density of covariance ERM (*ρ* = 1024, 256, 10.24 respectively). Other simulation parameters: *N* = 1024, *d* = 2, *L* = (*N*/*ρ*)^1/*d*^, *μ* = 0.5, *ϵ* = 0.03125. The ERM simulations were conducted 100 times. The results are presented as the mean ± SEM.

**Figure S4.**
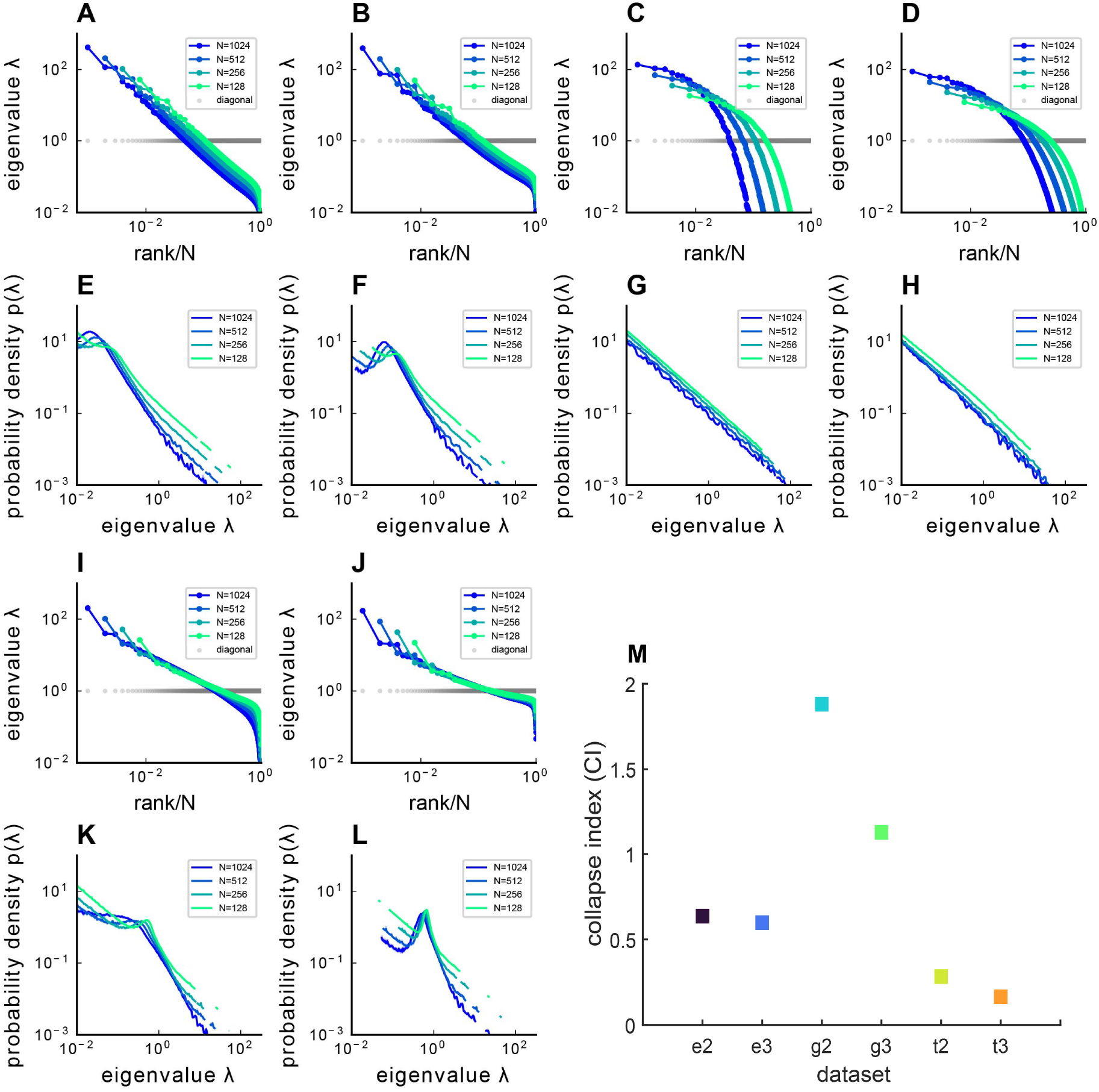
Covariance spectra under different kernel functions 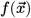. The figure presents both the sampled eigenvalue rank plot and the pdf of ERM with different functions 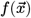 and varying dimensions *d*, where panels **A-D,I,J**. display the rank plot and panels **E-H,K,L**. show the pdf of ERM. **A,E**. Exponential function 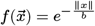 where *b* = 1 and dimension *d* = 2. **B,F**. Exponential 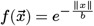 where *b* = 1 and dimension *d* = 3. **C,G**. Gaussian pdf 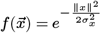 where 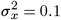 and dimension *d* = 2. **D,H**. Gaussian pdf 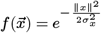 where 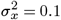 and dimension *d* = 3. **I,K**. t pdf (Eq. (11)) and dimension *d* = 2. **J,L**. t pdf (Eq. (11)) and dimension *d* = 3. The ERM simulations were conducted 100 times and each ERM used an identical sampling technique described in (Methods). The results represent mean ± SEM. **M**. Summary of CI’s for different 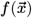 and *d*. On the x-axis labels, ‘e’ denotes the Exponential function 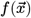,’g’ denotes the Gaussian pdf 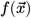,’t’ denotes the t-distribution pdf 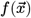,while ’2’ and ’3’ indicate *d* = 2 or *d* = 3, respectively.

**Figure S5.**
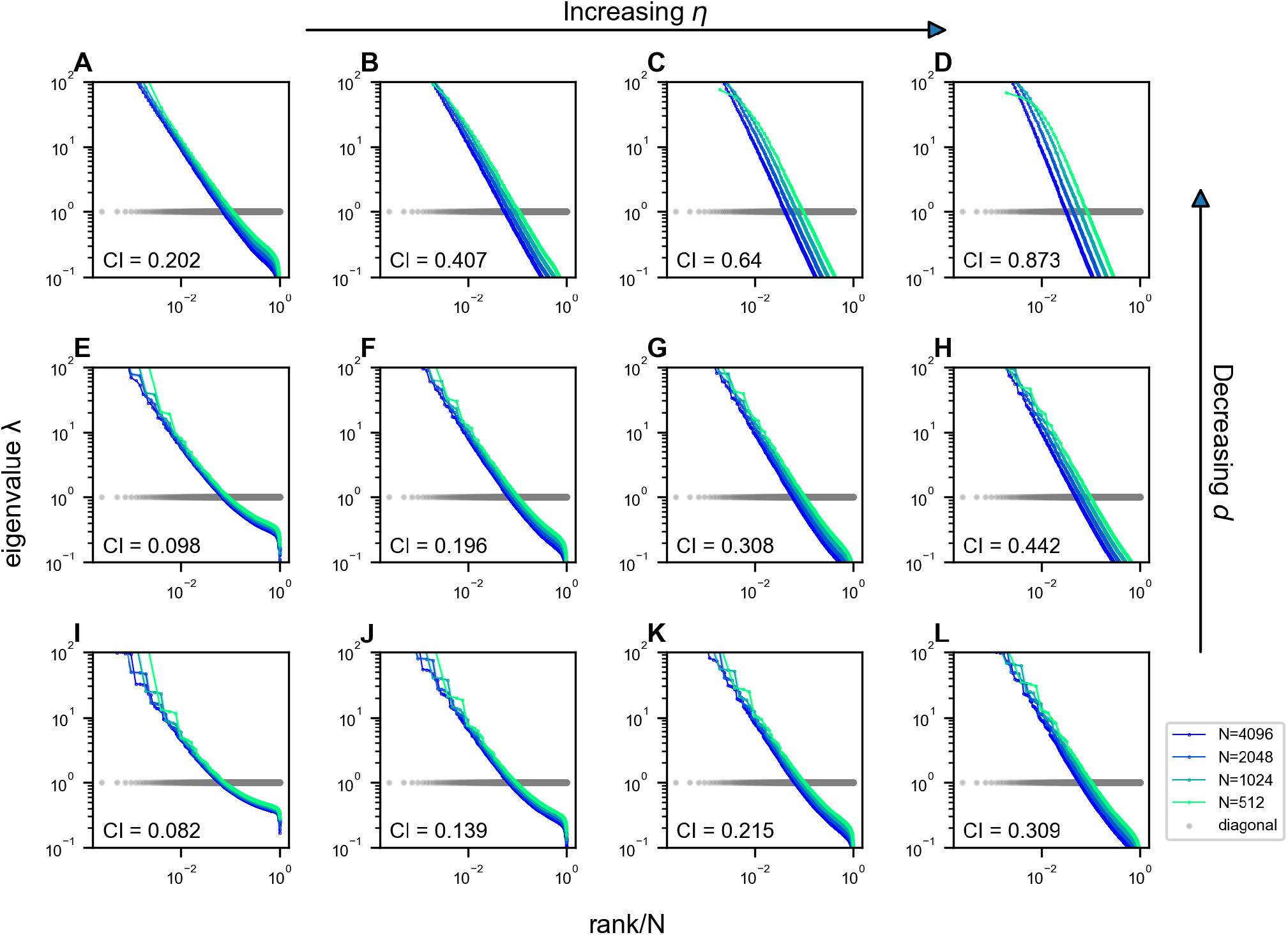
Impact of *η* and *d* on the scale invariance of covariance eigenspectra in the ERM with 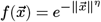. The columns from left to right correspond to *η* = 0.3, 0.5, 0.7, 0.9, and the rows from top to bottom correspond to *d* = 1, 2, 3 (Eq. (2) and Eq. (11)). Other ERM simulation parameters: *N* = 4096, *ρ* = 256, *L* = (*N*/*ρ*)^1/*d*,^ *ϵ* = 0.03125 and *σ*^2^ = 1. Each panel shows a single ERM realization. For visualization purposes, the views in some panels are truncated since we use the same range for the eigenvalues in all panels.

**Figure S6.**
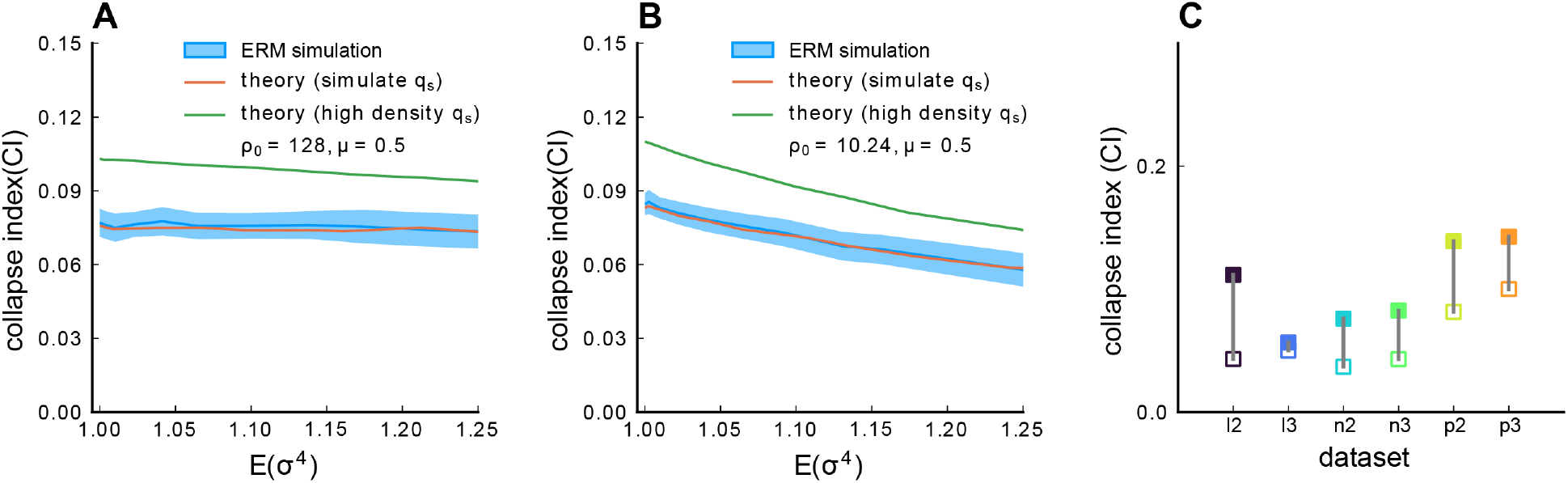
Impact of heterogeneous activity levels on the scale invariance. **A**. The CI as a function of the heterogeneity of neural activity levels 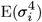.We generate ERM where each neuron’s activity variance 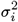 is i.i.d. sampled from a log-normal distribution where the logarithm of the variable follows a normal distribution with zero mean and a sequence of standard deviation (0, 0.05, 0.1,…, 0.5) in the natural logarithm of the values 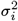.We also 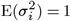 (Methods). The solid blue line is the average across 100 ERM simulations, and the shaded area represents the SD. The red line results from the Gaussian variational method with simulation value integration limit 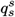.The green line is the result of the Gaussian variational method with high-density value integration limit 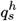 (Methods). *ρ*_0_ = 128. **B**. Same as A, but with a smaller *ρ*_0_ = 10.24. Other parameters: *μ* = 0.5, *d* = 2, *N* = 1024, *L* = (*N*/*ρ*)^1/*d*^, *ϵ* = 0.03125. **C**. The collapse index (CI) of the correlation matrix (filled symbols) is larger than that of the covariance matrix (opened symbols) across different datasets excluding those shown in Fig. 4. We use 7,200 time frame data across all the datasets. l2 to l3: light-sheet zebrafish data (2 Hz per volume); n2 to n3: Neuropixels mouse data, downsampled to 10 Hz per volume, p2 to p3: two-photon mouse data, (3 Hz per volume).

**Figure S7.**
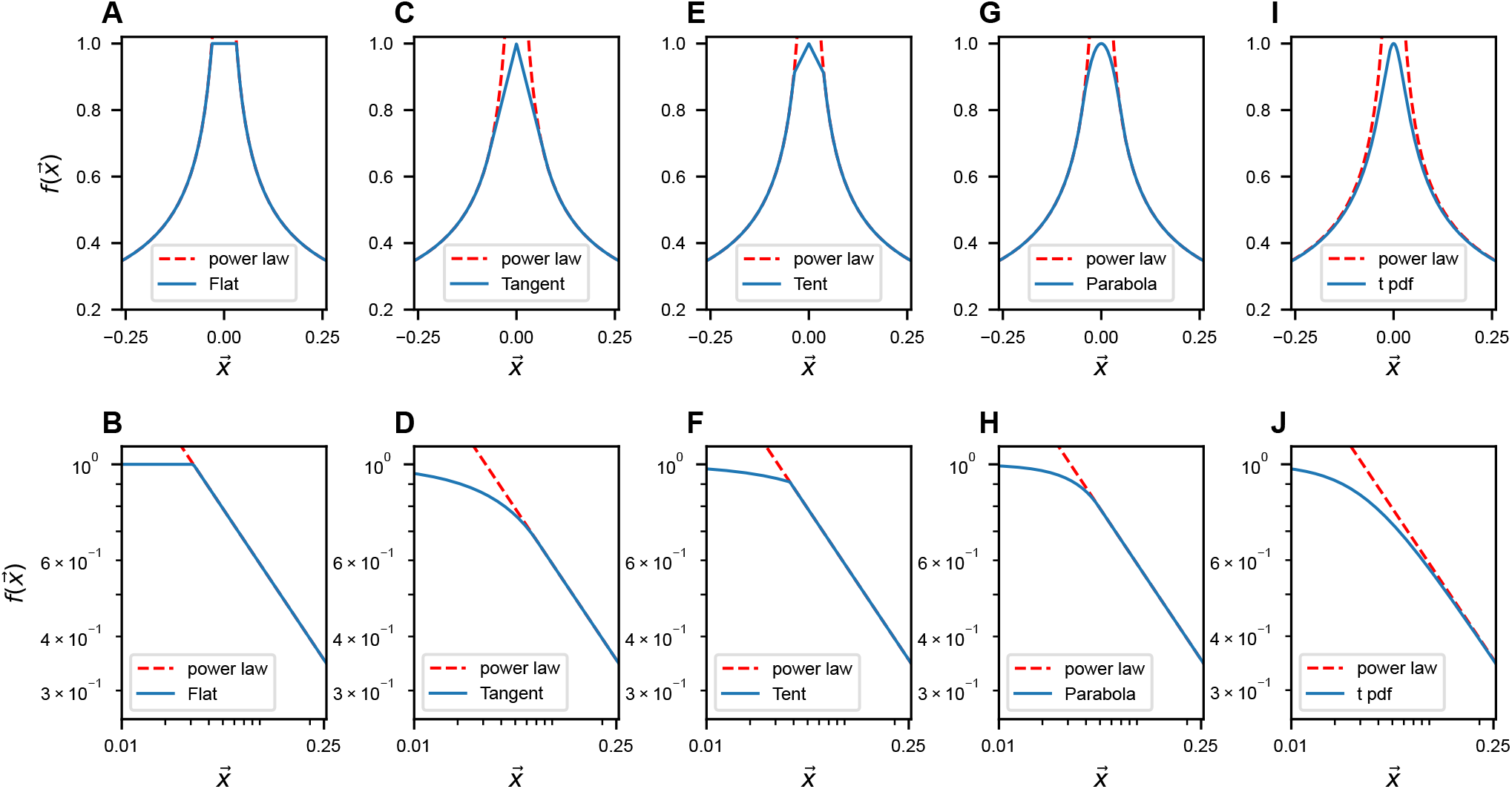
Modifications of 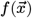 near *x* = 0. The upper row illustrates the slow-decaying kernel function 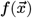 (blue solid line) and its power-law asymptote (red dashed line) along a 1D slice at various 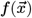.The lower row is similar to **A**, but on the log-log scale. The formulas for different 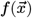’s are listed in table S3 in Methods.

**Figure S8.**
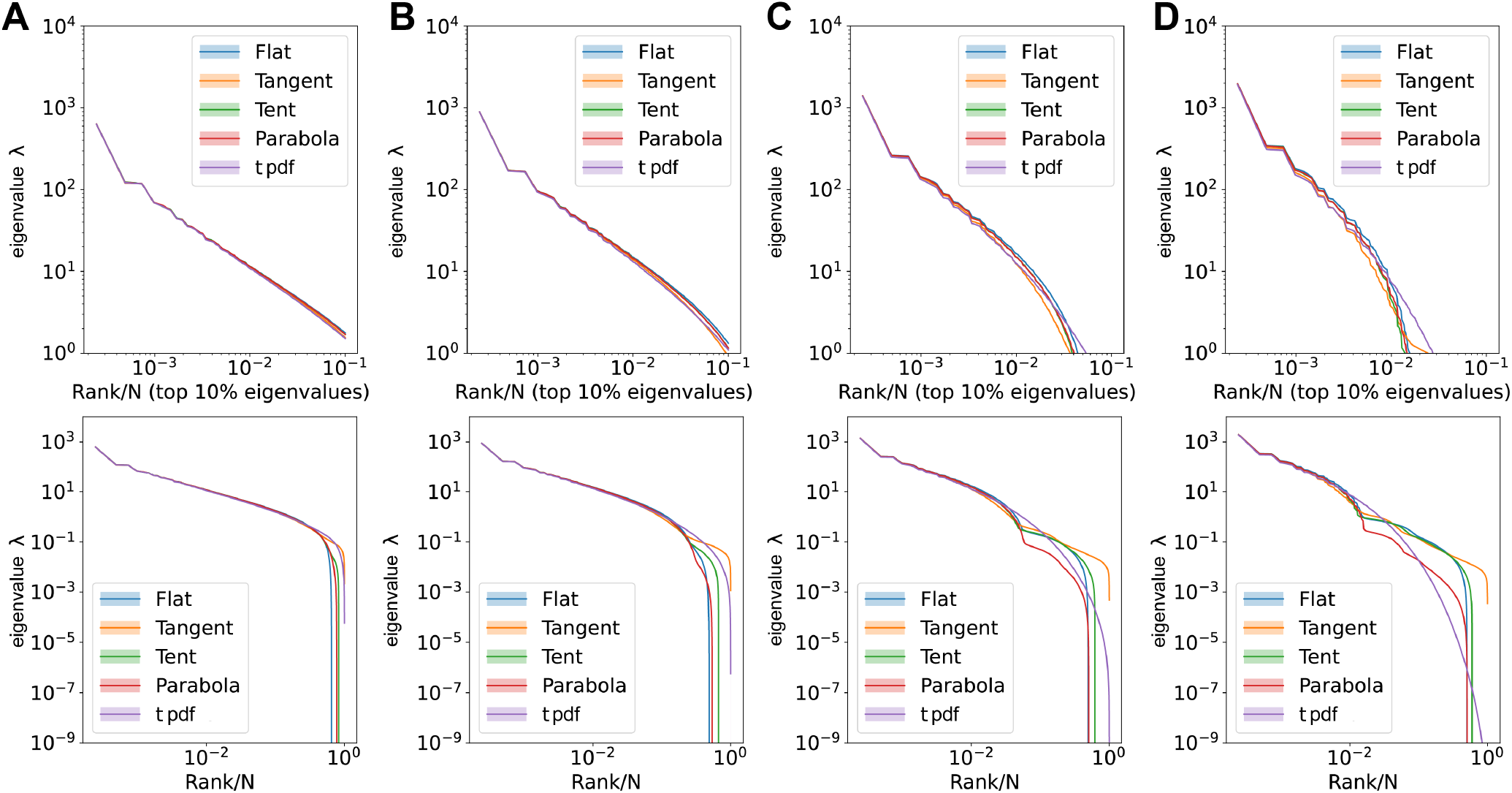
Comparisons of large eigenvalues across different smoothing interval sizes, *ϵ*. Rank plot (upper row) and pdf (lower row) of the covariance eigenspectrum for ERMs with different 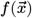.**A**. *ϵ* = 0.06. **B**. *ϵ* = 0.12. **C**. *ϵ* = 0.3. **D**. *ϵ* = 0.6. Other ERM simulation parameters: *N* = 4096, *ρ* = 100, *μ* = 0.5, *d* = 2, *L* = 6.4, 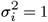.The formulas for different 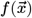’s are listed in table S3 in Methods.

**Figure S9.**
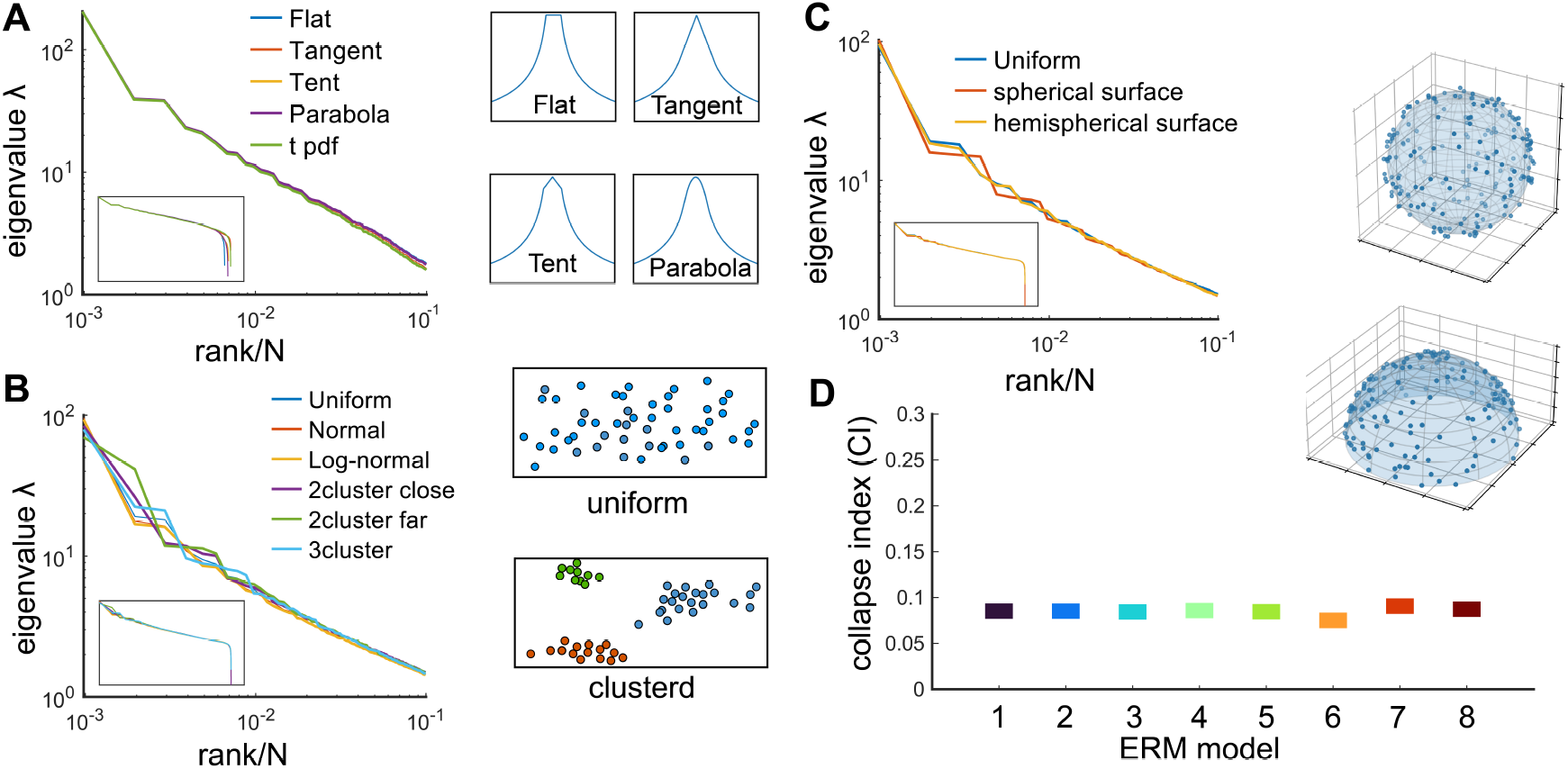
Factors that do not affect the scale invariance. **A**. Rank plot of the covariance eigenspectrum for ERMs with different 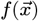 (see table S3). Diagrams show different slow-decaying kernel functions 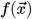 along a 1D slice. **B**. Same as **A** but for different coordinate distributions in the functional space (see text). The diagrams on the right illustrate uniform and clustered coordinate distributions. **C**. Same as **A** but for different geometries of the functional space (see text). Diagrams illustrate spherical and hemispherical surfaces. **D**. CI of the different ERMs considered in A-C. The range on the y-axis is identical to Fig. 4C. On the x-axis, 1: uniform distribution, 2: normal distribution, 3: log-normal distribution, 4: uniform two nearby clusters, 5: uniform two faraway clusters, 6: uniform 3-cluster, 7: spherical surface in ℝ^3^, 8: hemispherical surface in ℝ^3^. All ERM models in **B, C** are adjusted to have a similar distribution of pairwise correlations (Methods).

**Figure S10.**
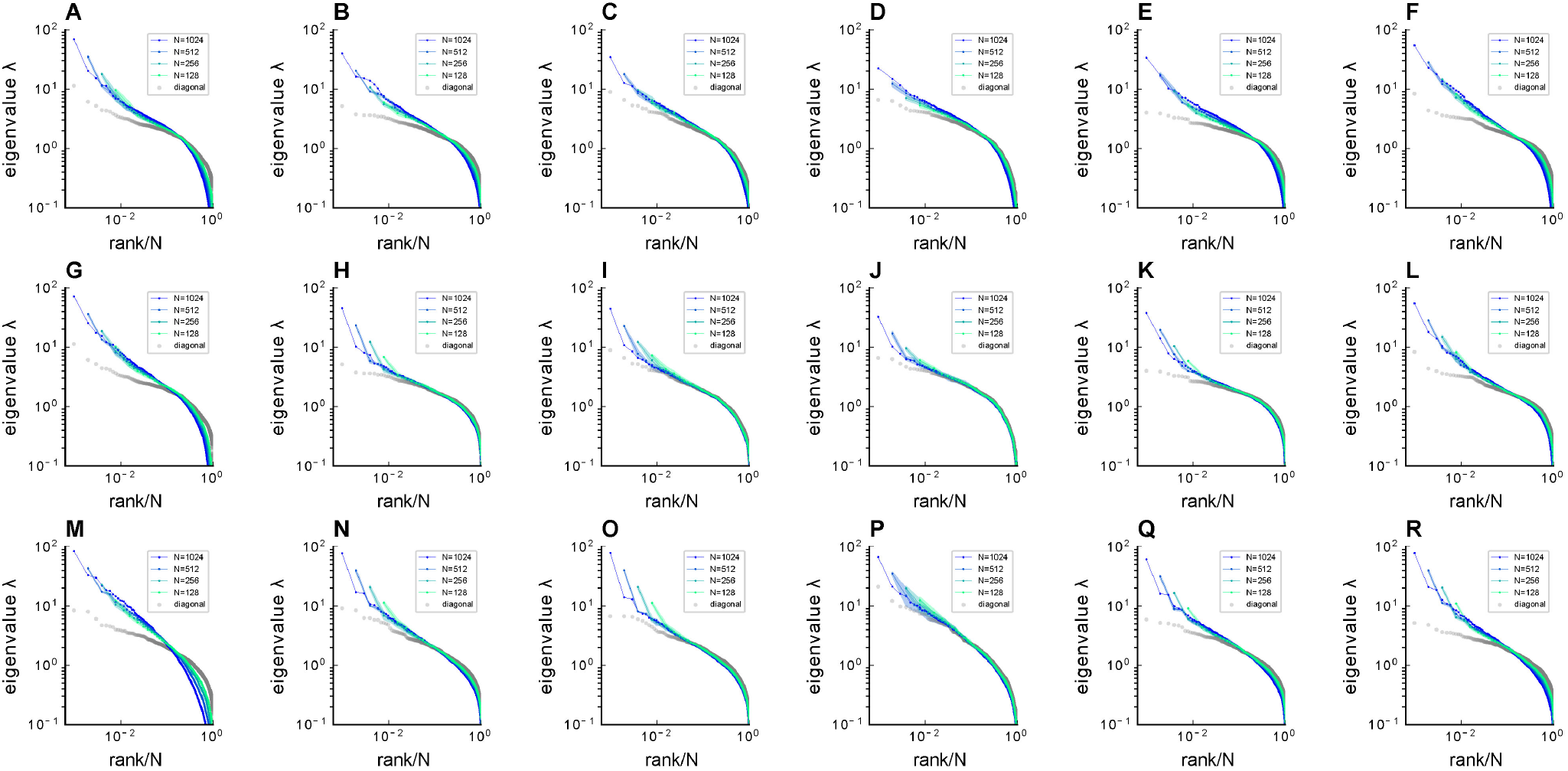
Fitting ERM to zebrafish data from our experiments (part 1). Comparison of sampled covariance eigenspectra in fish data and fitted ERM models. The columns correspond to six light-field zebrafish data: fish 1 to fish 6. Number of time frames: fish 1 - 7495, fish 2 - 9774, fish 3 - 13904, fish 4 - 7318, fish 5 - 7200 and fish 6 - 9388. **A-F**. sampled covariance eigenspectra for different fish data. **G-L**. Same as A-F but for ERM models with fitted parameters (*μ*/*d, L*), functional coordinates inferred using MDS, and the experimental *σi*. **M-R**. Same as A-F but for ERM models with fitted parameters (*μ*/*d, L*), uniform distributed functional coordinates, and a log-normal distribution of *σ*^2^. *μ*/*d* = [0.456, 0.258, 0.205, 0.262, 0.302, 0.308] in fish 1-6.

**Figure S11.**
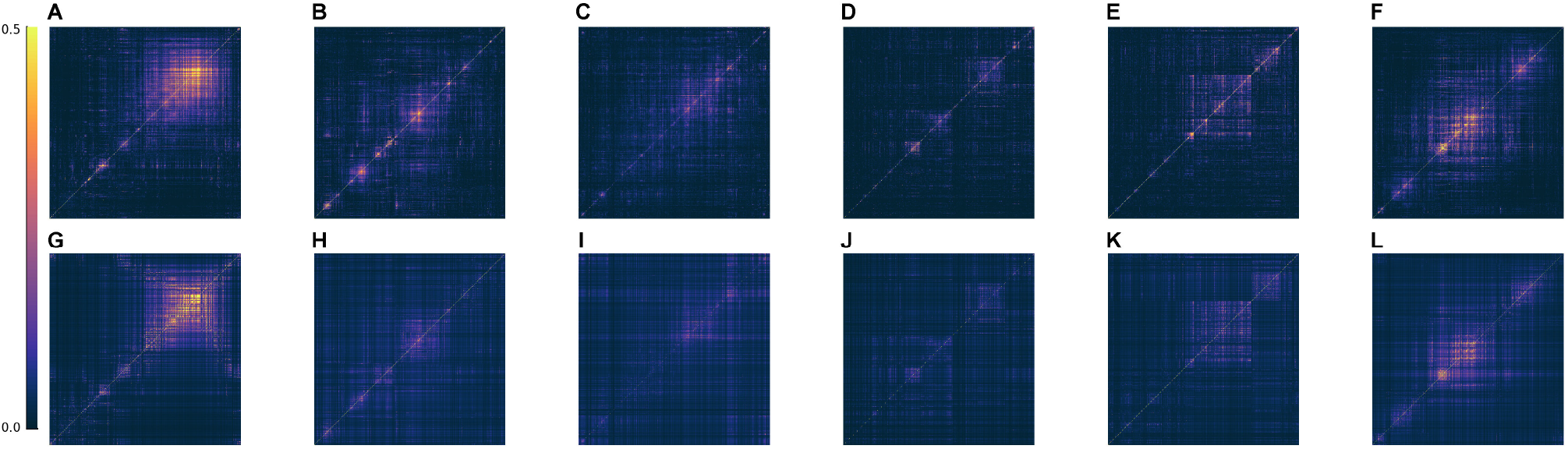
Fitting ERM to all six zebrafish data from our experiments (part 2). Comparison of the covariance matrix between fish data and our fitted model. The columns correspond to six light-field zebrafish data: fish 1 to fish 6. **A-F**. The covariance matrix of different fish data. **G-L**. The covariance matrix of ERM models with fitted parameters (*μ, L*) and functional coordinates inferred using MDS and the experimental *σi*.

**Figure S12.**
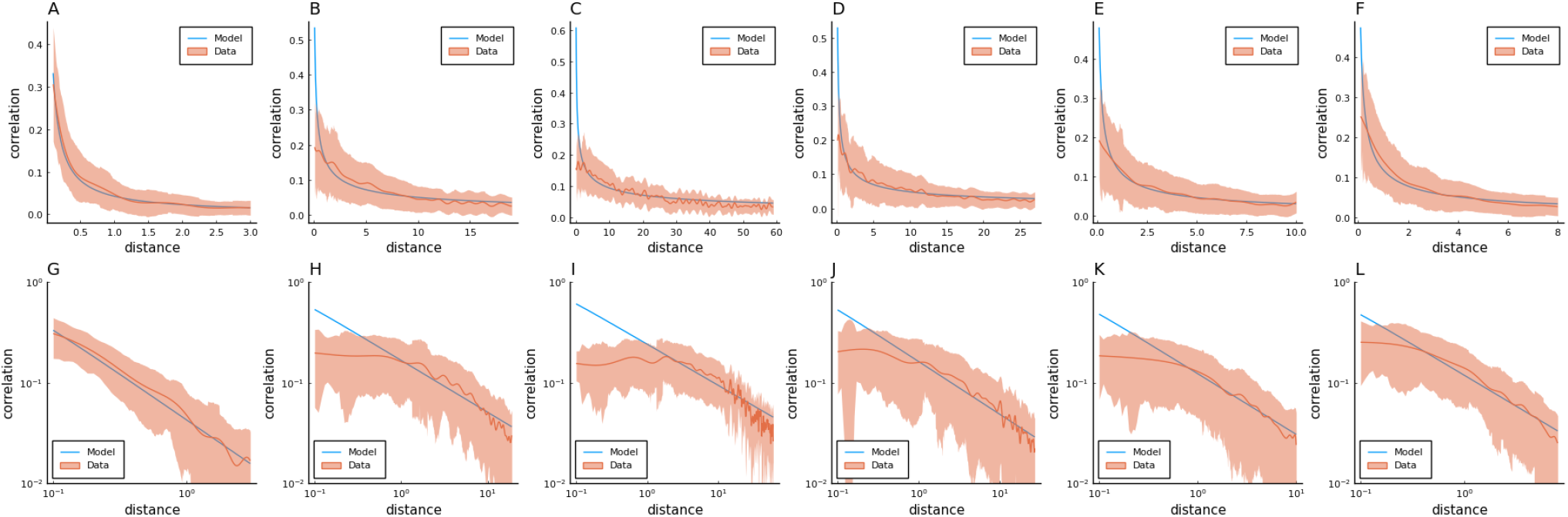
Fitting ERM to all six zebrafish data from our experiments (part 3). Columns correspond to five light-field zebrafish data: fish 1 to fish 6. **A-F**: Comparison of the power-law kernel 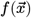 in the model (blue line) and the correlation-distance relationship in the data (red line). The distance is calculated from the inferred coordinates using MDS. The shaded area represents the SD. **G-L**: Same as A-D but on the log-log scale.

**Figure S13.**
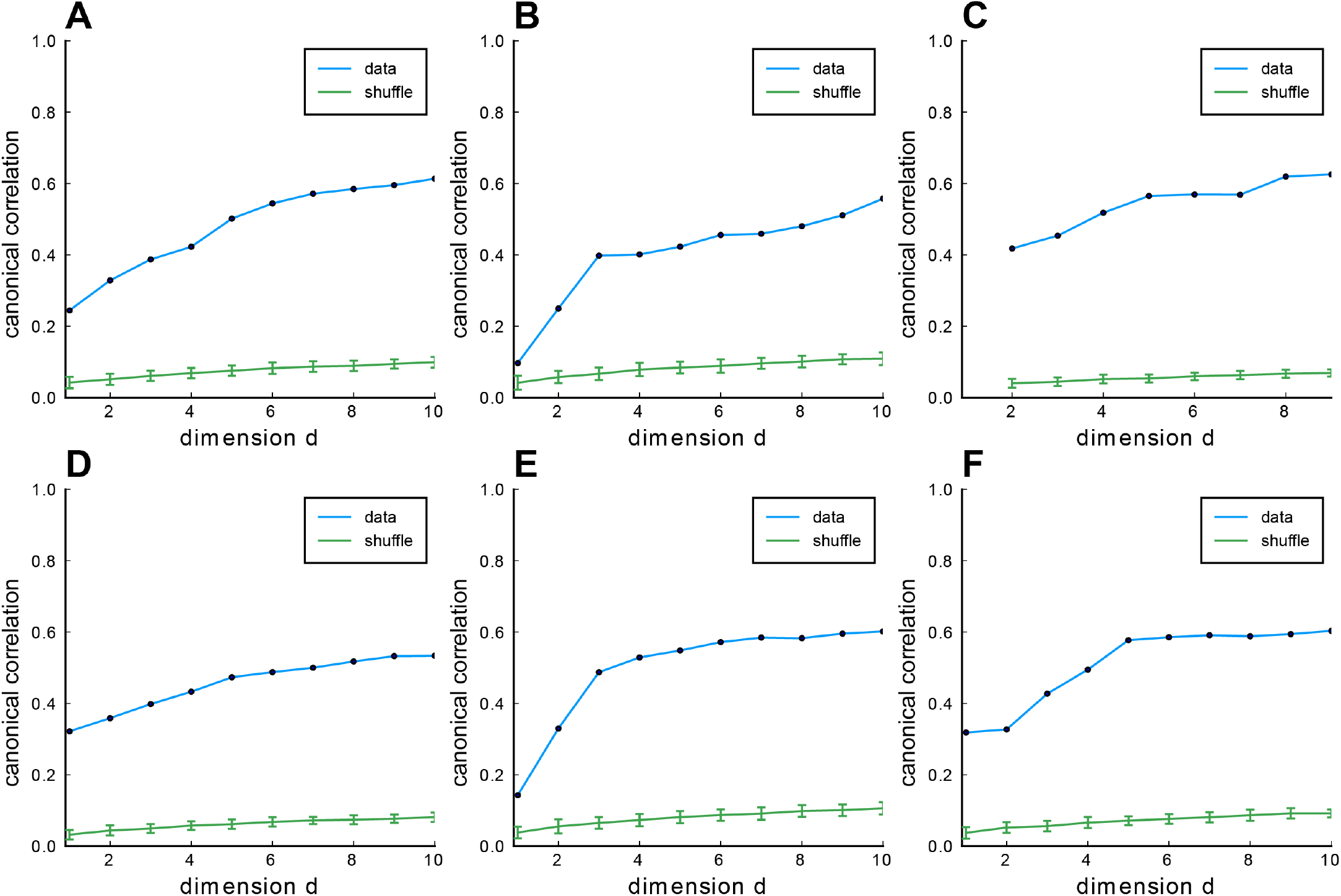
Fitting ERM to all six zebrafish data from our experiments (part 4). Columns correspond to 6 light-field zebrafish data: fish 1 to fish 6. **A-F**: CCA correlation between the first CCA variables with different embedding dimensions in the functional space. Blue line indicates the CCA correlation of example fish data, green line shows the CCA correlation of example fish data with shuffled functional coordinates, and error bars represent the SD.

**Figure S14.**
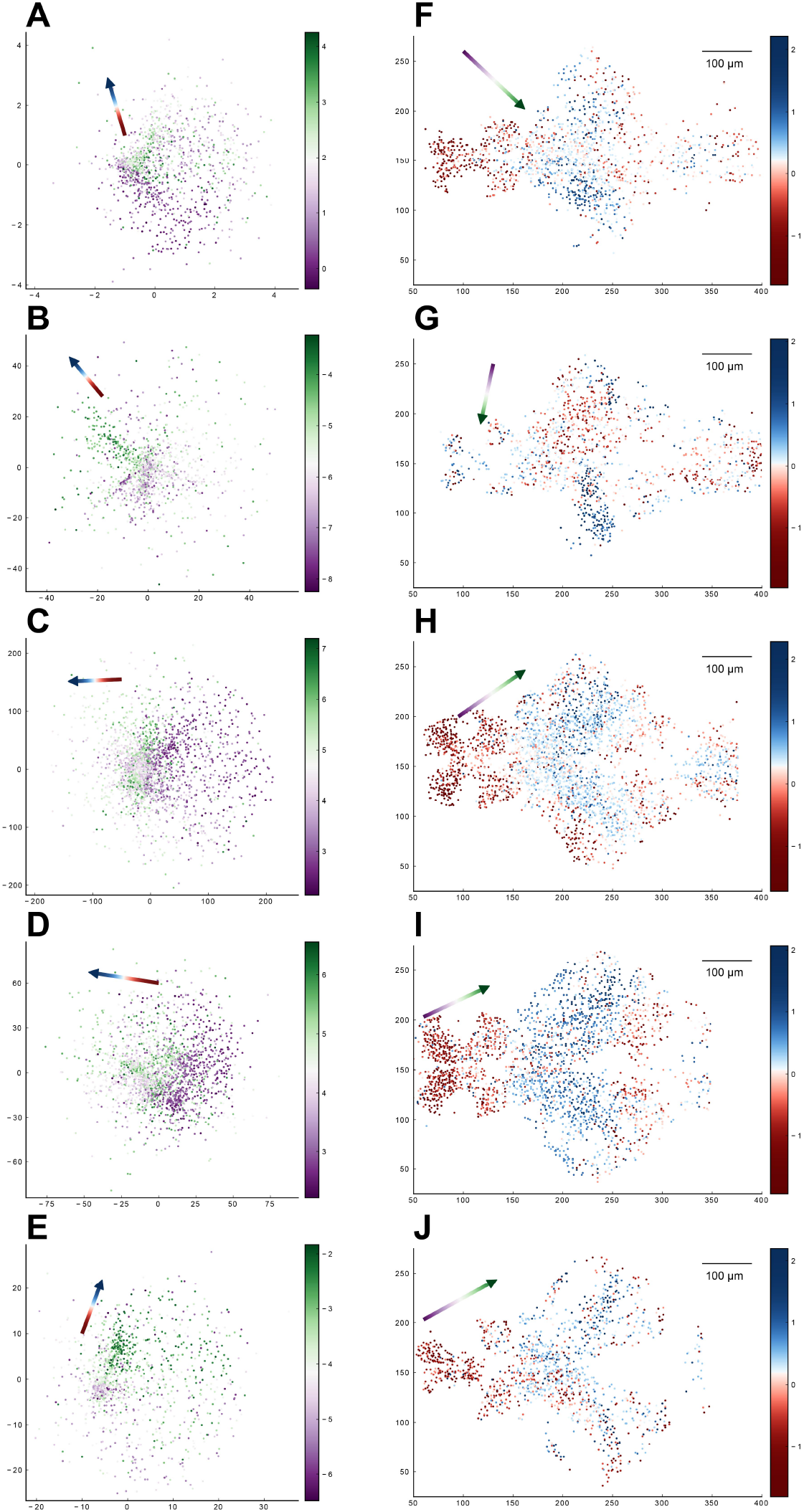
Relationship between the functional space and anatomical space for each zebrafish dataset from our experiments. Columns correspond to five light-field zebrafish data: fish 1 to fish 5 (with fish 6 has been shown in Fig. 5). **A-E**. Distribution of neurons in the functional space, where each neuron is color-coded by the projection of its coordinate along the canonical axis 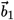 in anatomical space (see text in Result section 2.4). Arrow: the first CCA direction 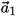 in functional space. **F-J**. Distribution of neurons in the anatomical space with the forebrain neuron located on the left side and the hindbrain neuron on the right side. Each neuron is color-coded by the projection of its coordinate along the canonical axis 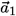 in functional space (see text in Result section 2.4). Arrow: the first CCA direction 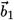 in anatomical space.

**Figure S15.**
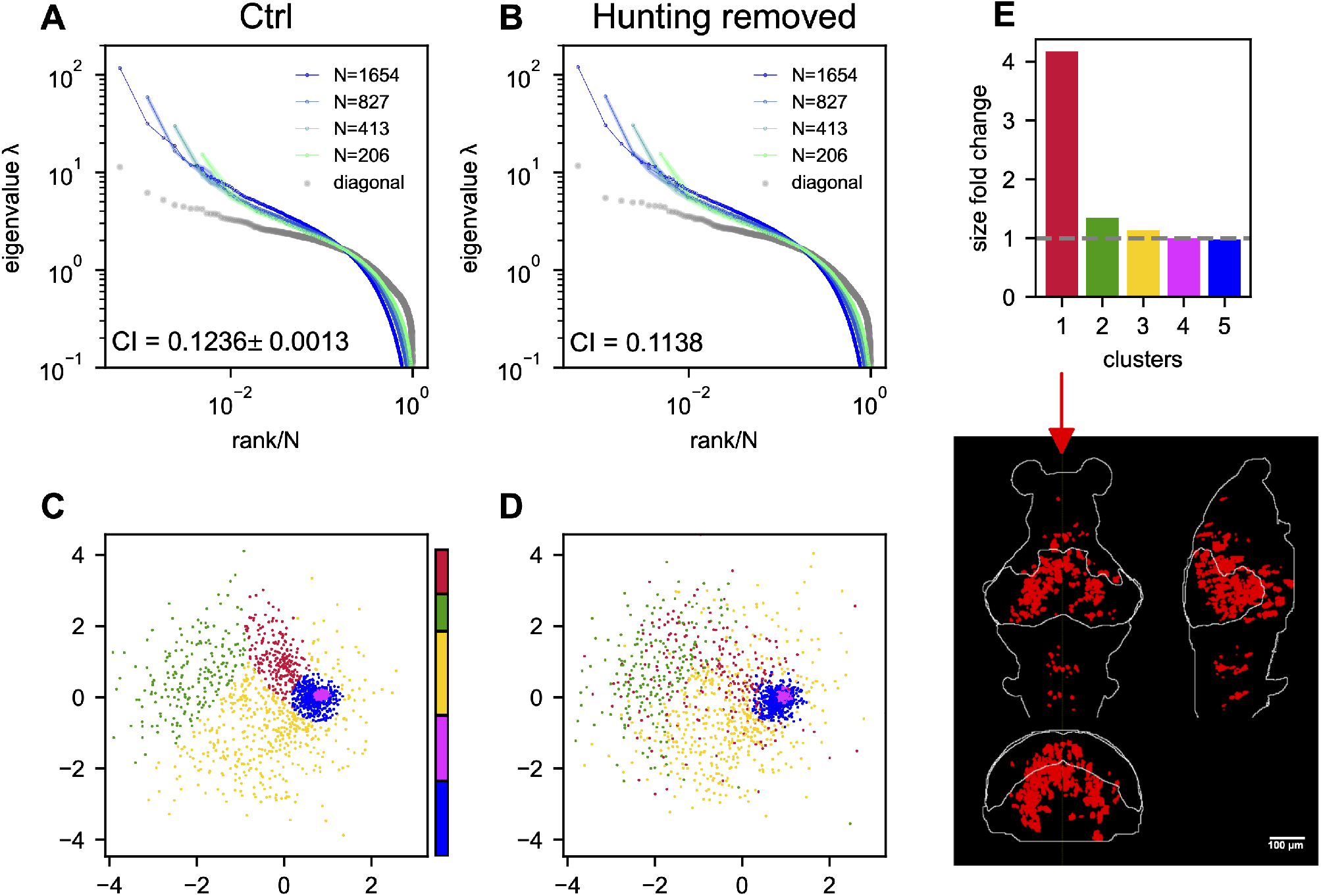
The effects of hunting behavior on scale invariance and functional space organization. **A,B**. sampled covariance eigenspectra of the data from fish 1 calculated from control (**A**) and hunting removed (**B**) data. Ctrl: We randomly remove the same number of non-hunting frames. This process is repeated 10 times, and the mean±SD of the CI is shown in the plot. Hunting removed: The time frames corresponding to the eye-converged intervals (putative hunting state) are removed when calculating the covariance (Methods). The CI for the hunting-removed data appears to be statistically smaller than in the control case (p-value= 1.5 × 10^−9^). **C**. Functional space organization of control data. The neurons are clustered using the Gaussian Mixture Models (GMMs) and their cluster memberships are shown by the color. The color bar represents the proportion of neurons that belong to each cluster. **D**. Similar to **C** but the functional coordinates are inferred from the hunting-removed data. The color code of each neuron is the same as that of the control data (**C**), which allows for a comparison of the changes to the clusters under the hunting-removed condition. See also the Movie. S1. **E**. Fold change in size / area (Methods) for each cluster (top; the gray dashed line represents a fold change of 1, that is, no change in size) and the anatomical distribution of the most dispersed cluster (bottom).

**Figure S16.**
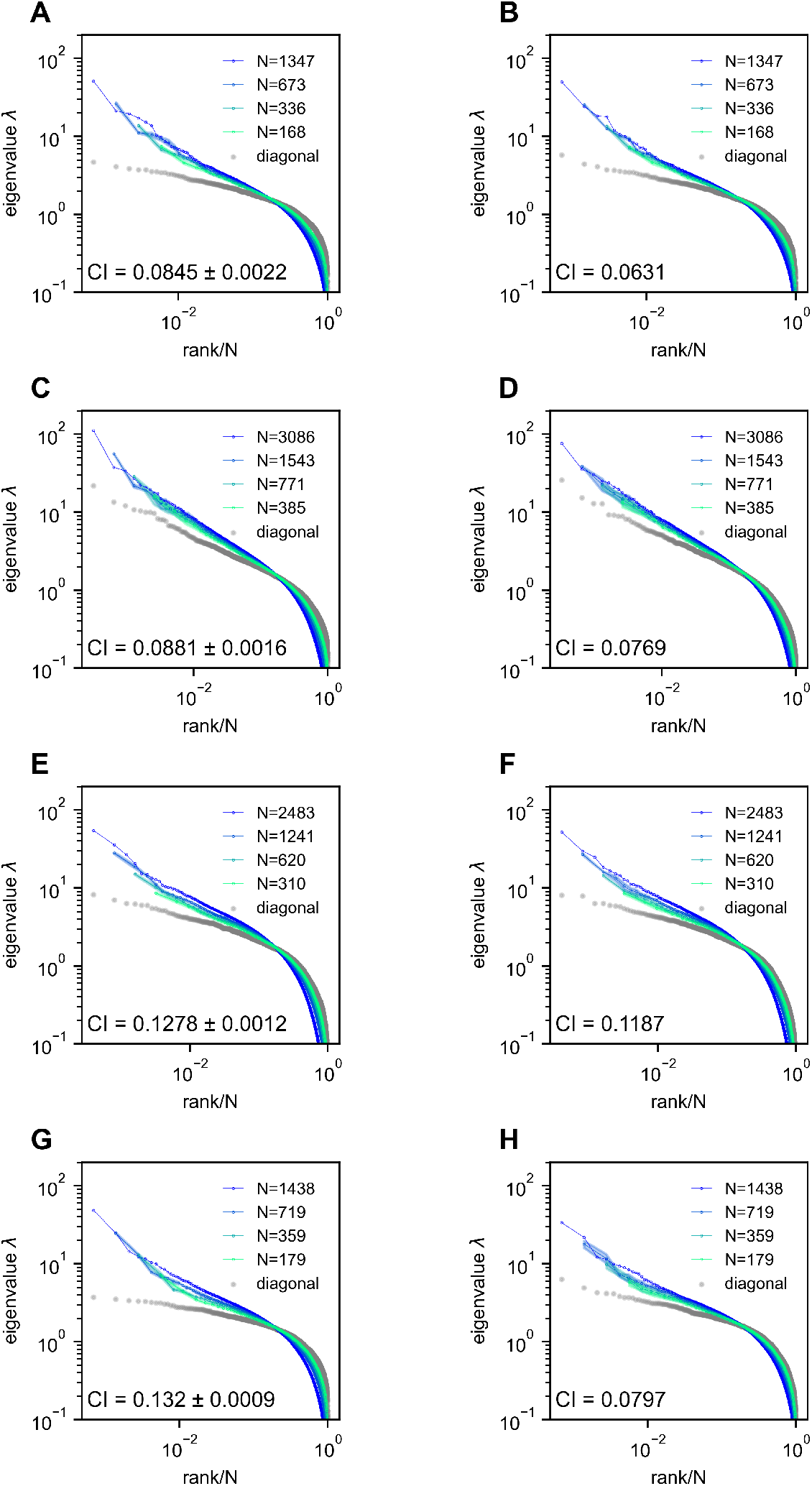
Removing the time segment of hunting behavior does not obliterate the scale-invariant eigenspectra. Rows correspond to 4 light-field zebrafish data: fish 2 to fish 5 (results for fish 1 have been shown in Fig. S15). **A,C,E,G**. Ctrl: we randomly remove the same number of time frames that are *not* the putative hunting frames. We repeat this process 10 times to generate 10 control covariance matrices and the CI is represented by mean±SD. **B,D,F,H**. Hunting removed: data obtained by removing hunting frames from the full data (Methods). The CI for the hunting removed data appears to be significantly smaller than that of the control case (one-sample t-test *p* = 2.2 × 10^−10^ in fish 2, *p* = 4.6 × 10^−9^ in fish 3, *p* = 1.7 × 10^−9^ in fish 4, and *p* = 3.4 × 10^−17^ in fish 5).

**Figure S17.**
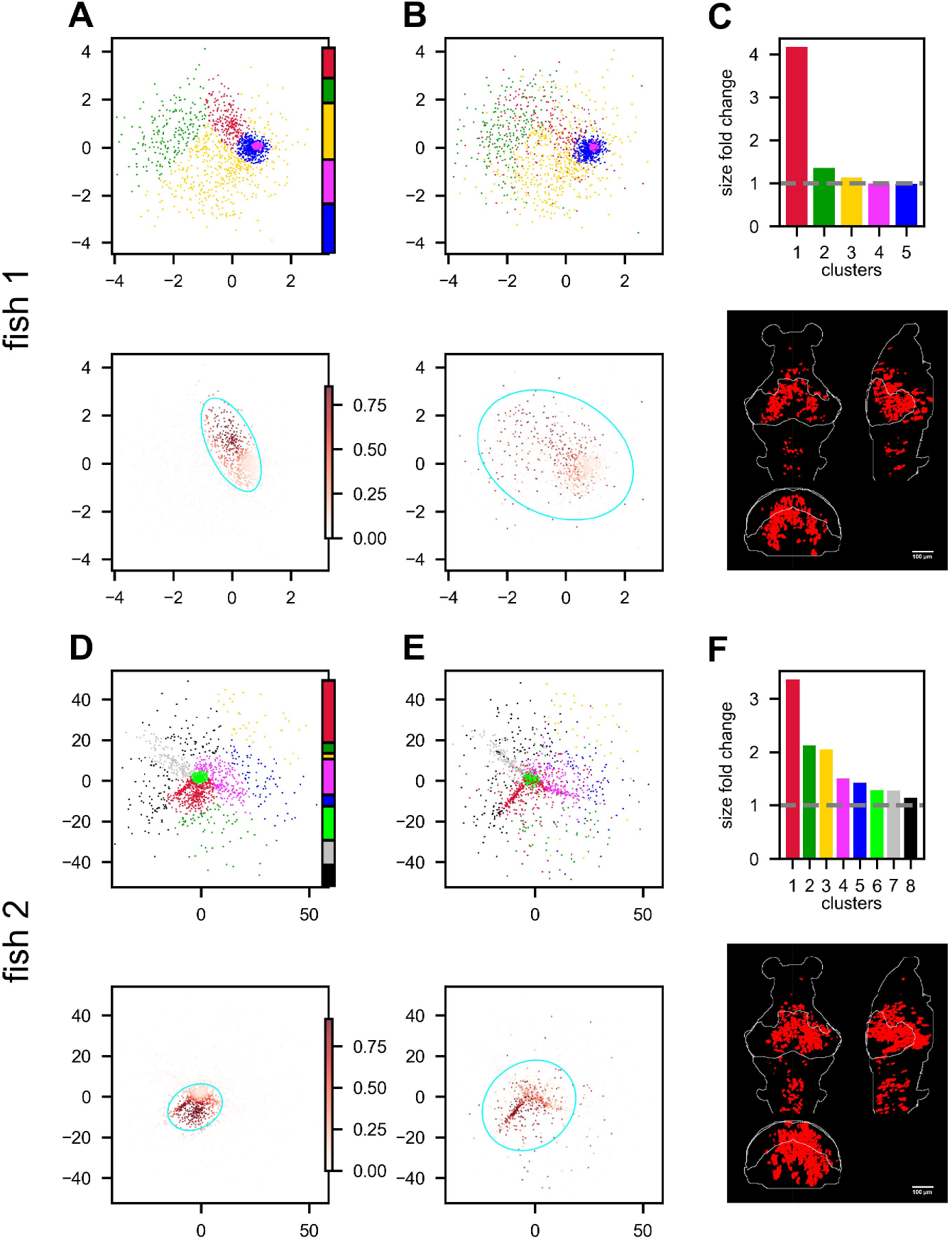
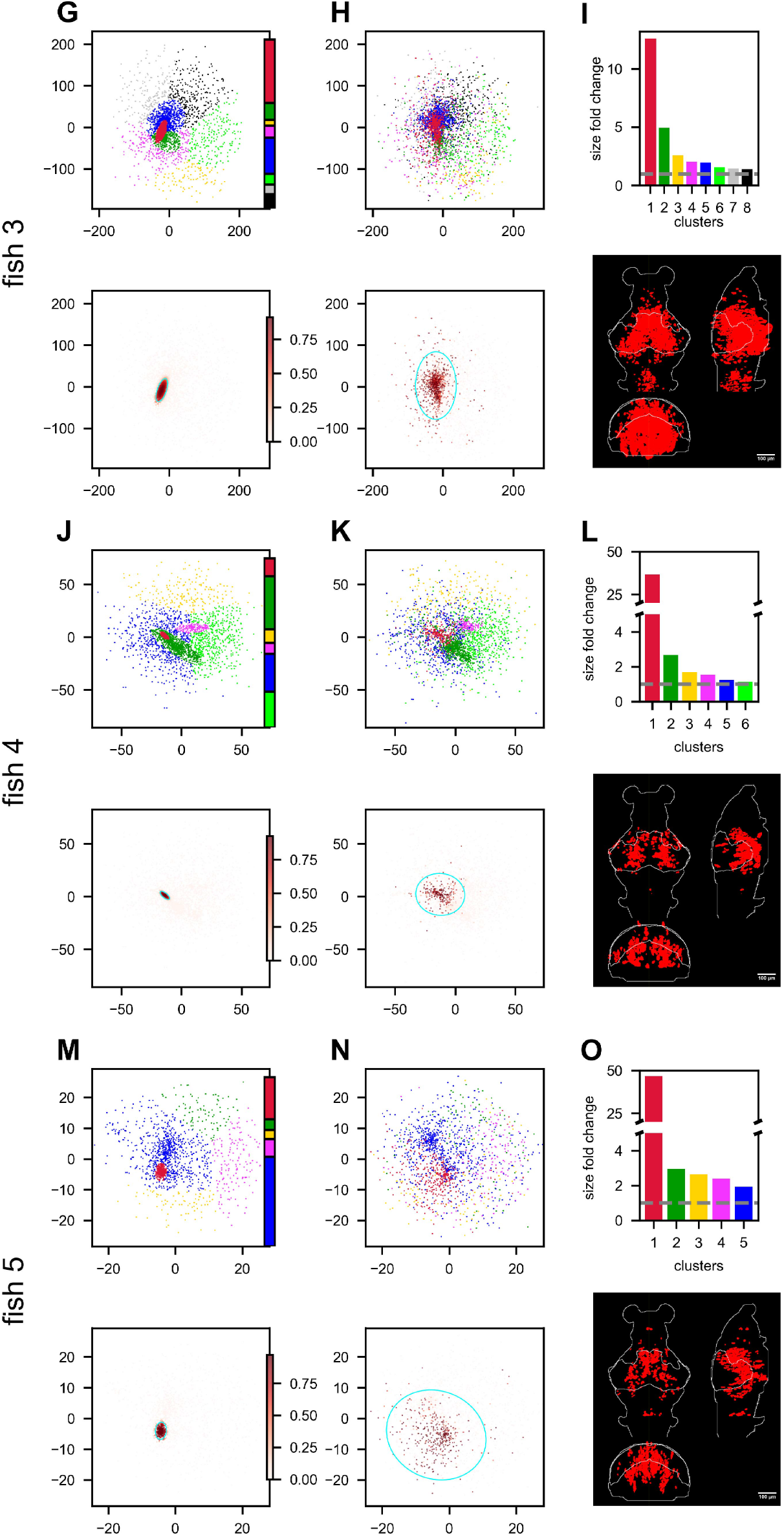
Hunting behavior reorganizes neurons in the functional space. Rows correspond to 5 light-field recordings of zebrafish engaged in hunting behavior: fish 1 to fish 5. **A,D,G,J,M. (top)** Functional space organization of the control data inferred by fitting the ERM and MDS (section 2.4). Neurons are clustered using the Gaussian Mixture Models (GMMs) and their cluster memberships are shown by the color. The colorbar represents the proportion of neurons belonging to each cluster. **A,D,G,J,M. (bottom)** The coordinate distribution of the cluster in control data which is most dispersed (i.e., largest fold change in size, see below) after hunting-removal. The transparency of the dots (colorbar) is proportional to the probability of the neurons belonging to this cluster (Methods). The cyan ellipse serves as a visual aid for the cluster size: it encloses 95% of the neurons belonging to that cluster (Methods). **B,E,H,K,N. (top)** Similar to **A,D,G,J,M. (top)** but the functional coordinates are inferred from the hunting-removed data. The color code of each neuron is the same as that in the control data, which allows for a comparison of the changes to the clusters under the hunting-removed condition. **B,E,H,K,N. (bottom)** Similar to **A,D,G,J,M. (bottom)** but the functional coordinates are inferred from the hunting-removed data. The transparency of each neuron is the same as in **A,D,G,J,M. (bottom)**, and it represents the probability *pik* (Methods) of neurons belonging to the most dispersed cluster *k* in the control data. Likewise, the cyan ellipse encloses 95% of the neurons belonging to that cluster (Methods). **C,F,I,L,O**. Top, size/area fold change (Methods) for each cluster (the gray dashed line represents a fold change of 1, i.e., no change in size); bottom, the anatomical distribution of the neurons in the most dispersed cluster.

**Figure S18.**
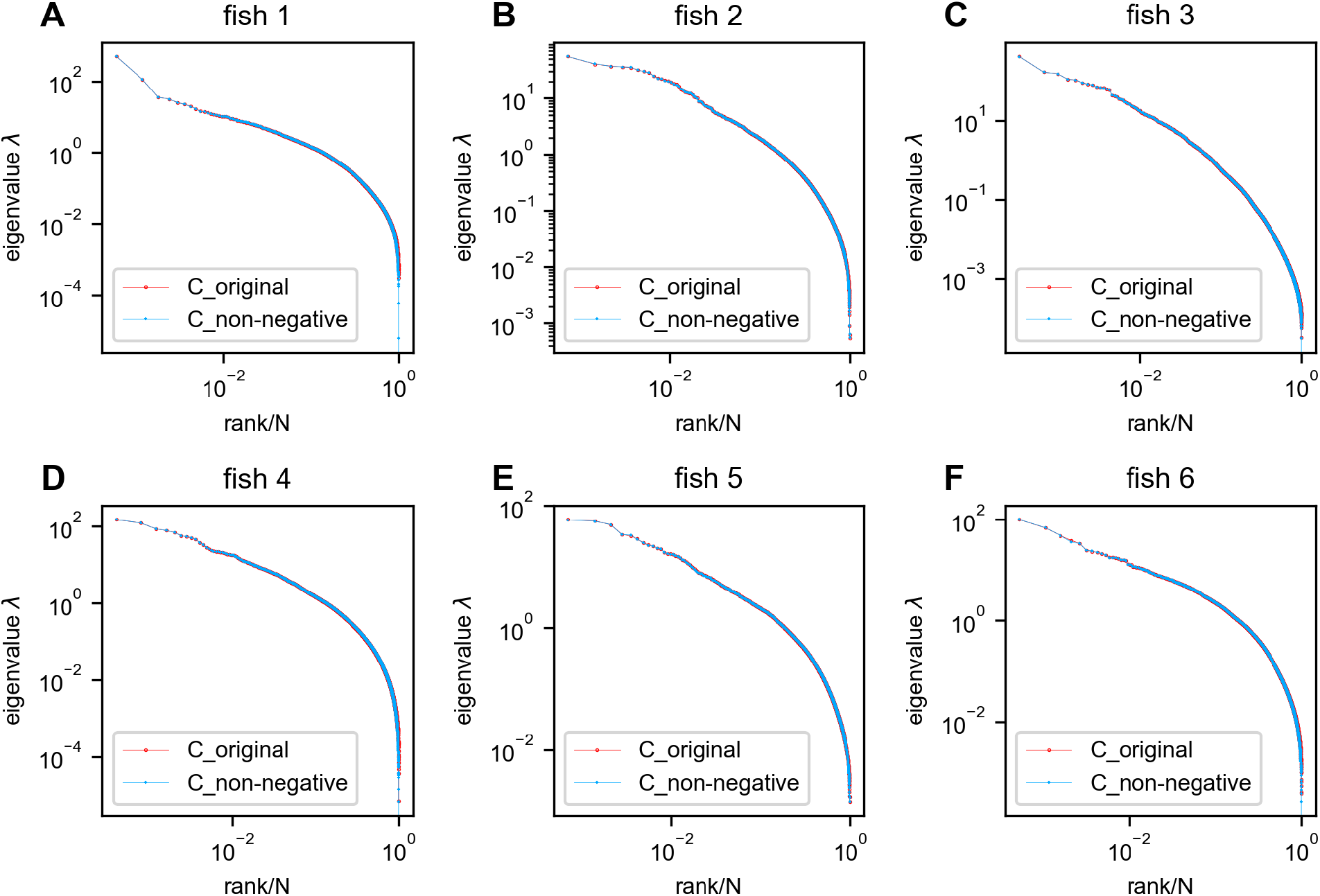
Negative covariances do not affect the eigenspectrum of the zebrafish data. Red: eigenspectrum of the original data covariance matrix. Blue: eigenspectrum of the covariance matrix with negative entries replaced by zeros. In this figure, all neurons recorded in each fish were utilized without any sampling.

**Figure S19.**
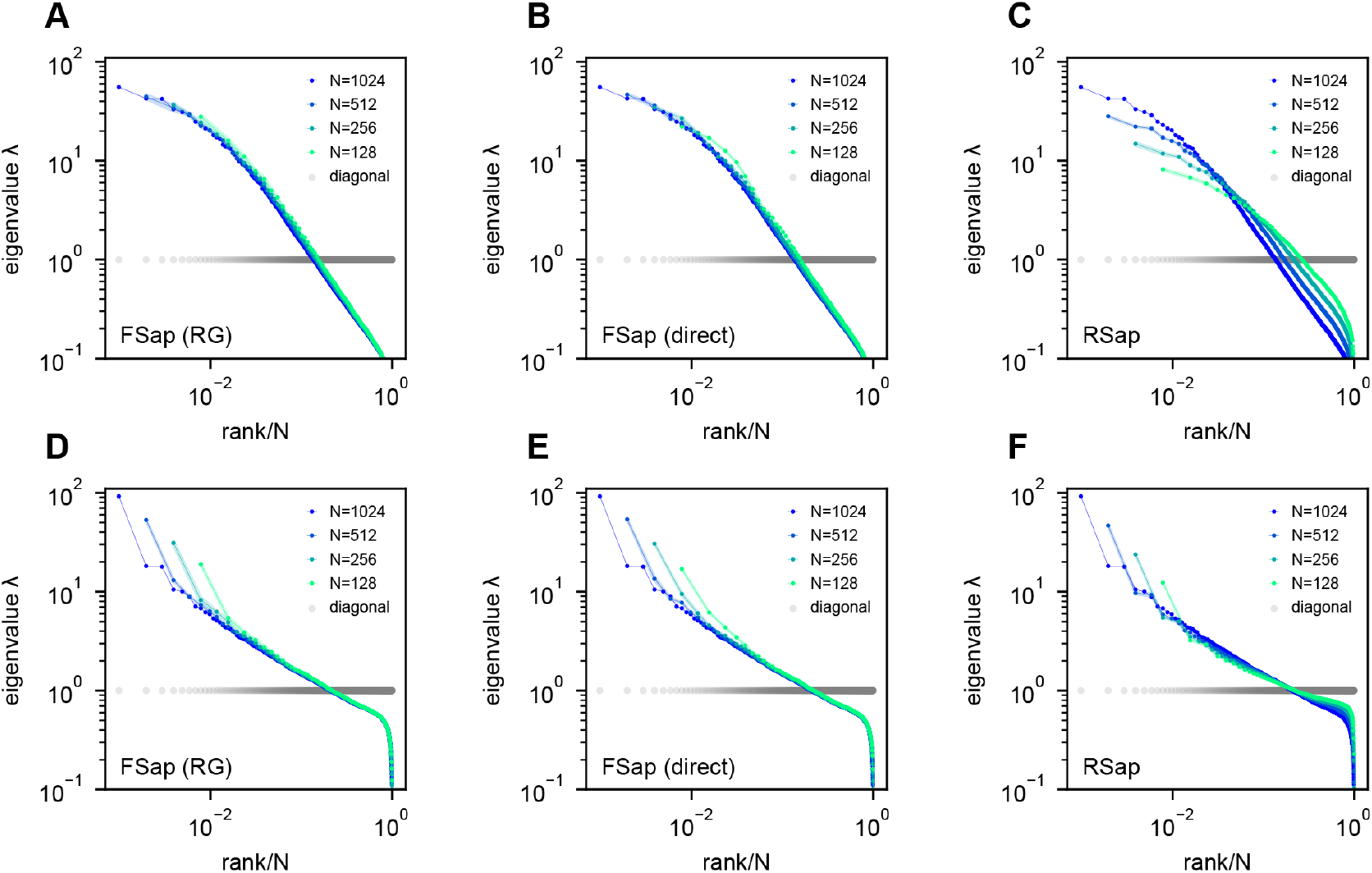
Eigenspectra of RG-inspired clustering, direct functional region sampling (FSap), and random sampling (RSap) in ERM. **A,D**. Renormalization-Group (RG) clustered eigenspectra of ERM. The size of the cluster is denoted by *N*, which is the number of neurons in each cluster. We adopt the RG approach (20, 67), but with a specific modification (Methods). **B,E**. Direct spatial sampling in the functional space (FSap) and the corresponding ERM eigenspectra. We began our analysis with a set of *N*_0_ neurons distributed in the functional space. Initially, we chose *N* = *N*_0_/2 neurons that were located exclusively on one side of the x-axis of this space. We then proceeded to select *N* = *N*_0_/4 neurons from 4 quadrants. This sampling process was repeated iteratively, generating successively smaller subsets of neurons. **C,F**. Random sampled (RSap) eigenspectra of ERM. ERM parameters: **A-C** Exponential function 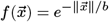 where *b* = 1, *ρ* = 10.24 and dimension *d* = 2. **D-F** Approximate power law Eq. (11) with *μ* = 0.5, *ρ* = 10.24 and dimension *d* = 2. Other parameters are the same as Fig. 3. The standard error of the mean (SEM) across the clusters is represented by the shaded area of each line.

**Figure S20.**
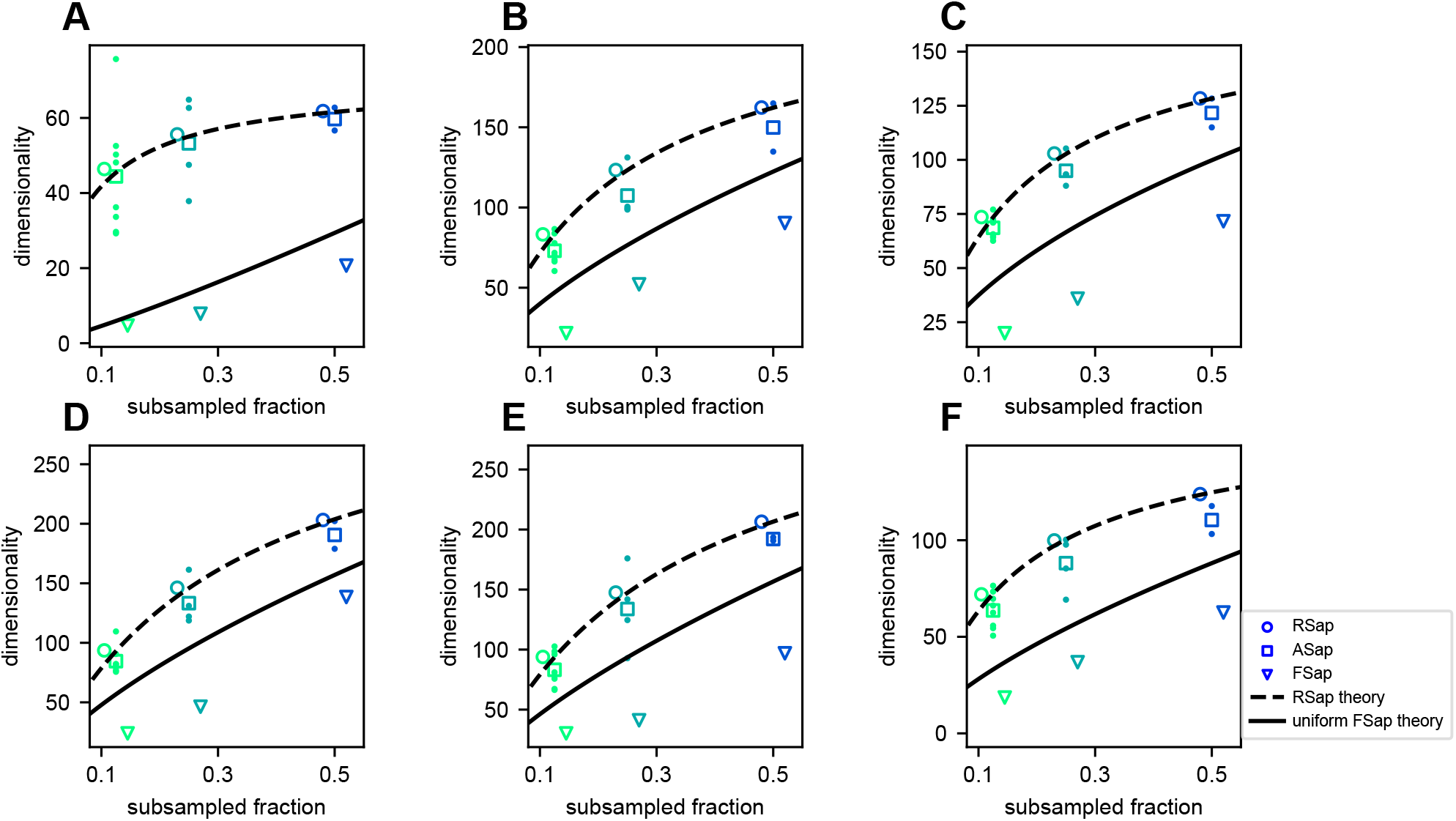
Dimensionality (*D*_PR_) across sampling methods in fish data. **A-F** Result from fish 1 to fish 6: mean RSap *D*_PR_ (circles), mean (squares) and individual ASap *D*_PR_, and FSap’s most correlated cluster *D*_PR_ (triangles). Dashed and solid lines indicate RSap and uniform FSap theoretical predictions, respectively. ERM parameter: *μ* = 0.6, *d* = 2, functional coordinates follow a multivariate normal distribution with variance 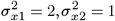, anatomical coordinates follow a multivariate normal distribution with variance 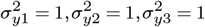.

**Figure S21.**
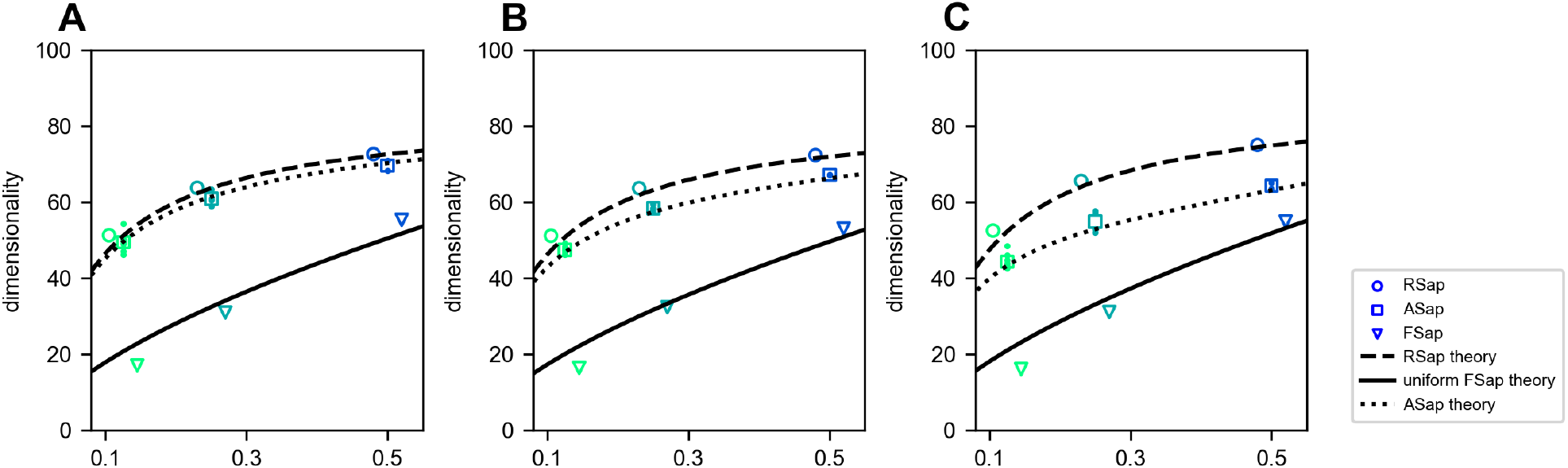
Dimensionality (*D*_PR_) across sampling methods in ERM. PR dimensionality result of ERM model, coordinate in funcitonal and anatomical space are multivariate Gaussian distribution, the CCA correlation between funcitonal and anatomical space are *RCCA* = 0.4, 0.6, 0.8 in **A-C**. Mean RSap *D*_PR_ (circles), mean (squares) and individual ASap *D*_PR_, and FSap’s most correlated cluster *D*_PR_ (triangles). Dashed and solid lines indicate RSap and uniform FSap theoretical predictions, 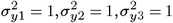.

**Figure S22.**
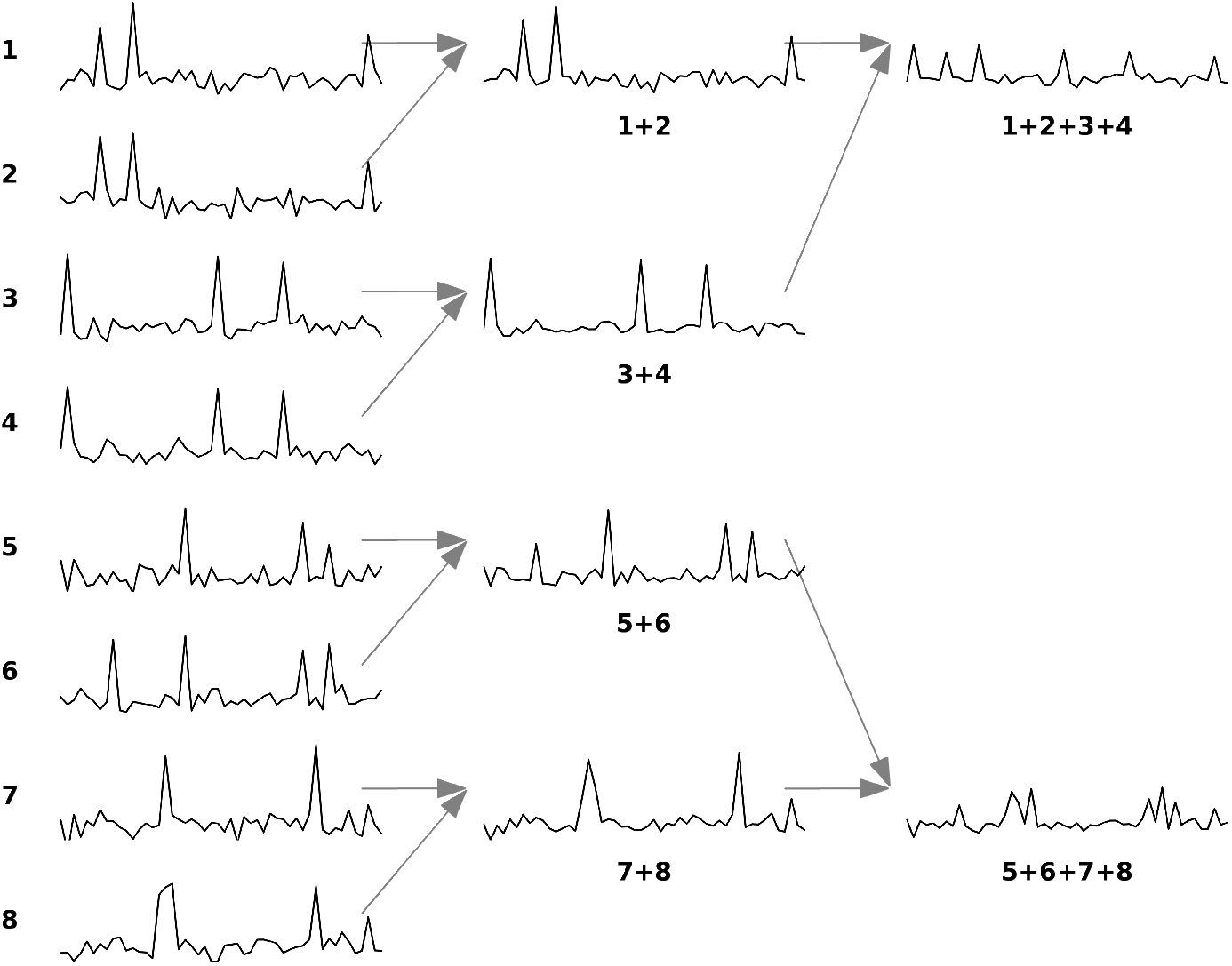
Example of Renormalization Group (RG) approach for a set of eight neurons. The figure is adapted from (20). The diagram illustrates the iterative clustering process for eight neurons. In each iteration, neurons are paired based on maximum correlation, with their activities combined through summation and normalized to maintain unit mean for nonzero values. Each neuron can only be paired once per iteration, ensuring all neurons are grouped by the iteration’s end.

**Figure S23.**
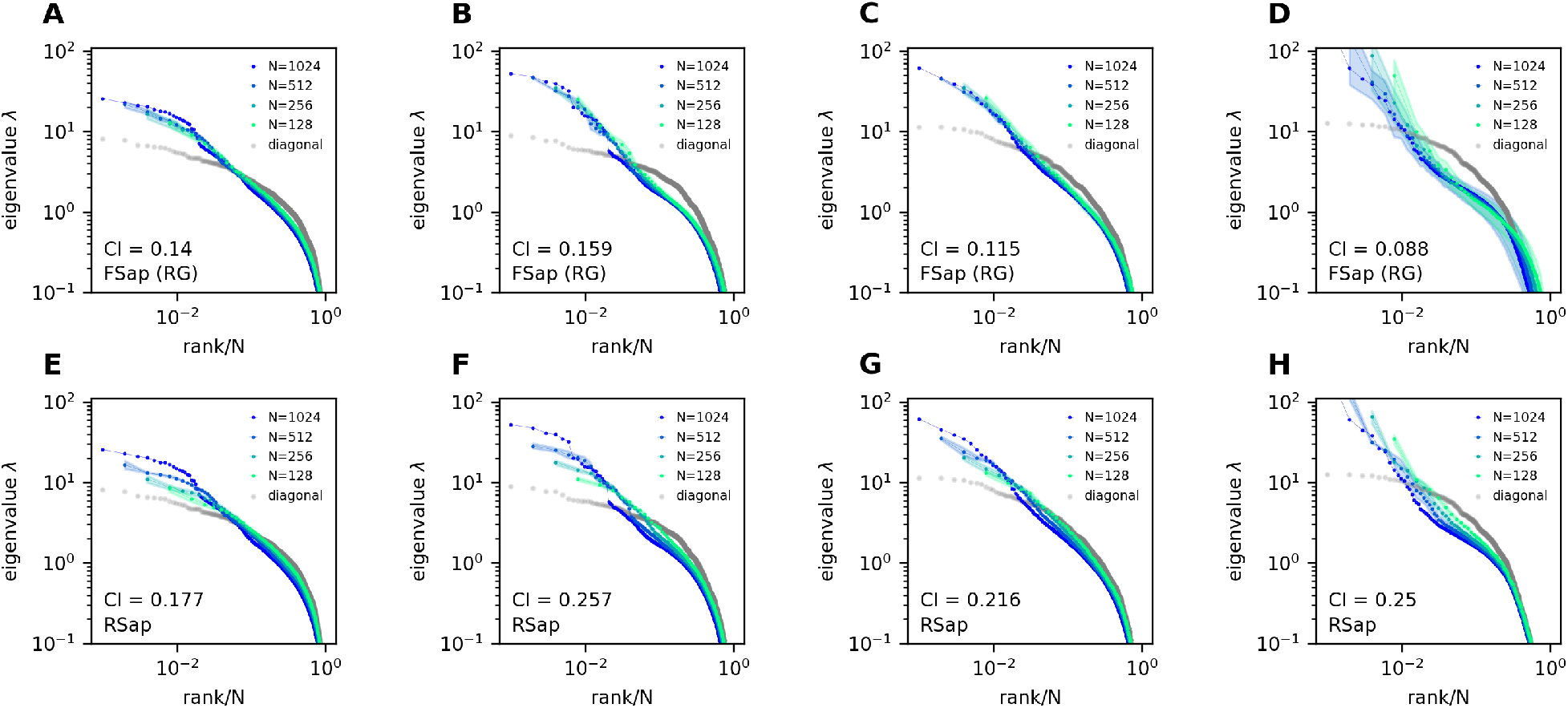
Morrell et al.’s latent variable model. **A-D**: Functional sampled (FSap) eigenspectra of the Morrell et al. model. **E-H**: Random sampled (RSap) eigenspectra of the same model. Briefly, in Morrell et al.’s latent variable model (36, 52), neural activity is driven by *Nf* latent fields and a place field. The latent fields are modeled as Ornstein-Uhlenbeck processes with a time constant *τ*. The parameters *ϵ* and *η* control the mean and variance of individual neurons’ firing rates, respectively. The following are the parameter values used. **A,E**: Using the same parameters as in (52): *Nf* = 10, *ϵ* = −2.67, *η* = 6, *τ* = 0.1. Half of the cells are also coupled to the place field. **B,C,D,F,G,H**: Using parameters from (36): *Nf* = 5, *ϵ* = −3, *η* = 4. There is no place field. The time constant *τ* = 0.1, 1, 10 for **B,F, C,G**, and **D,H**, respectively.

**Figure S24.**
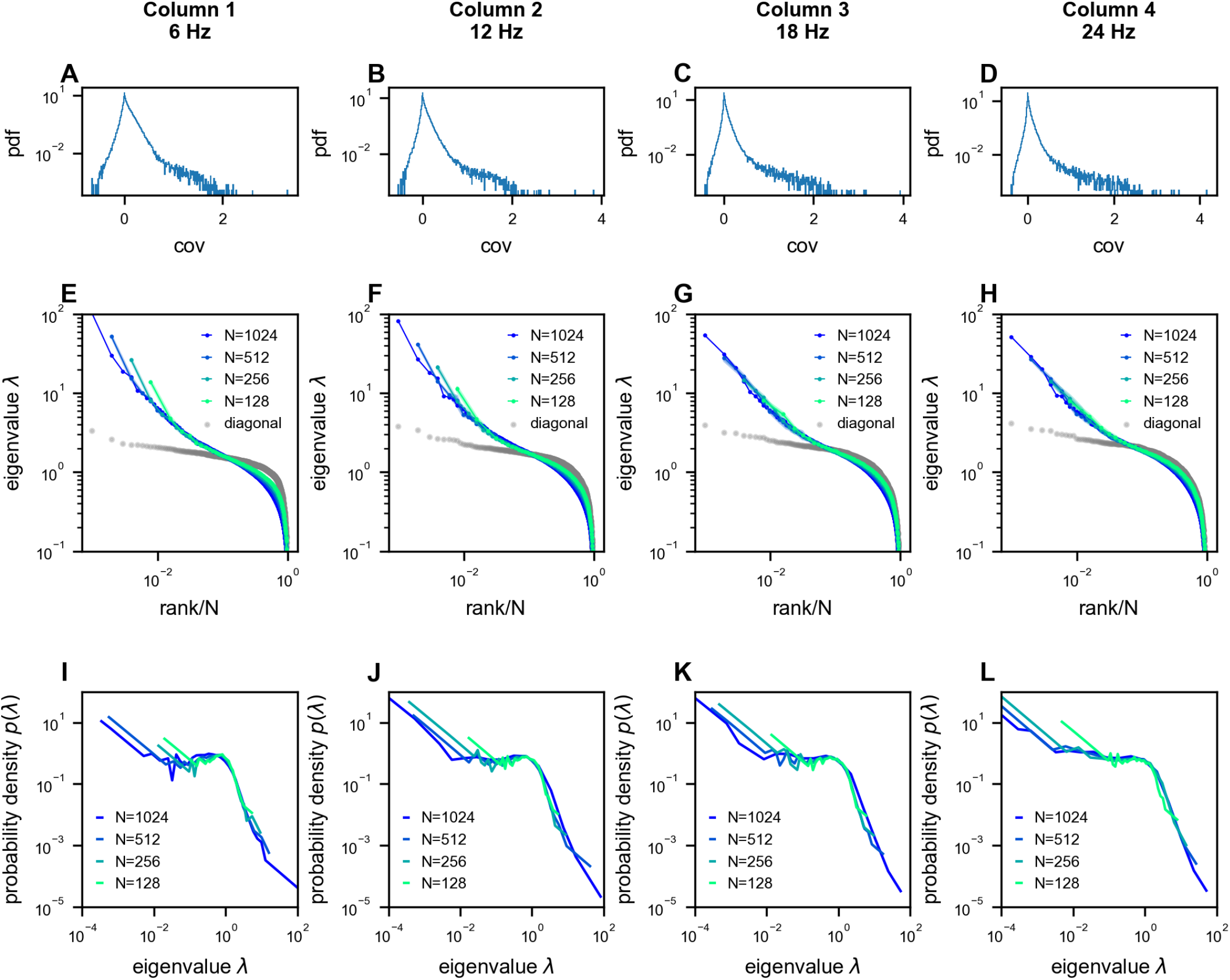
Scale-invariant properties persist across different temporal sampling rates in neural recordings. Analysis of multi-area Neuropixels recordings (23) from 1024 neurons, downsampled to different rates resulting in 7200 time frames per condition (6 Hz, 12 Hz, 18 Hz, and 24 Hz; columns 1-4 respectively). **A-D**. Distribution of pairwise covariances after normalization to unit variance (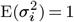, see Methods). **E-H**. Eigenvalue spectra of the covariance matrices, showing similar power-law scaling across sampling rates. **I-L**. Probability density functions (PDFs) of the eigenvalues, demonstrating that the characteristic shape of the distribution is preserved across different temporal resolutions.

## 6 Supplementary note

In this appendix, we elaborate upon the sketch introduced in the Methods, and present a full derivation of the covariance eigenspectrum of our ERM model, This section is organized as follows. First, we will briefly introduce the relationship between the eigenvalue probability density distribution and the resolvent. Second, we will turn the problem of calculating the resolvent to a calculation of the partition function using a field-theoretic representation and proceed to manipulate the partition function using the replica method. Third, we will introduce two approximate methods for calculating the partition function, leading to the high-density theory and the Gaussian variational method. We will discuss the implications and predictions of each method. Finally, we will discuss the relationship between the two methods and identify the parameter regime where the high-density theory agrees with the numerical simulation.

**Table S4.** Table of notations.

| Notation | Description |
| --- | --- |
| $g(z)$ | resolvent, Eq. (S2) |
| $\langle \dots \rangle$ | the average across realizations of $C$ (i.e., random $\vec{x}_i$'s and $\sigma_i^2$'s), Eq. (S1) |
| $\Xi(z)$ | Canonical partition function, Gaussian integral representation of the determinant $[\det(z - C)]^{-1/2}$, Eq. (S5) |
| $\phi$ | intermediate variable for Gaussian integral representation $\Xi(z)$, Eq. (S5) |
| $\psi$ | density field of $\phi$ |
| $\hat{\psi}$ | respective Lagrange multiplier fields of $\psi$ |
| $S_1$ | the action in $\Xi(z)$ (by analogy with the path integral formulation of quantum mechanics) |
| $S_h$ | the action in the high-density approximation of $\Xi(z)$ |
| $S_v$ | the action in the variational approximation of $\Xi(z)$ |
| $A$ | term in $S_1$ |
| $f^{-1}$ | the operator inverse of $f$, Eq. (S23) |
| $G$ | quadratic kernel in the Gaussian integral approximation of $\Xi(z)$ |
| $G^{-1}$ | the operator inverse of $G$, same definition as $f^{-1}$ |
| $\tilde{G}$ | the Fourier transform of $G$ |

### 6.1 Resolvent

The eigenvalues λ_*n*_ of a Hermitian matrix *C* are real. Their probability density function or eigendensity is formally given by

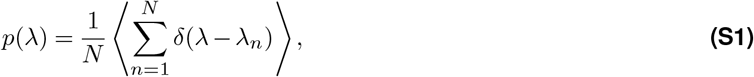

where ⟨… ⟩represents an average across different realizations of *C*. The eigendensity is connected with the resolvent (34, 35)

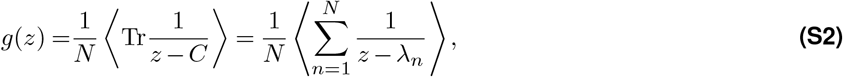

we therefore compute the eigendensity using the standard inverse formula of Stieltjes tranform:

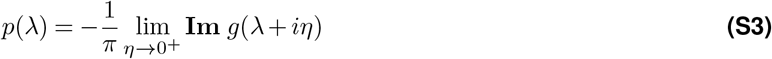

### 6.2 Field representation

In this section, we discuss a field-theoretical representation of the resolvent *g*(*z*). First, we rewrite Eq. (S2) as

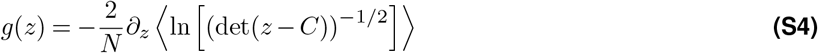

The determinant (det(*z* −*C*))^−1/2^ can be represented as a Gaussian integral

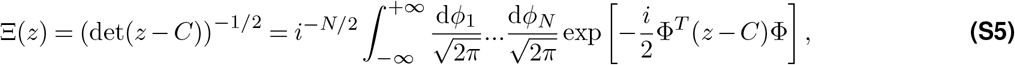

where Φ = [ϕ_1_, …, *ϕ*_*N*_]^*T*^, and 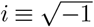.

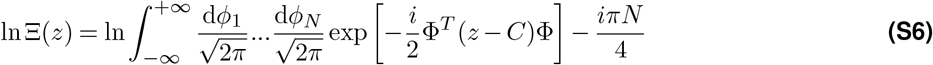

We thus establish a relationship between the resolvent and Ξ

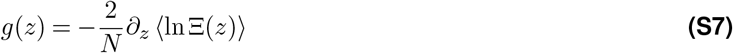

Note that the constant term in Eq. (S6) can be killed by ∂_*z*_ and we will ignore it in the sequel. Eq. (S7) is the central formula in this note. Ξ(*z*) is also called the partition function in statistical physics. We endeavor to find a way to compute the average of ln Ξ(*z*).

Recall that in our ERM model (Result Eq. (2) and Fig. 3A), the covariance between neuron *i* and neuron *j* is determined by the distance kernel function and their neural activity variances:

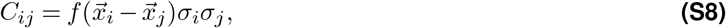

where 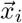 are sampled from a uniform coordinate distribution 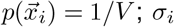 are i.i.d. chosen from a probability density distribution *p*(*σ*) and are independent of the neuron coordinates 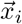. The ⟨*…* ⟩ in Eq. (S7) is therefore an average over all possible 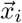 and *σ*_*i*_.

In order to compute ⟨ln Ξ(*z*)⟩, we apply the replica method based on a smart use of the identity

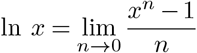

Eq. (S7) now becomes

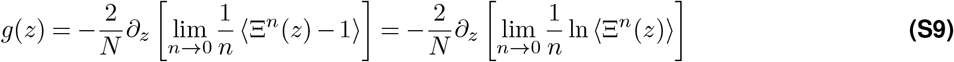

The idea is to compute the right-hand side for finite and integer *n* and then perform the analytic continuation to *n* → 0. Now we seek to determine the value of ⟨Ξ^*n*^(*z*)⟩. It contains *n* copies (replicas) of the original system

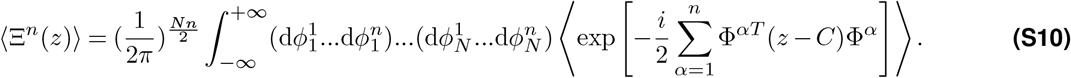

Writing it down explicitly, we have

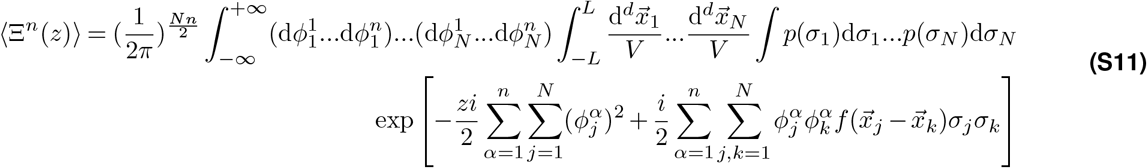

In order to proceed further, we introduce the following auxiliary fields :

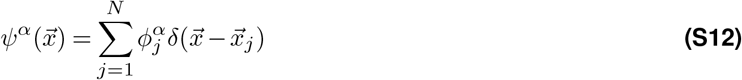

Eq. (S12) can be represented as a following functional integral

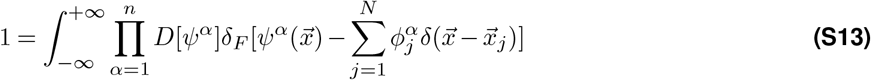

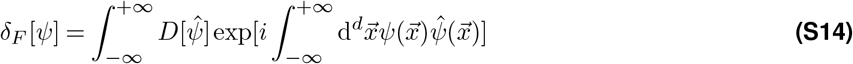

or we can combine Eq. (S13) and Eq. (S14) as

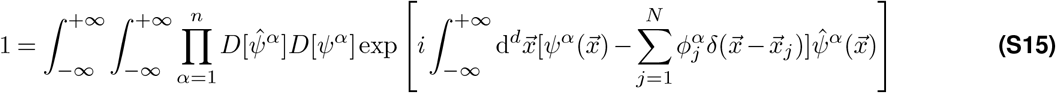

Using Eq. (S12), we can write the term 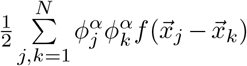 in Eq. (S11) as

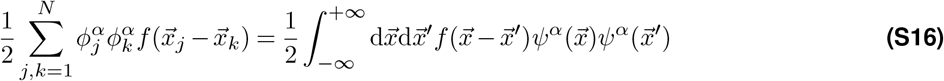

We insert the relation Eq. (S15) and Eq. (S16) into Eq. (S11),

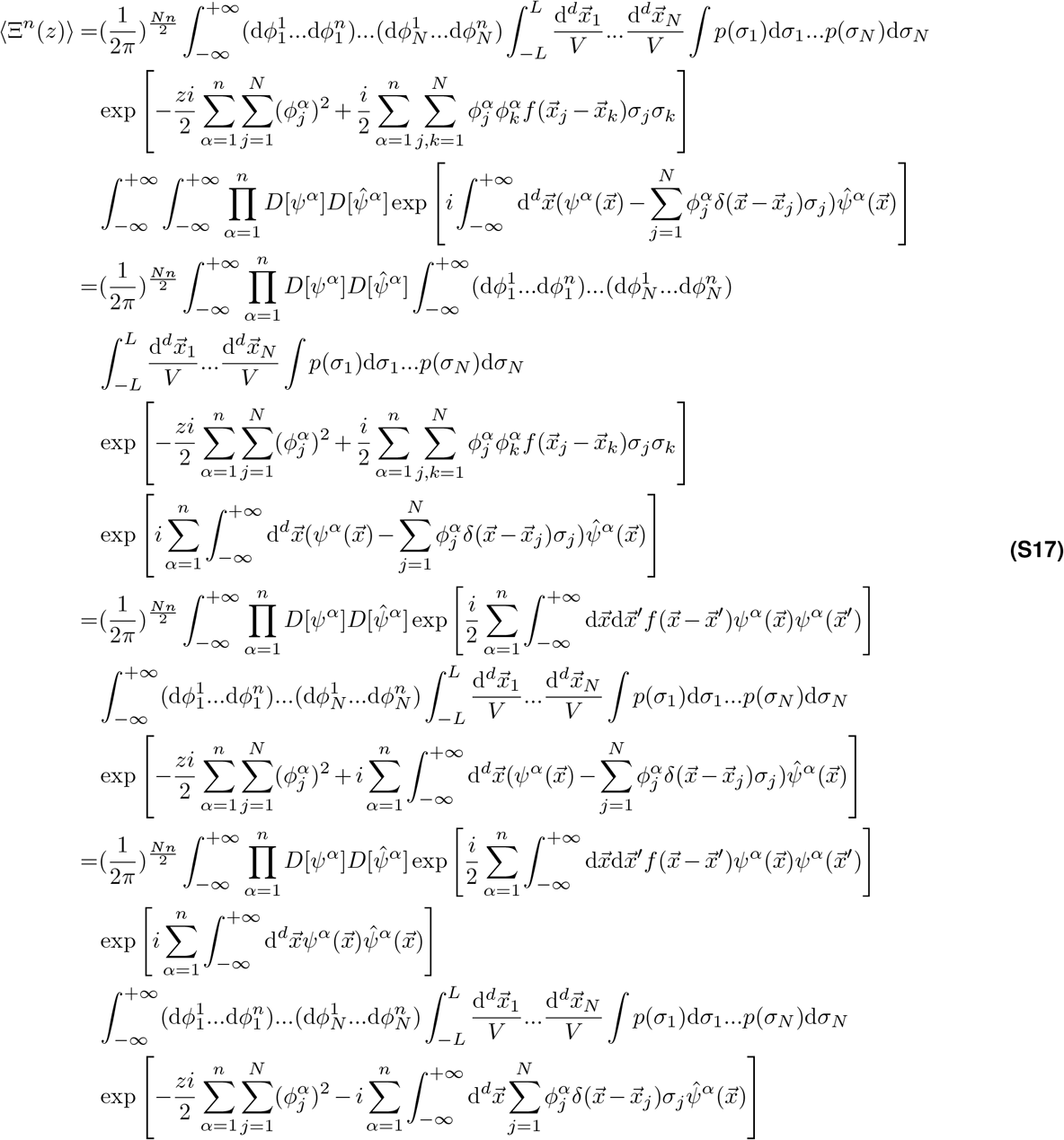

Integrating the last term in Eq. (S17)

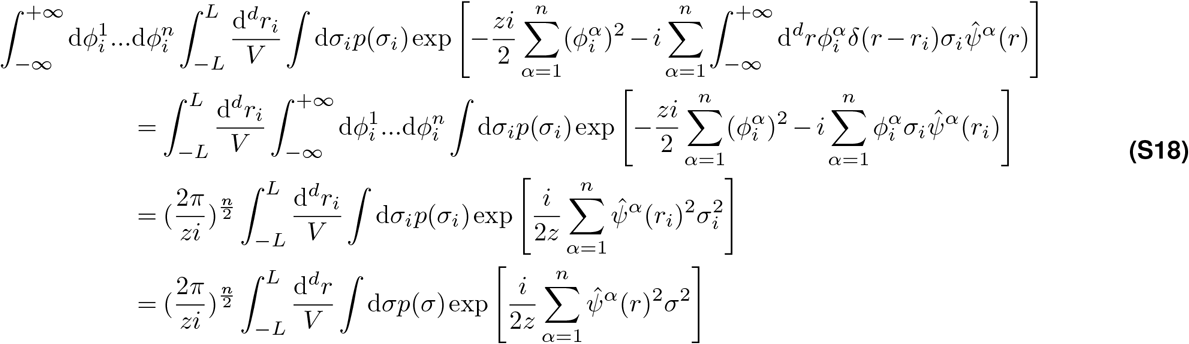

so that ⟨Ξ^*n*^(*z*)⟩ from Eq. (S11) can be written as

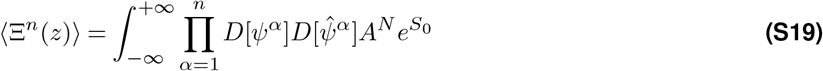

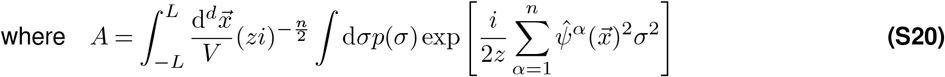

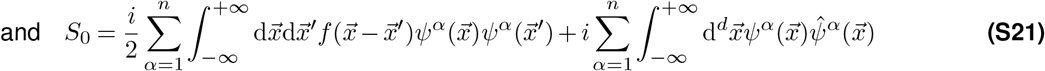

Integrating out the *ψ*^*α*^ in ⟨Ξ^*n*^(*z*)⟩ Equations (S19) and (S21)

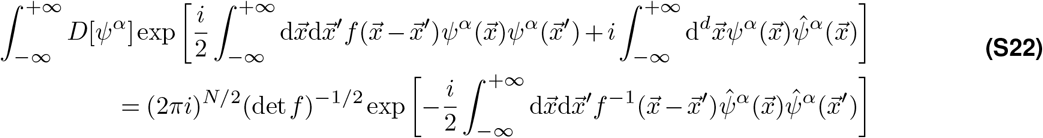

Here *f*^−1^ is the inverse kernel satisfying:

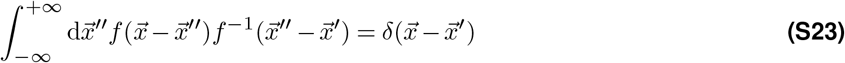

so that ⟨Ξ^*n*^(*z*)⟩ can be written as

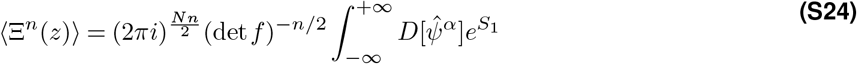

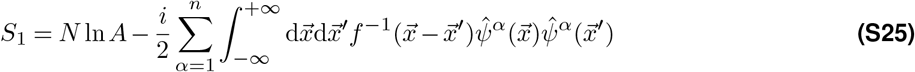

The constant term 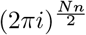 of ⟨Ξ^*n*^(*z*)⟩ can be ignored because we should compute ∂_*z*_ ⟨ln Ξ(*z*)⟩ Eq. (S7) in the end.

To ensure the mathematical rigor in section 6.4, Eq. (S42), we next apply the Wick rotation 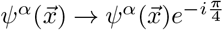 (section 6.7).

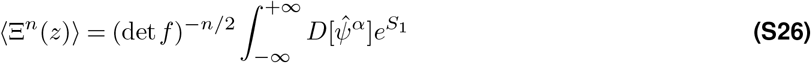

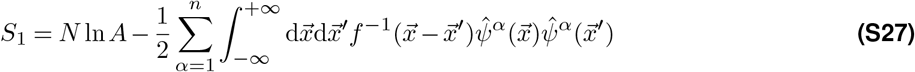

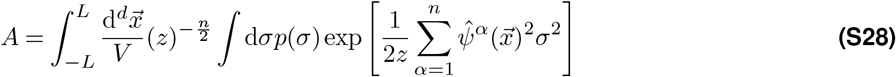

### 6.3 High-Density Expansion

In this section, we directly calculate the canonical partition function ⟨Ξ^*n*^(*z*) ⟩ in the *z* limit by approximating the term *N* ln *A* (Eq. (S27)) to a quadratic action, from which the partition function (Eq. (S26)) would become a Gaussian integral.

Let us first calculate the *A*^*N*^ in *z* → ∞ limit

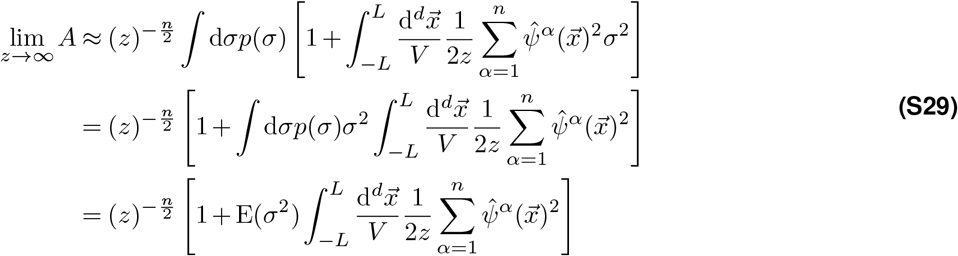

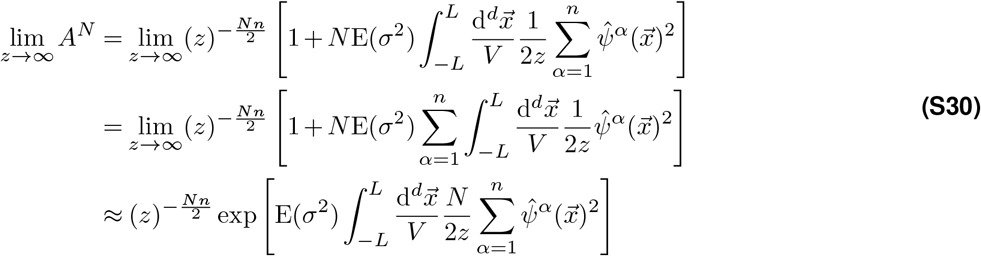

Now let us calculate ⟨Ξ^*n*^(*z*)⟩ (Equations (S26) to (S28)) by letting *L* → ∞

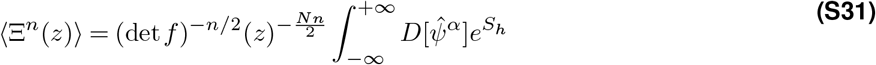

where the high-density quadratic action

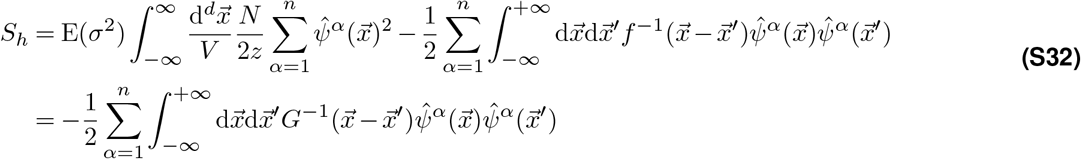

where 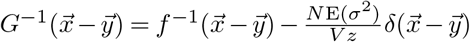. Next, by integrating out the 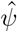 field, we find

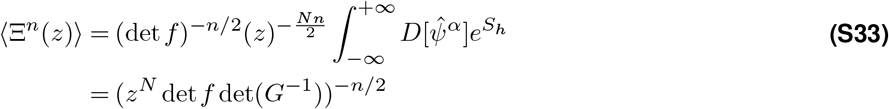

Using Eq. (S9) that connects the partition function with the resolvent, we have

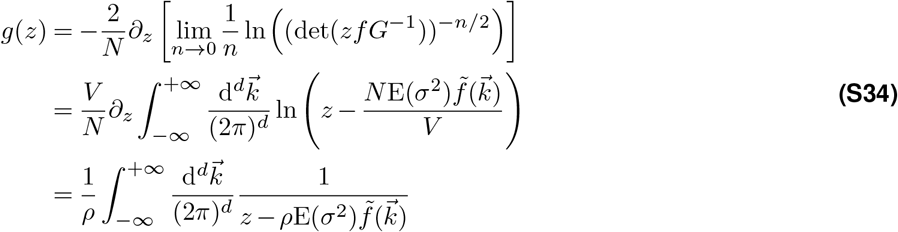

where 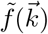 is the Fourier transform of 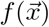.

Finally, the eigendensity *p*(*λ*) (Eq. (S3)) is given by

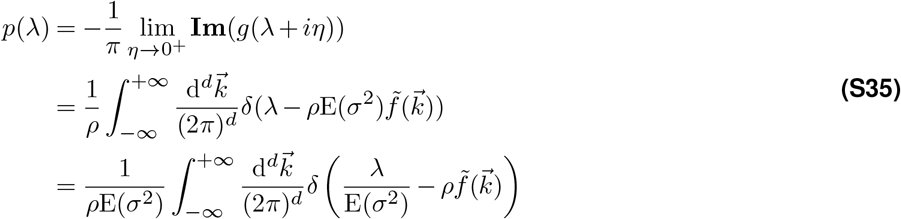

#### 6.3.1 Derivation of power-law eigenspectrum in high-density limit

Here we calculate the eigendensity of our model, with the kernel function 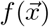 (table S3). The Eq. (S35) (set E(*σ*^2^) = 1 as in Result section 2.2) can be written as:

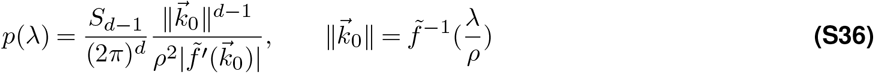

where *S*_*d*−1_ is the surface area of *d* − 1 dimensional sphere. Here we consider the approximation 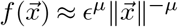, whose Fourier transform and its derivative are 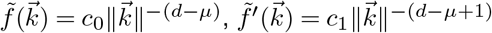 and 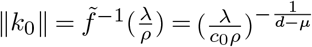. The constants are given by 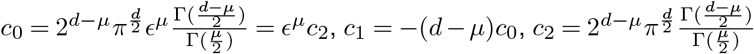

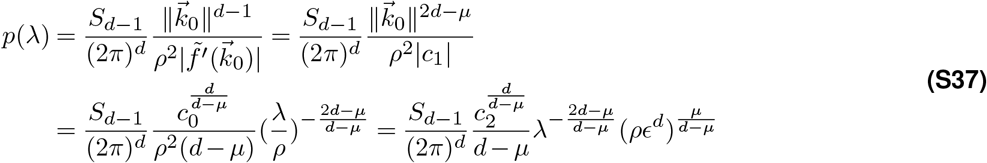

#### 6.3.2 Derivation of eigenspectrum with exponential kernel function in high-density limit

Here we consider the exponential kernel function 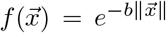, whose Fourier transform and its derivative are 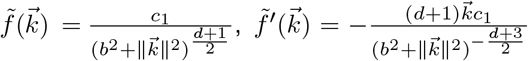 and 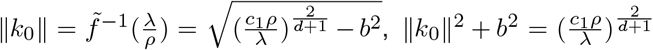, where 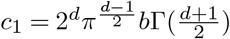.

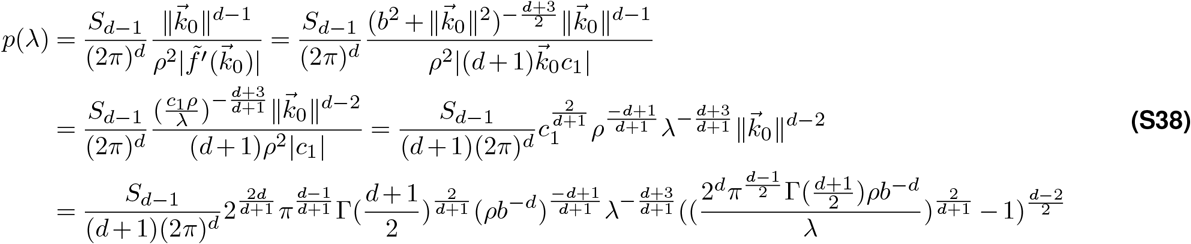

It is straightforward to see that this spectrum is not scale invariant. For example, when *d* = 2, the above expression reduces to a perfect power law spectrum 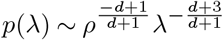, which changes with scale over sampling.

### 6.4 Variational Approximation

To find a general approximation for the eigenspectrum that goes beyond the high-density limit, we use Gaussian variational approximation in the field representation, namely by looking for the best quadratic action *S*_*v*_,

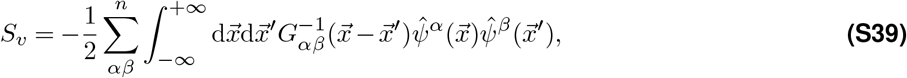

to approximate the action *S*_1_ in the partition function (Equations (S26) to (S28)). This enables us to represent the partition function by a Gaussian integral, which can be evaluated analytically. We find the best quadratic action *S*_*v*_ by minimizing the difference between *S*_1_ and *S*_*v*_, which is defined as KL divergence between two distributions that are proportional to 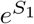 and 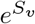.

In this section, we will proceed by using the grand canonical ensemble formulation, namely the average in Eq. (S1), instead of using a fixed covariance matrix size *N*, which is now carried out across all different sizes. If *N* follows a Poisson distribution, it is easy to show (section 6.8) that the grand canonical partition function is given by Eq. (S113):

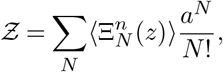

where *a* = ⟨*N*⟩. As a result, the new action *S*_1_ becomes

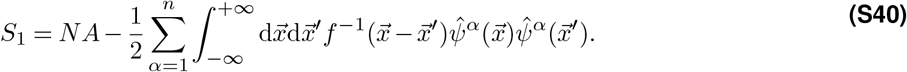

Here and below, *N* should be viewed as the average matrix size. The resolvent *g*(*z*) in Eq. (S9) can be similarly generalized to Eq. (S114),

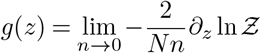

As in statistical physics, we define the free energy as

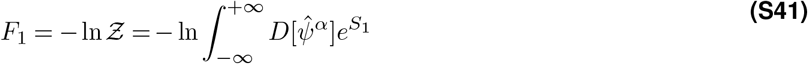

We shall define the variational free energy *F*_*v*_ such that it would approximate the true free energy *F*_1_ by minimizing *D*_*KL*_(*P*_*v*_||*P*_1_),

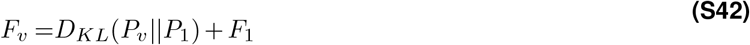

where

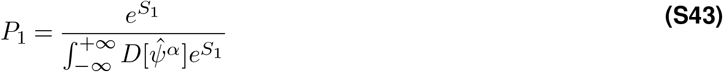

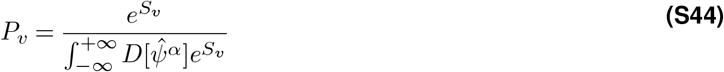

The KL divergence *D*_*KL*_(*P*_*v*_ *P*_1_) is always nonnegative and the free energy *F*_1_ is independent of the quadratic action *S*_*v*_. Therefore, we need to minimize the variational free energy *F*_*v*_. Let us now examine the variational free energy *F*_*v*_

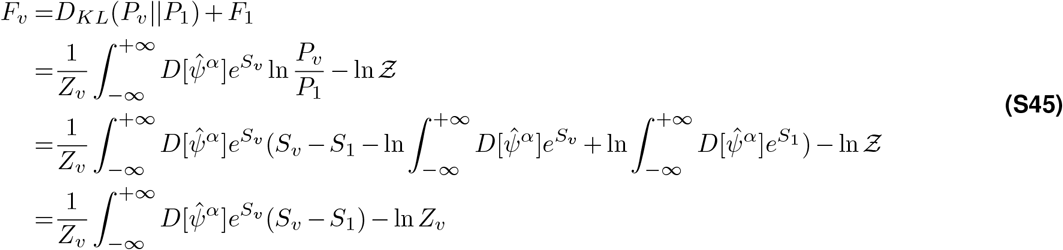

Here *Z*_*v*_ is the normalization factor

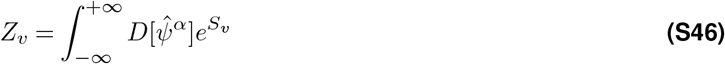

Since we want to minimize *F*_*v*_, the constant term

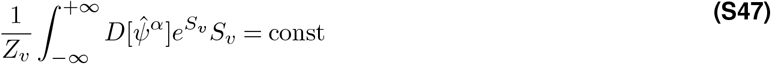

can be ignored, and Eq. (S45) is reduced to

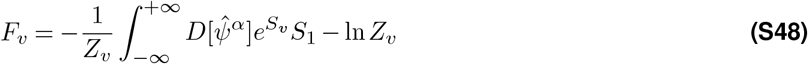

To simplify the formula, let us introduce *S*_2_

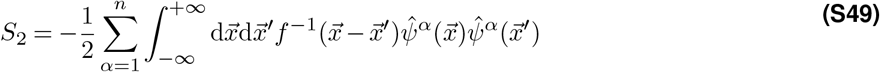

and rewrite Eq. (S48) as

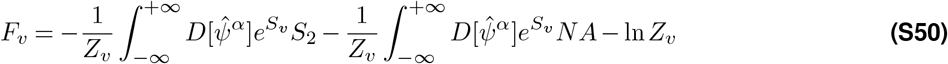

Next, we will compute each term in the variational free energy *F*_*v*_

First, we calculate the third term ln *Z*_*v*_ in Eq. (S50) by Equations (S39) and (S46)

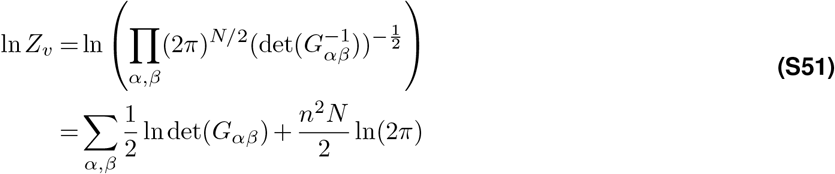

Second, we calculate the first term 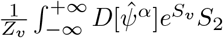 in Eq. (S50)

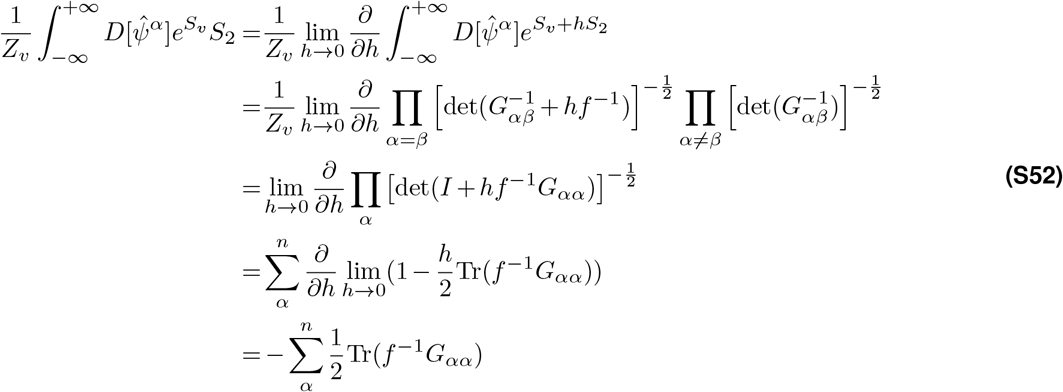

Third, we calculate the second term 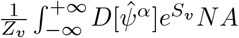 in Eq. (S50), recall the term *A* (Eq. (S28))

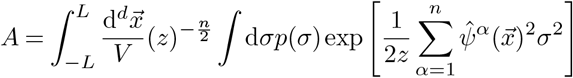

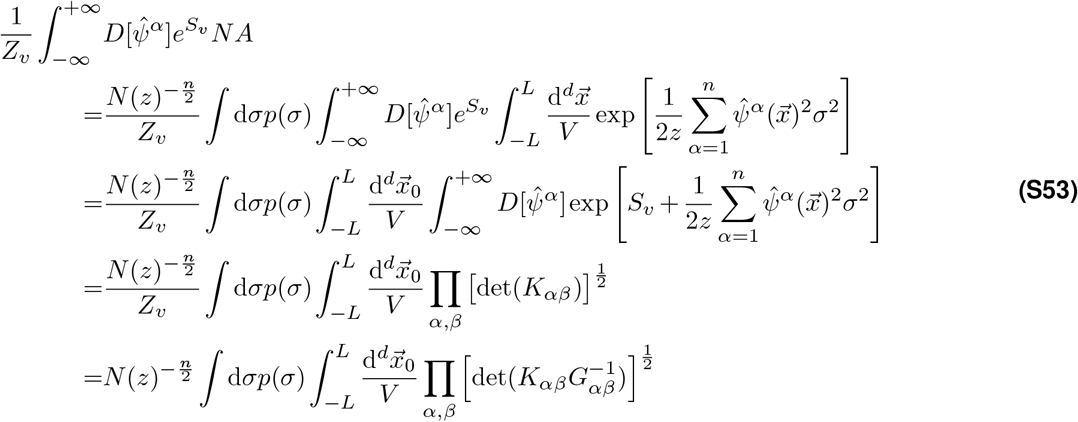

where

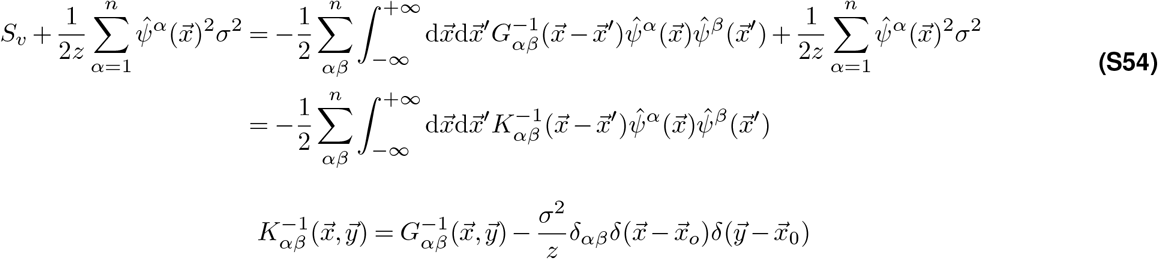

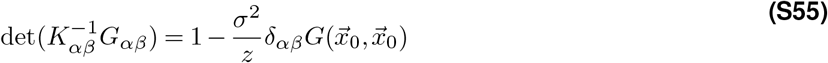

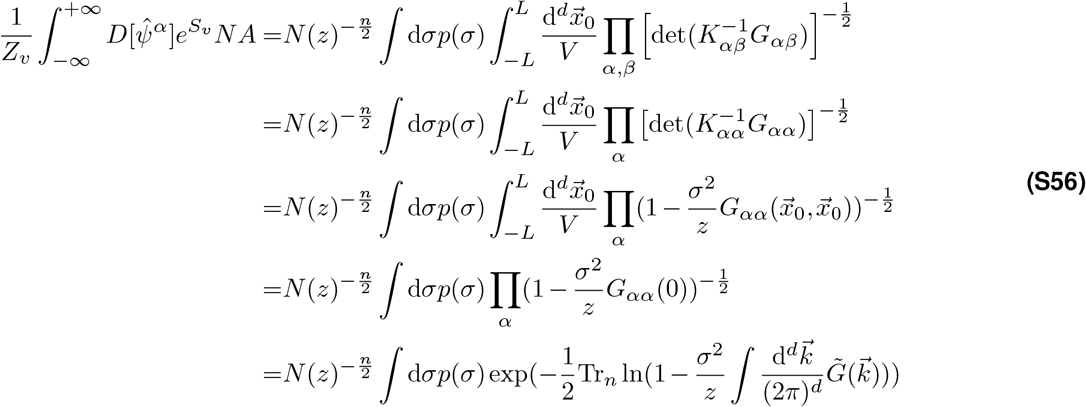

In sum, the variational free energy *F*_*v*_ is equal to

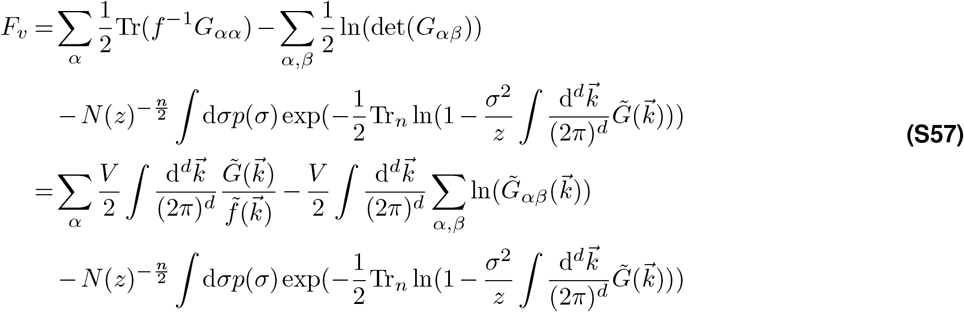

Now let us find the best quadratic action *S*_*v*_ that minimizes the variational free energy *F*_*v*_

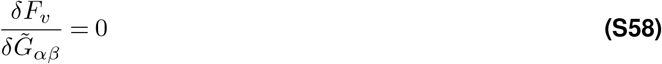

The solution of Eq. (S58) is given by

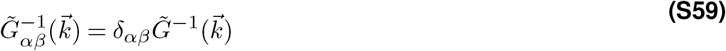

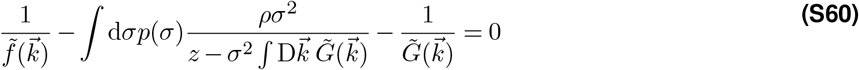

where 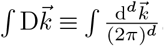. By using Eq. (S114)), we finally obtain

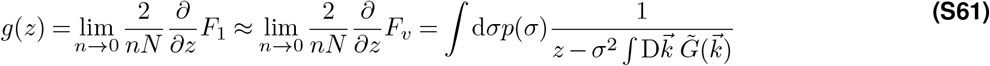

### 6.5 Scale invariance of the covariance spectrum in the Gaussian variational Model

In section 2.2 (Result), we point to two factors that contribute to the scale-invariance of eigenspectrum using the high-density theory. In this section, we show that the same conclusion can be drawn by using the Gaussian variational method. Furthermore, we examine how the heterogeneity of neural activity influences the eigendensity calculated by the Gaussian variational model. We show that 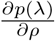, which characterizes the change of eigendensity due to sampling in the functional space, decreases with the heterogeneity of neural activity described by higher-order moment of neural activity variance, e.g., E(*σ*^4^).

Let us rewrite Eq. (S60) as

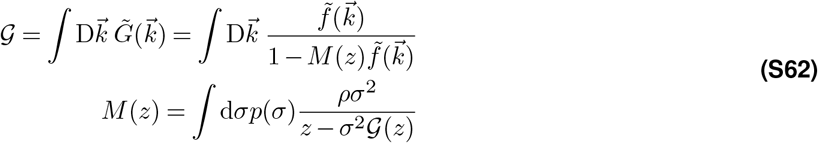

To present a formal expression for the eigendensity, let us define **Re**(𝒢) ≡ *g*_*r*_, **Im**(𝒢) *g*_*i*_. From Equations (S3) and (S61), we find

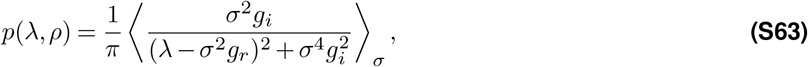

where ⟨*…*⟩_*σ*_ = ∫ *…p*(*σ*)d*σ*.

A direct computation of Eq. (S63), however, remains difficult: the complication arises from the complex function *M*(*z*) in Eq. (S62), which in turn is an integral function of 𝒢. To streamline the calculation, let us further define **Re**(*M*) ≡ *ρa*, **Im**(*M*) ≡ *ρb*. Writing it down explicitly, we have

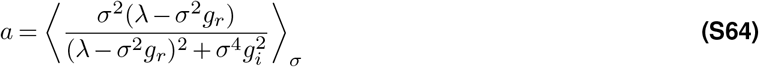

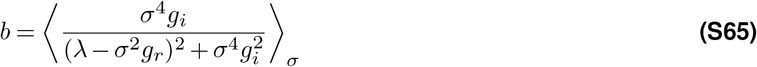

The real and imaginary part of 𝒢 can now be expressed as functions of *a* and *b*. Integrating Eq. (S62) in the spherical coordinates, we have

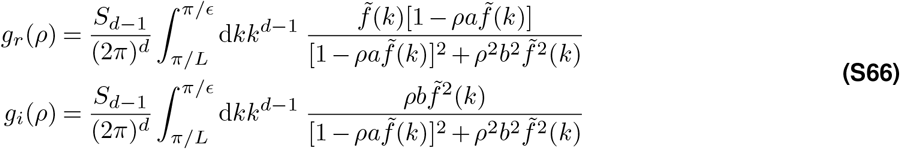

where for clarity, we have abused the notation a bit by defining 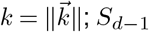 is the surface area of unit *d*-ball in the momentum space. In order to evaluate the integrals analytically, we introduce an ultraviolet cutoff *π/ϵ*. Numerically, whether integrating up to *π/ϵ* or greater than this bound shows little difference.

#### 6.5.1 Numerical solution of the Gaussian variational method

With Equations (S63) to (S66), we numerically calculate the eigendensity iteratively from the following steps:

- Step 1: set the initial values of *a* and *b* as *a*_0_ = 1, *b*_0_ = 1
- Step 2: solve for *a* in Eq. (S64) with fixed *b*
- Step 3: solve for *b* in Eq. (S65) with fixed *a*
- Step 4: iterate Step 2 and Step 3 10 times
- Step 5: calculate *p*(*λ*) using Eq. (S63)

Note that we plug Eq. (S66) into Equations (S64) and (S65) in step 2-3.

#### 6.5.2 Two contributing factors on the scale invariance

We next derive an analytical expression for Eq. (S66) by considering the approximate power law kernel function 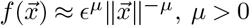, from which the high-density theory results on the scale invariance can be extended.

By a change of variable 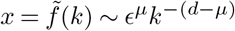, and let 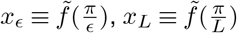, we have

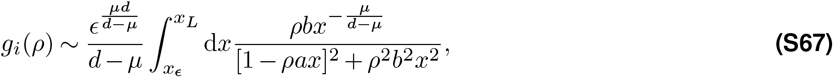

where ∽ indicates that all constant numerical factors (e.g., *π* and Γ(*d/*2)) are ignored. To compute Eq. (S67), we perform a branch cut at [0,], and perform a contour integral on the complex plane following the path in Fig. S25A. When 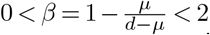, the integral on the large circle Γ_*R*_ and the small circle Γ_*ϵ*_ goes to zero as *x*_*L*_ ⟶ ∞ *x*_*ϵ*_ ⟶0, leaving only two simple poles (zeros of the function in the denominator) in the complex plane. By applying the residue theorem, we find an expression for *g*_*i*_ in the limit *L* → ∞, *ϵ* → 0

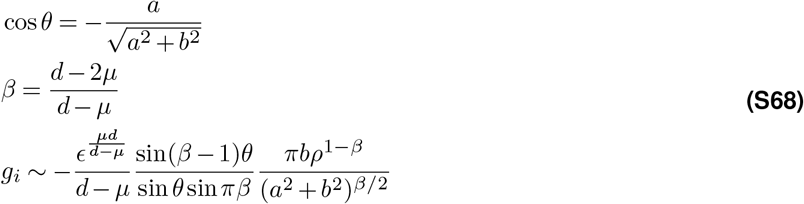

The analytical expression for *g*_*r*_ is a bit more involving.

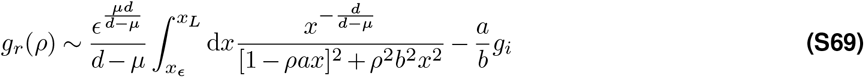

It has two terms, the second term is similar to Eq. (S67); the first term, however, diverges as *x*_*ϵ*_ ⟶ 0. Thus, the radius of the small circle Γ_*ϵ*_ in Fig. S25A cannot shrink to zero: this is precisely the requirement of an ultraviolet cutoff in the wave vector 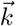 The contour integral on the large circle Γ_*R*_, on the other hand, goes to zero as *x*_*L*_ ⟶ ∞. Thus, the integral on Γ_*ϵ*_ contributes to the final result. By considering leading order term of *x*_*ϵ*_ for finite but small *x*_*ϵ*_, we find

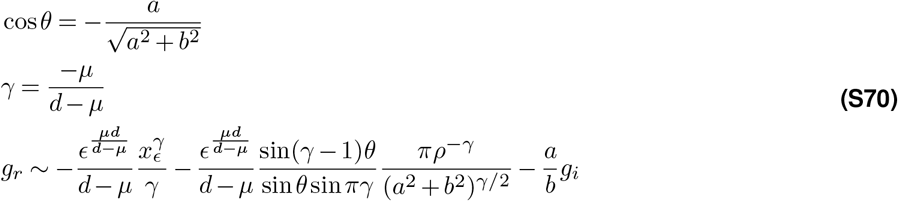

Recall 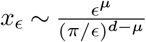, and we find that the first term in *g*_*r*_ is proportional to *π*^*µ*^*/µ*, independent of *ϵ*.

**Figure S25.**
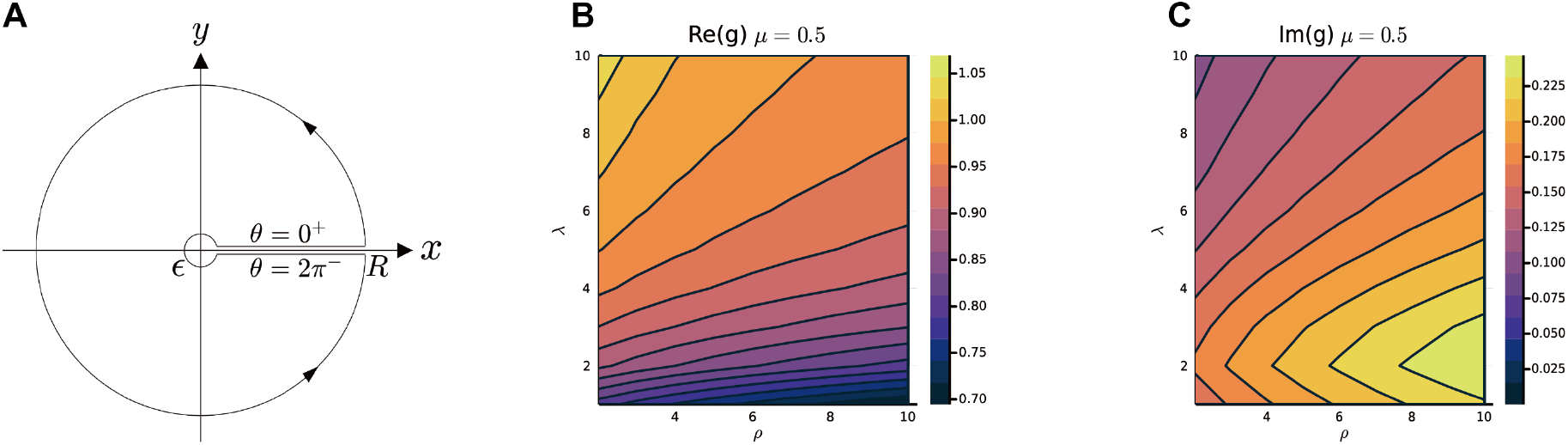
Calculate *g*_*i*_ and *g*_*r*_. **A**. The path of the contour integral for *g*_*i*_, *g*_*r*_ (Eq. (S67)). **B-C**. The heatmap of *g*_*r*_ and *g*_*i*_ with respect to *λ* and *ρ. g*_*i*_, *g*_*r*_ in B, C are calculated by the numerical method (section 6.5.1). The parameters are *N* = 1024, *ρ* = 10.24, *d* = 2, *L* = 10, *µ* = 0.5, *ϵ* = 0.03125. 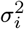 is i.i.d. sampled from a log-normal distribution with zero mean and a standard deviation of 0.5 in the natural logarithm of the 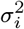 values; we also normalize 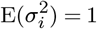.

According to Equations (S68) and (S70), one can immediately see that as *µ/d* ⟶ 0, the *ρ*-dependence relationship vanishes for *g*_*r*_ and *g*_*i*_. We therefore conclude that a slower power-law decay in the kernel function and/or a higher dimension of the functional space are two contributing factors for the scale-invariance of the covariance spectrum.

#### 6.5.3 Heterogeneity of neural activity across neurons enhances scale invariance

Next, we take a more close look at how the eigendensity changes with *ρ* for finite but small *µ/d* and when *λ* ≫ 1. Using Eq. (S63), we have

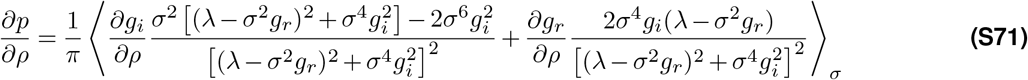

From numerical calculation, we find that typically *g*_*r*_ ≫ *g*_*i*_, so one can use the approximation

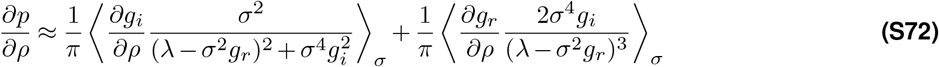

Recall Eq. (S63)

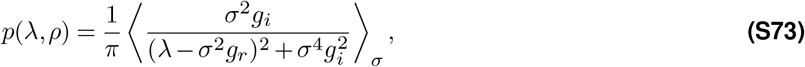

Since *p*(*λ, ρ*) is very small for large *λ*, a more appropriate measure is to examine

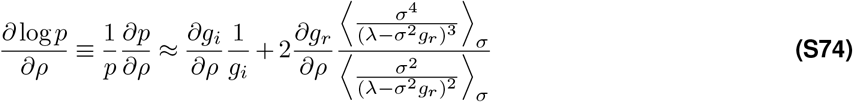

Considering the large eigenvalue case *λ σ*^2^*g*_*r*_ (the numerical value of *g*_*r*_ is on the order of 1), we perform Taylor expansion and arrive at

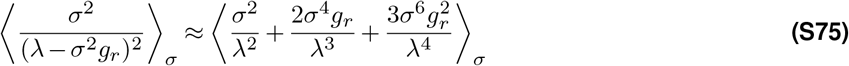

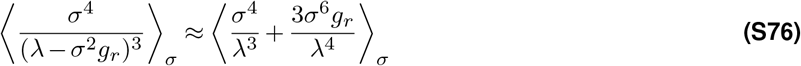

Note ⟨*σ*^2^⟩_*σ*_ ≡ E(*σ*^2^) is normalized to 1.

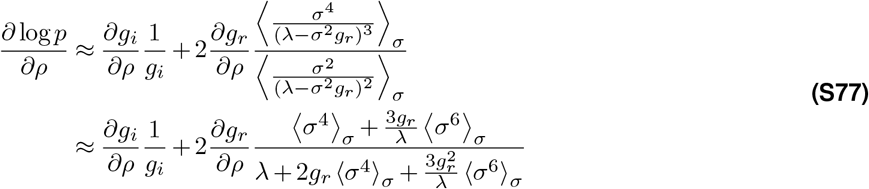

By examining Equations (S68) and (S70), we find that when *λ* ≫ *g*_*r*_, *a* ≫ *b, θ ≈ π, g*_*r*_ decays weakly with *ρ* while *g*_*i*_ increases weakly with *ρ* (also confirmed by numerical calculation, Fig. S25B,C)

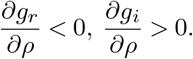

It is therefore straightforward to see from Eq. (S77) that the higher-order moment (e.g., E(*σ*^4^)) in the activity variance contributes to reducing the *ρ*-dependence in the eigendensity function.

#### 6.5.4 The relationship between collapse index (CI) and eigendensity

In this section, we show how the collapse index (CI) introduced in section 4.7 is related to Eq. (S77), namely how the eigendensity changes with the neuronal density in the functional space. Recall the definition of CI in Eq. (13):

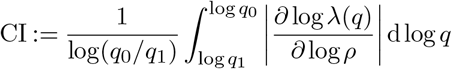

where

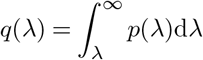

we used implicit differentiation to compute 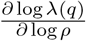. For clarity, we write the function *q*(*λ, ρ*) explicitly involving *λ* and *ρ* as *Q*(*λ, ρ*) in Equations (S78) to (S80).

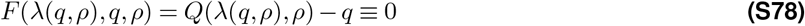

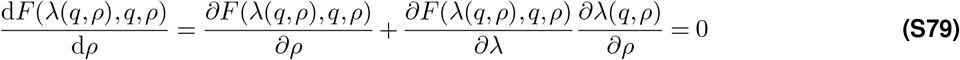

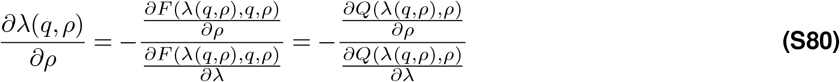

Now we can write CI as

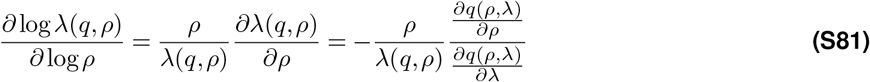

from which we arrive at Eq. (15) in Methods:

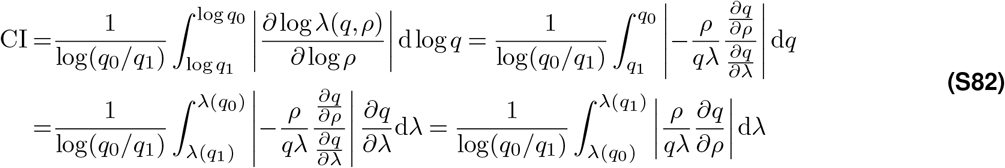

Finally, we can rewrite CI as a function of 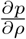 using a double integral:

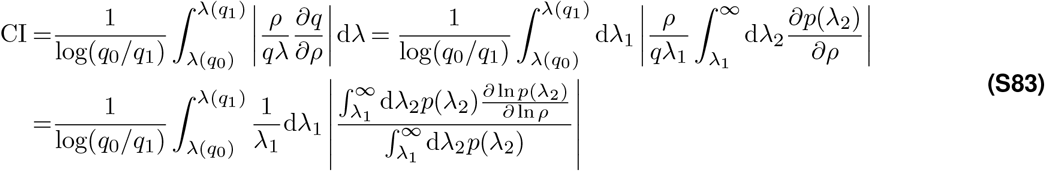

### 6.6 Compare high-density theory and Gaussian variational method

This section aims to determine the conditions under which the high-density approximation aligns with the simulation results. To this end, we begin by comparing the kernel operator 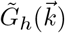 in the high-density quadratic action and 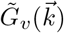 in the variational approximation. We identify the condition when high-density theory would agree with the variational method as well as the numerical simulation, namely 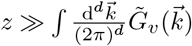. Secondly, we give a precise re-derivation of the high-density result by incorporating this condition into the variational approximation. Finally, we substitute 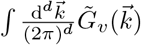 with 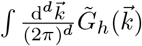 and estimate the parameter regime where the high-density theory would agree with numerical simulation. This analysis yields a deeper understanding of the relationship between high-density theory and variational method, and how they relate to simulation results.

#### 6.6.1 A simple comparison of the two methods

For the sake of simplicity, we consider the correlation matrix with *p*(*σ*) = *δ*(*σ* − 1) in this section. Returning to the explicit result (Equations (S26) to (S28)),

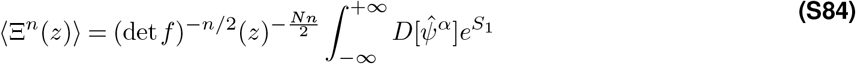

In the high-density approximation (Eq. (S32))

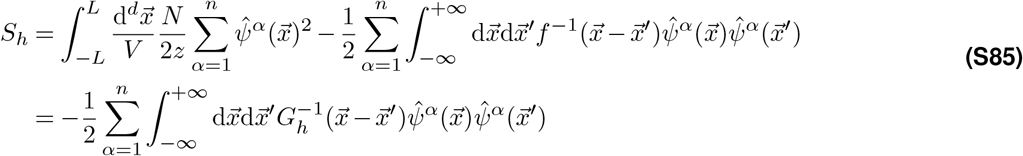

Here we introduce *G*_*h*_ as the kernel operator in the high-density quadratic action.

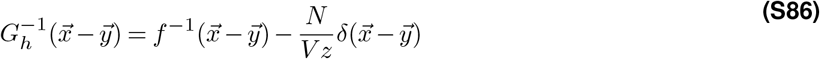

Fourier transform of *G*_*h*_ leads to

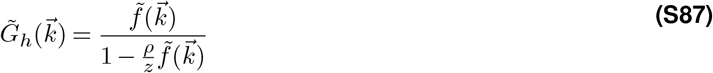

In the variational method (Eq. (S60)), we have

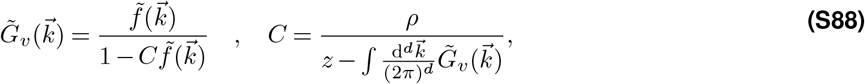

where we introduce *G*_*v*_ as the kernel operator in the variational quadratic action. Clearly, the condition that 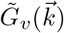 approaches 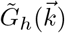 is given by

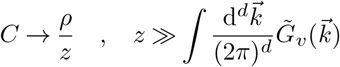

The function *ratio*_*v*_(*z*) is defined as:

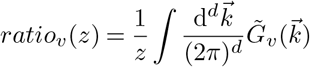

As *ratio*_*v*_(*z*) approaches 0, 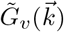 becomes identical to 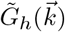. Note that 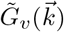 is difficult to compute; instead, we can compute and analyze 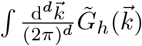 (see section 6.6.3)

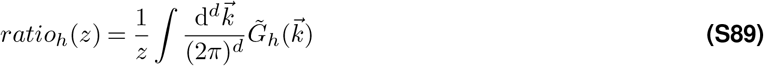

#### 6.6.2 A re-derivation of the high-density result using the grand canonical ensemble

In this section, we re-derive the high-density result from the grand canonical ensemble and the variational method. The derivation also allows us to reproduce the approximation condition discussed in the previous section.

Let us recall the calculation of the free energy *F*_*v*_ (Eq. (S57)) in the variational approximation with *p*(*σ*) = *δ*(*σ* − 1)

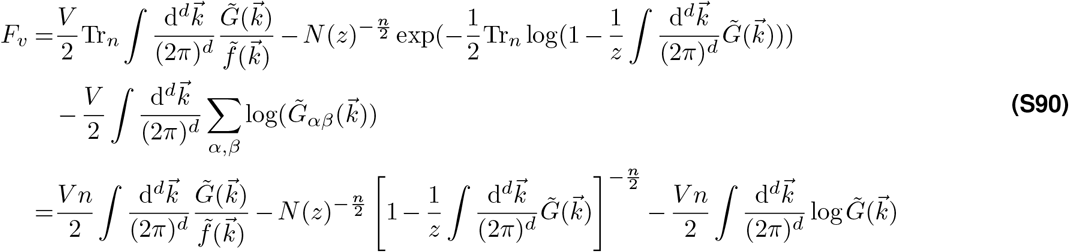

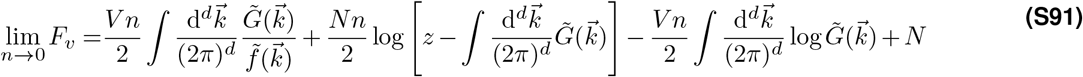

Following Equations (S58) and (S60):

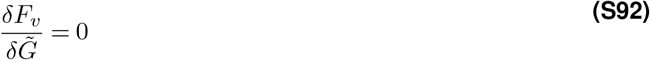

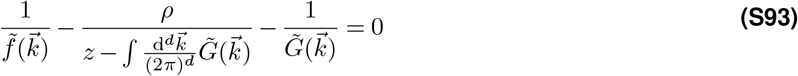

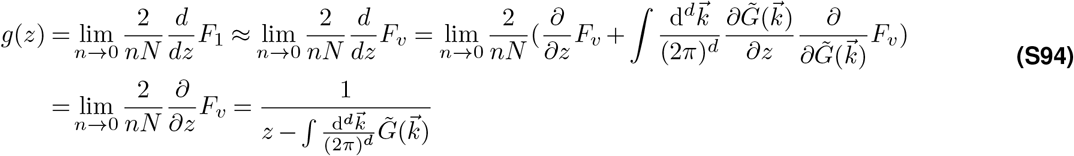

We can perform the same calculation in the high-density theory by considering the limit 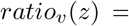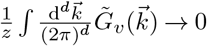

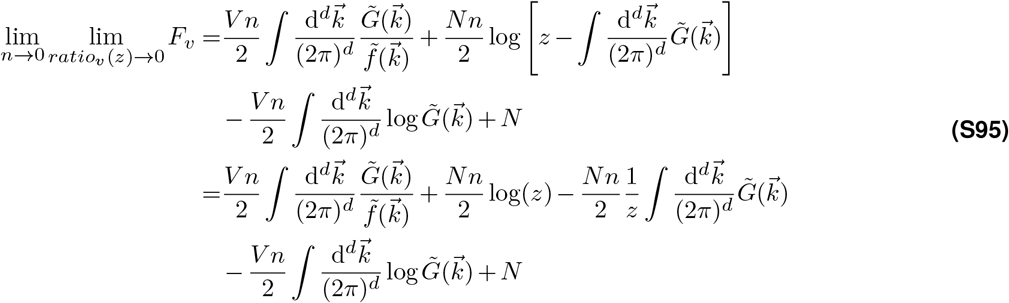

Therefore, we can define the free energy *F*_*h*_ in the high-density theory as

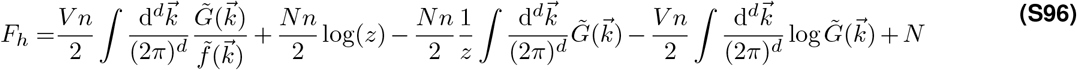

then

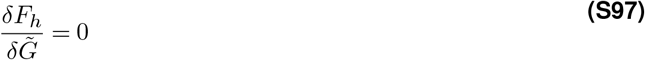

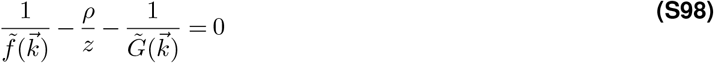

This is precisely Eq. (S87) derived in the previous section.

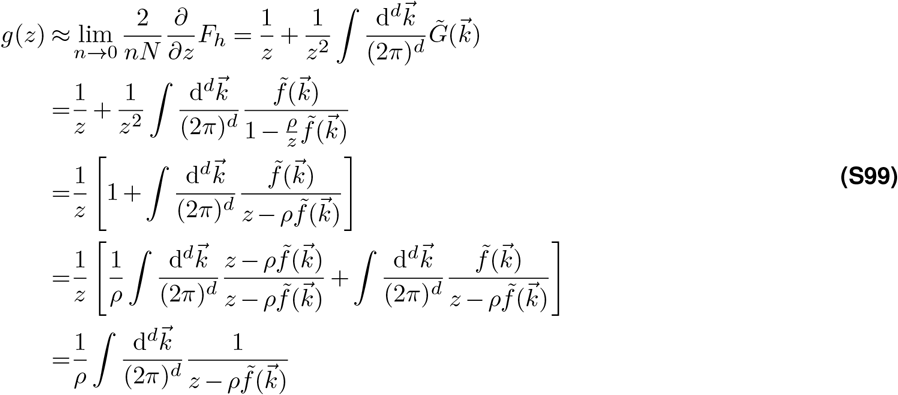

which is the resolvent of high-density approximation (Eq. (S34)).

#### 6.6.3 compute

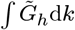 In this section, we provide an explicit expression for the integral 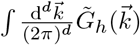 instead of 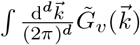, which is implicit and can not be calculated analytically. Like the derivation in section 6.5, we consider the lower and upper limits of integration for 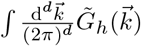 as 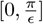 We then approximate the Fourier transform 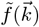 as a power-law function. To ensure that the singularity 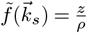 of 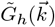 falls within the integration range of 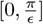,we introduce a simple correction 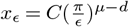 to 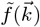:

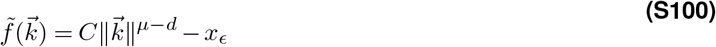

where 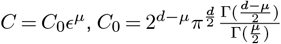 are all constants depending on the parameters, *d, μ*, and *ϵ*.. Then we compute the contour integral (Fig. S25A) by Taylor expansion. As a result, we have

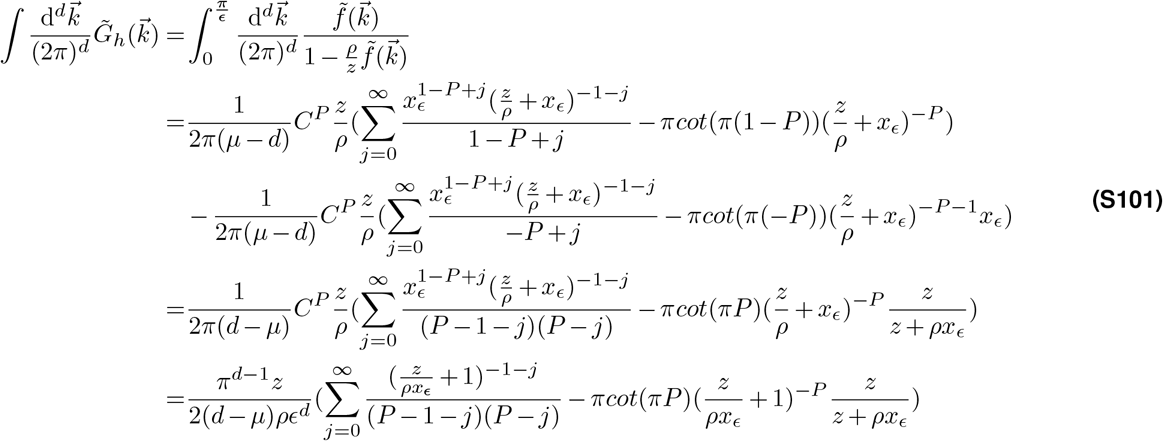

where 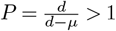

Now let us take a close look at the behavior of the function *ratio*_*h*_(*z*) (Eq. (S89)), plotted in Fig. S26A,B. For small *z*, this function is negative. It then crosses zero and has a peak. As *z*, the *ratio*_*h*_ approaches zero. This is because Eq. (S101) approaches a positive constant, which is given by

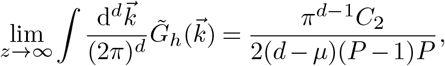

where 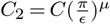. For *z* ≥ 1, we find the leading order expansion at *j* = 1 already gives an accurate approximation (Fig. S26A,B).

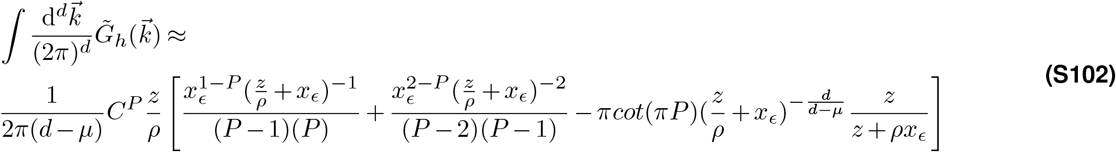

**Figure S26.**
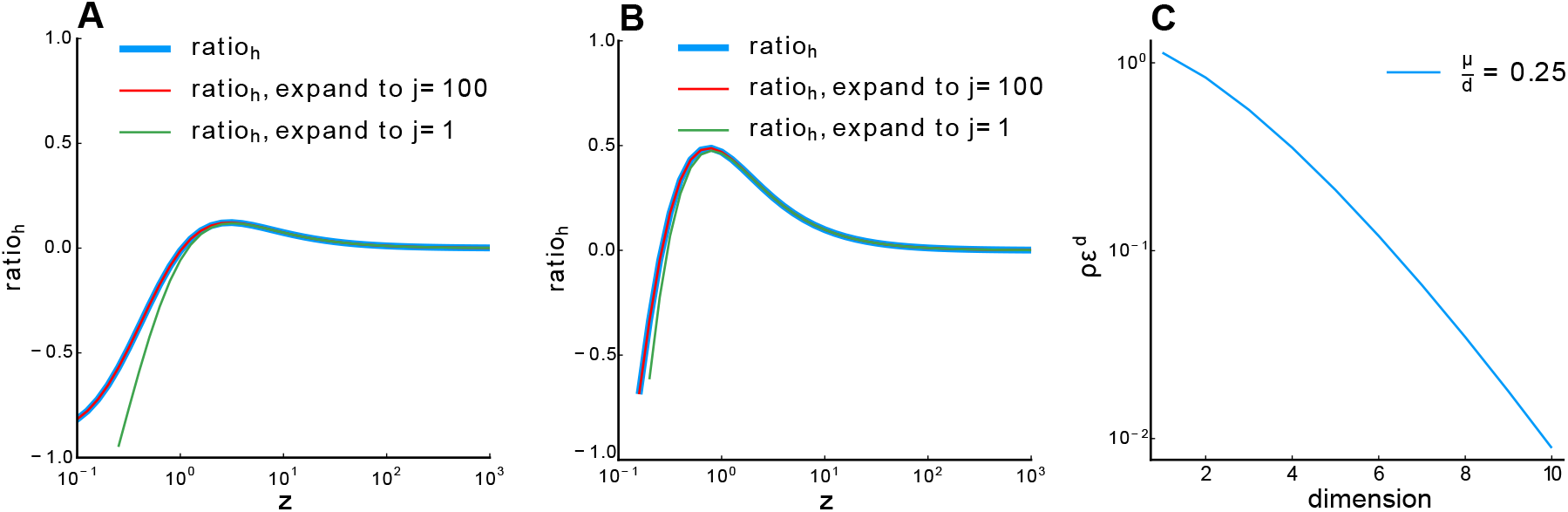
Relationship between *ratio*_*h*_ and *z*. **A**. *ρ* = 1024, **B**. *ρ* = 256. Blue line: *ratio*_*h*_ calculated numerically. Red line: 100-order expansion of Eq. (S101), which perfectly overlaps with the blue line. Green line: expansion to the first order. Other parameter: *µ* = 0.5, *d* = 2, *ϵ* = 0.03125. **C**. Relationship between *ρϵ*^*d*^ and dimension *d* with fixed 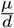 (Eq. (S105)).

#### 6.6.4 Estimate the parameter condition when the high-density theory best agrees with numerical simulation

By analyzing the properties of the function 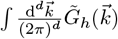, we think the high-density theory provides an accurate approximation when the zero-crossing of 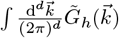 is near *z* = 1 (the peak of low-density result (34))

The root *z*_0_ of the integral 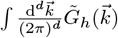 is given by

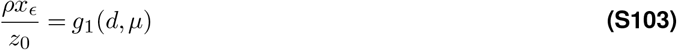

It is easy to see that *g*_1_(*d, µ*) is a function of *P* (or 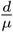) from Eq. (S101). We can rewrite Eq. (S103) as

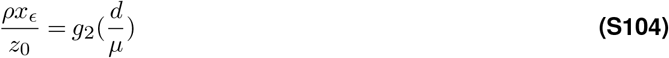

Here, we can also see that *z*_0_ can be expressed as:

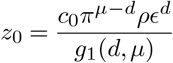

Using this expression for *z*_0_ and letting *z*_0_ = 1, we can derive the following equation for *ρϵ*^*d*^, a *dimensionless parameter* that determines the condition when the high-density theory is an accurate approximation of our ERM model:

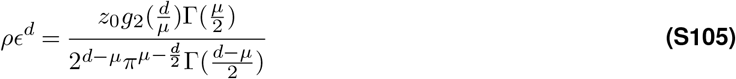

Fig. S26C shows how *ρϵ*^*d*^ changes as a function *d* for a small and fixed *µ/d*. For example, when *d* = 2, *µ* = 0.5, *ϵ* = 0.03125, we find

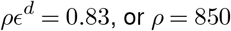

This estimate is also consistent with our numerical simulation (Fig. S3).

### 6.7 Wick rotation

To ensure mathematical rigor in section 6.4, we should make sure that the action *S*_1_ in Eq. (S43) is a real number. Here we use Wick rotation to transform Eq. (S25) to Eq. (S27). The Gaussian integral Eq. (S26) can be divergent when 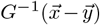 is not positive definite, To address this issue, we can always write the partition function Ξ^*n*^(*z*) as a Gaussian integral by choosing the appropriate axes with Wick rotation.

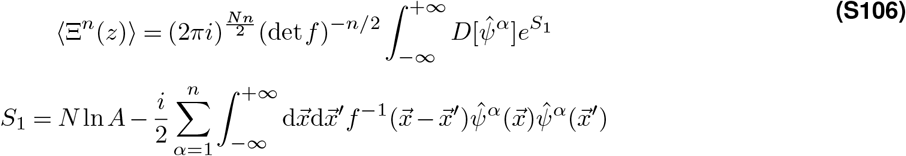

We can now change the integration variables by diagonalizing 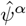 to 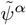 via 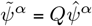,where *Q* is Fourier base

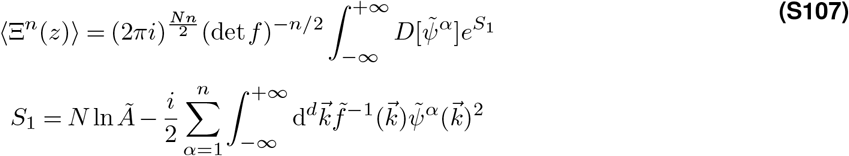

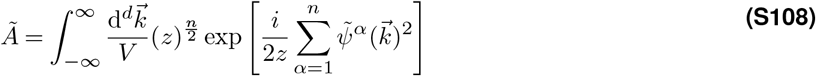

by letting *L* → ∞. Note that 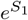 is analytic. Thus if

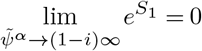

and the convergence rate is faster than 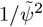, we can apply the Wick rotation 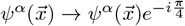: instead of computing the integral on the real axis *C*_1_, we now rotate the integral line 45 degree clockwise to *C*_3_ in the complex plane:

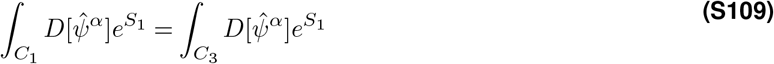

**Figure S27.**
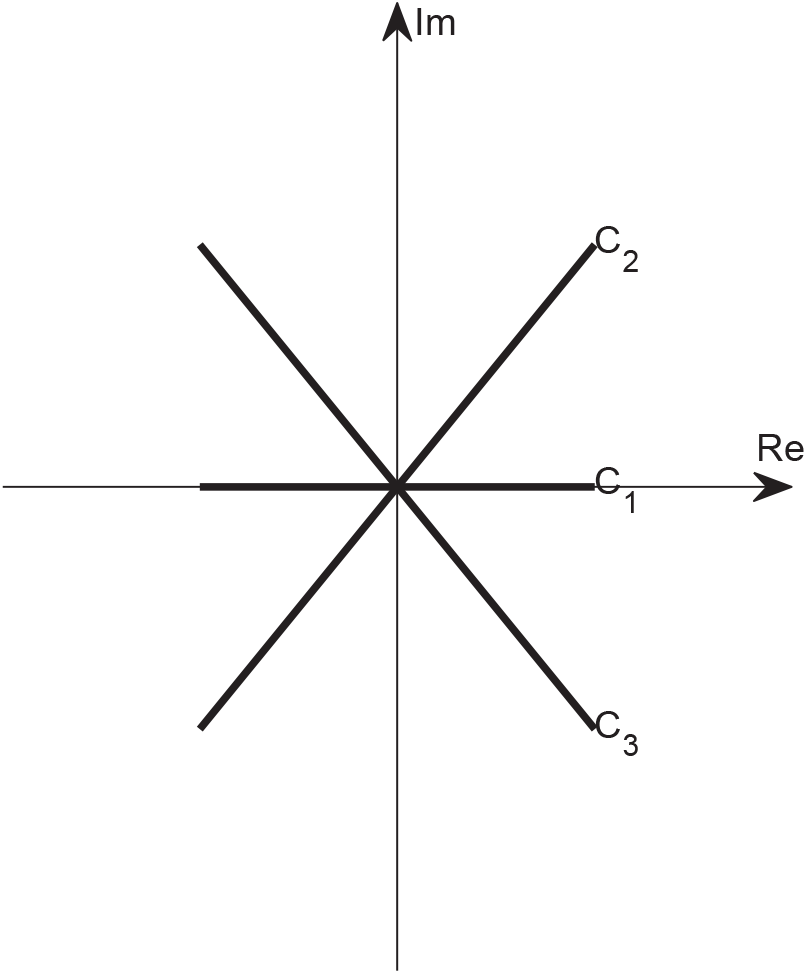
Wick rotation in complex plane.

On the other hand, if

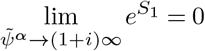

and the convergence rate is faster than 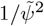, we can apply the Wick rotation 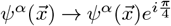, namely to rotate the integral line 45 degree counterclockwise to *C*_2_:

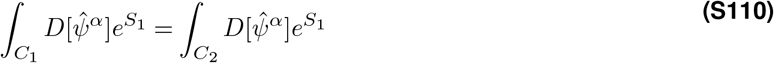

As a simple example, consider a one-dimensional Gaussian integral

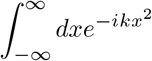

When *k >* 0, we can use the Wick rotation 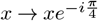

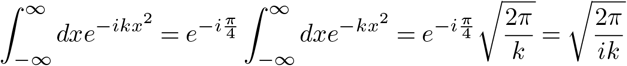

When *k <* 0, we can use the Wick rotation 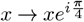

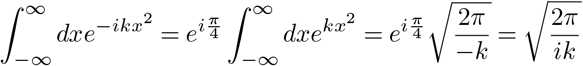

Without loss of generality, we rotate 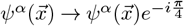 in section 6.4 for subsequent calculations.

### 6.8 Grand Canonical Ensemble

When using the Gaussian variational Approximation, we consider a critical extension from the *canonical ensemble* to the *grand canonical ensemble* when computing the partition function (Eq. (S6)). We would like to justify this approximation in this section. Recall that the resolvent is given by

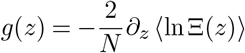

where Ξ(*z*) can be viewed as the canonical partition function, the ⟨*…* ⟩ is the average over all random matrices *C* for a given *N*. Let us now generalize (Eq. (S6)) into grand canonical ensemble, namely

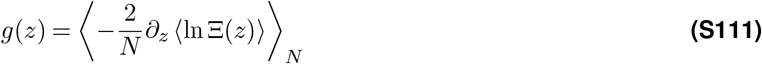

where ⟨*…* ⟩_*N*_ indicates that we need to average over all possible random matrices and across *all possible N*, with the probability to have a matrix size *N* given by the Poisson distribution 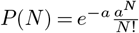, where *a* = ⟨*N*⟩. When ⟨*N*⟩ is large, *P*(*N*) has a very sharp peak at ⟨*N*⟩, and Eq. (S111) can be approximated as

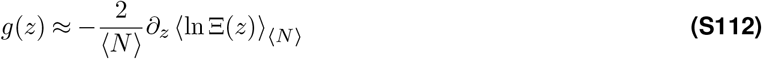

Using the replica trick, we recall Eq. (S9)

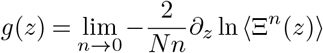

Let us now define the grand canonical partition function as

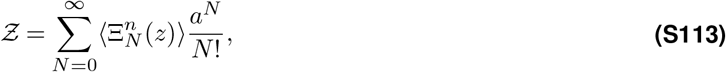

Likewise, the resolvent in Eq. (S9) is generalized to

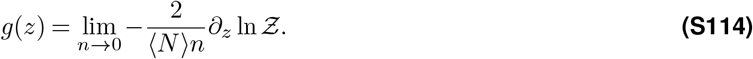

To see whether this definition makes sense, we write

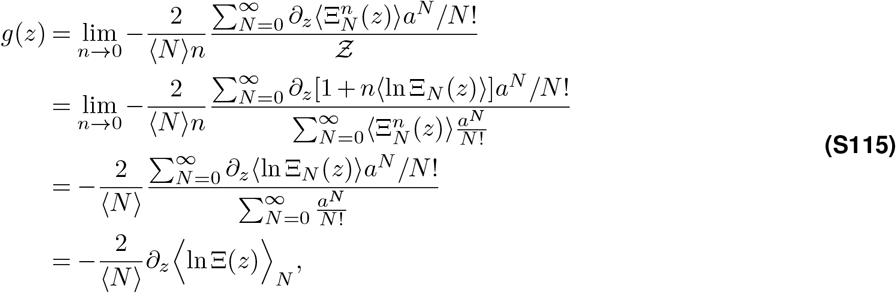

where the second equality uses the identity

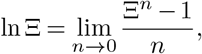

and the last equality is indeed Eq. (S112) discussed earlier.

Returning back to the explicit form of the grand canonical partition function in our ERM model (Equations (S26) to (S28)), we have

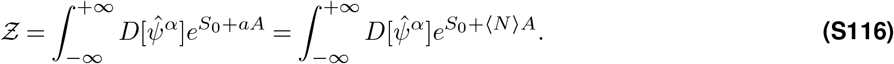

Here *ψ* is the auxiliary fields (Eq. (S12)),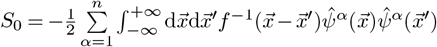 and are terms defined in Equations (S26) to (S28). Eq. (S116) is used in section 6.4 to compute the free energy.

### 6.9 E-I balanced asynchronized model Summary

In this section, we discuss the E-I balanced asynchronized model (53), which predicts a different scaling D N under random sampling, since the variance 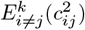 scales as 1/N and diminishes as N approaches large limit.

#### 6.9.1 Model

The simulation of binary networks involves updating neuron states within a network architecture identical to analytical studies. The update rule is probabilistic, with neuron activities set based on synaptic currents and a firing threshold. The dynamics resolution improves with network size, with neuron time constants effectively representing changes in firing activity. Parameters for simulations include connection probabilities, mean rates, thresholds, and synaptic strengths, scaled appropriately for network size.

Update Rule: 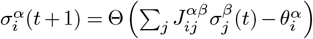

Dynamics Resolution: 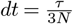

In the simulation of binary networks, the model’s dynamics are governed by a set of parameters, each with a specific role:

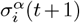: This represents the state of neuron *i* in population *α* at the next time step *t* + 1. The state is binary, where 1 indicates the neuron is active (firing) and 0 indicates it is inactive.

Θ(): The Heaviside step function used in the update rule. It determines the neuron’s next state by comparing the net input to the neuron against its firing threshold. If the net input exceeds the threshold, the neuron’s state is set to active; otherwise, it remains or becomes inactive.

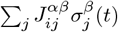: This sum represents the total synaptic input to neuron *i* from all neurons *j* in population *β* at time *t. j* at time *t*.

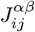 is the synaptic weight from neuron *j* in population *β* to neuron *i* in population *α*, and 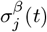 is the state of neuron *j* at time *t*.

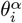: The firing threshold of neuron *i* in population *α*. It is the value against which the net synaptic input is compared to determine whether neuron *i* will fire (transition to state 1) or not (remain in state 0).

*α* = {*E, I*}, *β* = {*E, I, X}* : Represents a specific population of neurons within the network. E: excitatory neurons, I: inhibitory neurons, or X: external source of neurons that provide inputs to the network but are not influenced by the network’s internal dynamics.

#### 6.9.2 Firing Rate Correlation r

The mean firing rate correlation *E*(*r*) scales inversely with the network size *N*, specifically, *E*(*r*) ∼ 1*/N*. The standard deviation *σ*_*r*_ of *r* decays only as 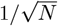 (53).

Given that the variance of *r*, denoted as Var(*r*), is 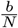, and the expected value of *r*, denoted as *E*(*r*), is 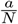, where *N* is the sample size, and *a* and *b* are constants, we aim to calculate *E*(*r*^*2*^), the expected value of the square of the correlation coefficient *r*.

The term 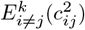 in PR dimension is given by:

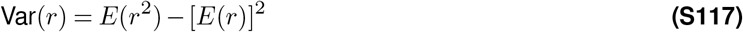

Substituting 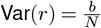 and 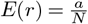 into the equation, we get:

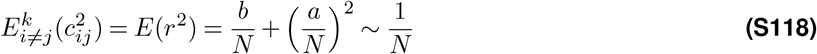

Thus in PR dimension 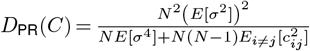, the term 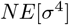 and 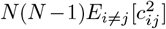 are of the same order, and the PR dimension will not reach the upper bound 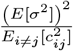.

## Supplementary videos

**Movie S1.**
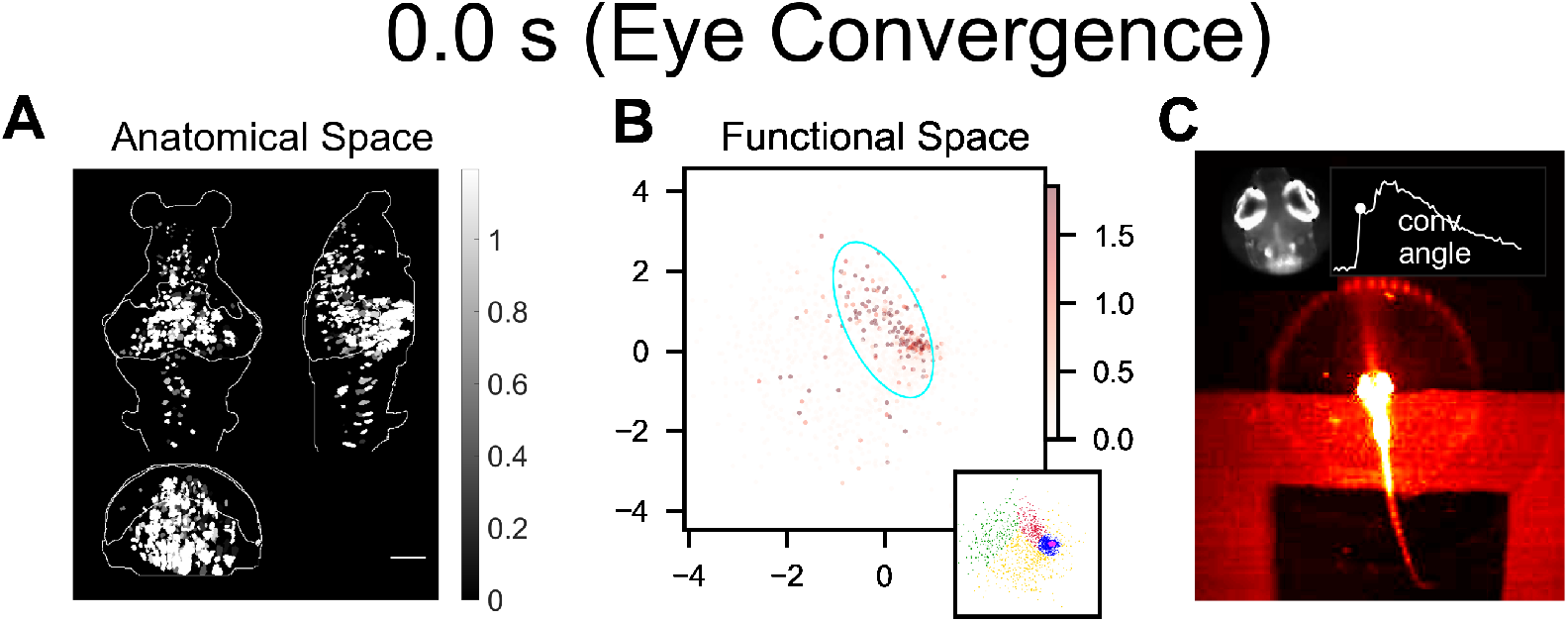
Neural activity patterns in anatomical and functional space during hunting (click here). Single-trial examples of fish 1 and fish 3. **A**. Inferred firing rate activity in anatomical space. Scale bar, 100 µm. **B**. Inferred firing rate activity in functional space. Functional space organization of the control data inferred by fitting the ERM and MDS in section 2.4. The cyan ellipse serves as a visual aid for the cluster size: it encloses 95% of the neurons belonging to that cluster (Methods). The inset illustrates the functional space organization, similar to that shown in Fig. S15C. The colorbars in panels **A** and **B** depict the inferred activity magnitude of individual neurons. **C**. Simultaneous behavior recording alongside the neural activity. Time, seconds.

## References

1. Churchland, M. M., Cunningham, J. P., Kaufman, M. T., Foster, J. D., Nuyujukian, P., Ryu, S. I., and Shenoy, K. V. Neural population dynamics during reaching. Nature, 487(7405):51–56, July 2012. doi: 10.1038/nature11129.

2. Zhang, H., Rich, P. D., Lee, A. K., and Sharpee, T. O. Hippocampal spatial representations exhibit a hyperbolic geometry that expands with experience. Nature Neuroscience, 26(1):131–139, Jan. 2023. doi: 10.1038/s41593-022-01212-4.

3. Kriegeskorte, N. and Wei, X.-X. Neural tuning and representational geometry. Nature Reviews Neuroscience, 22(11): 703–718, 2021. doi: 10.1038/s41583-021-00502-3.

4. Chung, S. and Abbott, L. F. Neural population geometry: An approach for understanding biological and artificial neural networks. Current opinion in neurobiology, 70:137–144, 2021.

5. Cunningham, J. P. and Yu, B. M. Dimensionality reduction for large-scale neural recordings. Nature Neuroscience, 17(11): 1500–1509, Nov. 2014. doi: 10.1038/nn.3776.

6. Stringer, C., Pachitariu, M., Steinmetz, N., Carandini, M., and Harris, K. D. High-dimensional geometry of population responses in visual cortex. Nature, 2019. doi: 10.1038/s41586-019-1346-5.

7. Si, G., Kanwal, J. K., Hu, Y., Tabone, C. J., Baron, J., Berck, M., Vignoud, G., and Samuel, A. D. Structured odorant response patterns across a complete olfactory receptor neuron population. Neuron, 101(5):950–962.e7, Mar. 2019. doi: 10.1016/j.neuron.2018.12.030.

8. Mante, V., Sussillo, D., Shenoy, K. V., and Newsome, W. T. Context-dependent computation by recurrent dynamics in prefrontal cortex. Nature, 503(7474):78–84, Nov. 2013. doi: 10.1038/nature12742.

9. Yang, W., Tipparaju, S. L., Chen, G., and Li, N. Thalamus-driven functional populations in frontal cortex support decision-making. Nature Neuroscience, 25(10):1339–1352, Oct. 2022. doi: 10.1038/s41593-022-01171-w.

10. Xie, Y., Hu, P., Li, J., Chen, J., Song, W., Wang, X.-J., Yang, T., Dehaene, S., Tang, S., Min, B., and Wang, L. Geometry of sequence working memory in macaque prefrontal cortex. Science, 375(6581):632–639, Feb. 2022. doi: 10.1126/science.abm0204.

11. Nguyen, J. P., Shipley, F. B., Linder, A. N., Plummer, G. S., Liu, M., Setru, S. U., Shaevitz, J. W., and Leifer, A. M. Whole-brain calcium imaging with cellular resolution in freely behaving Caenorhabditis elegans. Proceedings of the National Academy of Sciences, 113(8):E1074–E1081, Feb. 2016. doi: 10.1073/pnas.1507110112.

12. Lindén, H., Petersen, P. C., Vestergaard, M., and Berg, R. W. Movement is governed by rotational neural dynamics in spinal motor networks. Nature, 610(7932):526–531, Oct. 2022. doi: 10.1038/s41586-022-05293-w.

13. Urai, A. E., Doiron, B., Leifer, A. M., and Churchland, A. K. Large-scale neural recordings call for new insights to link brain and behavior. Nature Neuroscience, 25(1):11–19, Jan. 2022. doi: 10.1038/s41593-021-00980-9.

14. Buzsáki, G. Large-scale recording of neuronal ensembles. Nature Neuroscience, 7(5):446–451, May 2004. doi: 10.1038/nn1233.

15. Ahrens, M. B., Li, J. M., Orger, M. B., Robson, D. N., Schier, A. F., Engert, F., and Portugues, R. Brain-wide neuronal dynamics during motor adaptation in zebrafish. Nature, 485(7399):471–477, May 2012. doi: 10.1038/nature11057.

16. Jun, J. J., Steinmetz, N. A., Siegle, J. H., Denman, D. J., Bauza, M., Barbarits, B., Lee, A. K., Anastassiou, C. A., Andrei, A., Aydin Barbic, M., Blanche, T. J., Bonin, V., Couto, J., Dutta, B., Gratiy, S. L., Gutnisky, D. A., Häusser, M., Karsh, B., Ledochowitsch, P., Lopez, C. M., Mitelut, C., Musa, S., Okun, M., Pachitariu, M., Putzeys, J., Rich, P. D., Rossant, C., Sun, W.-l., Svoboda, K., Carandini, M., Harris, K. D., Koch, C., O’Keefe, J., and Harris, T. D. Fully integrated silicon probes for high-density recording of neural activity. Nature, 551(7679):232–236, Nov. 2017. doi: 10.1038/nature24636.

17. Stevenson, I. H. and Kording, K. P. How advances in neural recording affect data analysis. Nature neuroscience, 14(2): 139–142, 2011.

18. Sofroniew, N. J., Flickinger, D., King, J., and Svoboda, K. A large field of view two-photon mesoscope with subcellular resolution for in vivo imaging. eLife, 5:e14472, June 2016. doi: 10.7554/eLife.14472.

19. Lin, A., Witvliet, D., Hernandez-Nunez, L., Linderman, S. W., Samuel, A. D., and Venkatachalam, V. Imaging whole-brain activity to understand behaviour. Nature Reviews Physics, 4(5):292–305, 2022.

20. Meshulam, L., Gauthier, J. L., Brody, C. D., Tank, D. W., and Bialek, W. Coarse graining, fixed points, and scaling in a large population of neurons. Physical Review Letters, 123:178103, 2019. doi: 10.1103/PhysRevLett.123.178103.

21. Demas, J., Manley, J., Tejera, F., Barber, K., Kim, H., Traub, F. M., Chen, B., and Vaziri, A. High-speed, cortex-wide volumetric recording of neuroactivity at cellular resolution using light beads microscopy. Nature Methods, 18(9):1103–1111, Sept. 2021. doi: 10.1038/s41592-021-01239-8.

22. Musall, S., Kaufman, M. T., Juavinett, A. L., Gluf, S., and Churchland, A. K. Single-trial neural dynamics are dominated by richly varied movements. Nature Neuroscience, 22(10):1677–1686, Oct. 2019. doi: 10.1038/s41593-019-0502-4.

23. Stringer, C., Pachitariu, M., Steinmetz, N., Reddy, C. B., Carandini, M., and Harris, K. D. Spontaneous behaviors drive multidimensional, brainwide activity. Science, 364(6437):eaav7893, Apr. 2019. doi: 10.1126/science.aav7893.

24. Kleinfeld, D., Luan, L., Mitra, P. P., Robinson, J. T., Sarpeshkar, R., Shepard, K., Xie, C., and Harris, T. D. Can one concurrently record electrical spikes from every neuron in a mammalian brain? Neuron, 103(6):1005–1015, 2019.

25. Recanatesi, S., Ocker, G. K., Buice, M. A., and Shea-Brown, E. Dimensionality in recurrent spiking networks: Global trends in activity and local origins in connectivity. PLoS computational biology, 15(7):e1006446, 2019.

26. Litwin-Kumar, A., Harris, K. D., Axel, R., Sompolinsky, H., and Abbott, L. Optimal degrees of synaptic connectivity. Neuron, 93(5):1153–1164, 2017.

27. Gao, P., Trautmann, E., Yu, B., Santhanam, G., Ryu, S., Shenoy, K., and Ganguli, S. A theory of multineuronal dimensionality, dynamics and measurement. BioRxiv, page 214262, 2017.

28. Clark, D. G., Abbott, L., and Litwin-Kumar, A. Dimension of activity in random neural networks. Physical Review Letters, 131 (11):118401, 2023.

29. Dahmen, D., Recanatesi, S., Ocker, G. K., Jia, X., Helias, M., and Shea-Brown, E. Strong coupling and local control of dimensionality across brain areas. Biorxiv, pages 2020–11, 2020.

30. Cong, L., Wang, Z., Chai, Y., Hang, W., Shang, C., Yang, W., Bai, L., Du, J., Wang, K., and Wen, Q. Rapid whole brain imaging of neural activity in freely behaving larval zebrafish (Danio rerio). eLife, 6:e28158, Sept. 2017. doi: 10.7554/eLife.28158.

31. Hu, Y. and Sompolinsky, H. The spectrum of covariance matrices of randomly connected recurrent neuronal networks with linear dynamics. PLoS Computational Biology, 18, 7 2022. doi: 10.1371/journal.pcbi.1010327.

32. Morales, G. B., di Santo, S., and Muñoz, M. A. Quasiuniversal scaling in mouse-brain neuronal activity stems from edge-of-instability critical dynamics. Proceedings of the National Academy of Sciences, 120(9):e2208998120, Feb. 2023. doi: 10.1073/pnas.2208998120.

33. Chen, X., Mu, Y., Hu, Y., Kuan, A. T., Nikitchenko, M., Randlett, O., Chen, A. B., Gavornik, J. P., Sompolinsky, H., Engert, F., and Ahrens, M. B. Brain-wide organization of neuronal activity and convergent sensorimotor transformations in larval zebrafish. Neuron, 100(4):876–890.e5, Nov. 2018. doi: 10.1016/j.neuron.2018.09.042.

34. Mézard, M., Parisi, G., and Zee, A. Spectra of euclidean random matrices. Nuclear Physics B, 559(3):689–701, Oct. 1999. doi: 10.1016/S0550-3213(99)00428-9.

35. Goetschy, A. and Skipetrov, S. Euclidean random matrices and their applications in physics. arXiv preprint arXiv:1303.2880, 2013. doi: 10.48550/ARXIV.1303.2880.

36. Morrell, M. C., Nemenman, I., and Sederberg, A. Neural criticality from effective latent variables. eLife, 12:RP89337, Mar. 2024. doi: 10.7554/eLife.89337.3.

37. Hubel, D. H. and Wiesel, T. N. Receptive fields of single neurones in the cat’s striate cortex. The Journal of physiology, 148 (3):574, 1959.

38. Stefanini, F., Kushnir, L., Jimenez, J. C., Jennings, J. H., Woods, N. I., Stuber, G. D., Kheirbek, M. A., Hen, R., and Fusi, S. A distributed neural code in the dentate gyrus and in CA1. Neuron, 107(4):703–716.e4, Aug. 2020. doi: 10.1016/j.neuron.2020.05.022.

39. Kropff, E., Carmichael, J. E., Moser, M.-B., and Moser, E. I. Speed cells in the medial entorhinal cortex. Nature, 523(7561): 419–424, July 2015. doi: 10.1038/nature14622.

40. O’Keefe, J. Place units in the hippocampus of the freely moving rat. Experimental Neurology, 51(1):78–109, Jan. 1976. doi: 10.1016/0014-4886(76)90055-8.

41. Moser, E. I., Kropff, E., and Moser, M.-B. Place cells, grid cells, and the brain’s spatial representation system. Annual Review of Neuroscience, 31(1):69–89, July 2008. doi: 10.1146/annurev.neuro.31.061307.090723.

42. Tingley, D. and Buzsáki, G. Transformation of a Spatial Map across the Hippocampal-Lateral Septal Circuit. Neuron, 98(6): 1229–1242.e5, June 2018. doi: 10.1016/j.neuron.2018.04.028.

43. Tian, G. J., Zhu, O., Shirhatti, V., Greenspon, C. M., Downey, J. E., Freedman, D. J., and Doiron, B. Neuronal firing rate diversity lowers the dimension of population covariability. bioRxiv, 2024. doi: 10.1101/2024.08.30.610535.

44. Grewe, B. F. and Helmchen, F. Optical probing of neuronal ensemble activity. Current Opinion in Neurobiology, 19(5): 520–529, Oct. 2009. doi: 10.1016/j.conb.2009.09.003.

45. Gauthier, J. L. and Tank, D. W. A dedicated population for reward coding in the hippocampus. Neuron, 99(1):179–193.e7, July 2018. doi: 10.1016/j.neuron.2018.06.008.

46. Cox, T. and Cox, M. Multidimensional Scaling. Chapman and Hall/CRC, 0 edition, Sept. 2000. ISBN 978-0-367-80170-0. doi: 10.1201/9780367801700. URL https://www.taylorfrancis.com/books/9781420036121.

47. Bianco, I. H., Kampff, A. R., and Engert, F. Prey capture behavior evoked by simple visual stimuli in larval zebrafish. Frontiers in Systems Neuroscience, 5, 2011. doi: 10.3389/fnsys.2011.00101.

48. Kunst, M., Laurell, E., Mokayes, N., Kramer, A., Kubo, F., Fernandes, A. M., Förster, D., Dal Maschio, M., and Baier, H. A cellular-resolution atlas of the larval zebrafish brain. Neuron, 103(1):21–38, 2019.

49. Kardar, M. Statistical Physics of Fields. Cambridge University Press, Cambridge, 2007. ISBN 978-0-521-87341-3. doi: 10.1017/CBO9780511815881. URL https://www.cambridge.org/core/books/statistical-physics-of-fields/06F49D11030FB3108683F413269DE945.

50. Beggs, J. M. and Plenz, D. Neuronal avalanches in neocortical circuits. Journal of Neuroscience, 23(35):11167–11177, Dec. 2003. doi: 10.1523/JNEUROSCI.23-35-11167.2003.

51. Dahmen, D., Grün, S., Diesmann, M., and Helias, M. Second type of criticality in the brain uncovers rich multiple-neuron dynamics. Proceedings of the National Academy of Sciences, 116(26):13051–13060, June 2019. doi: 10.1073/pnas.1818972116.

52. Morrell, M. C., Sederberg, A. J., and Nemenman, I. Latent dynamical variables produce signatures of spatiotemporal criticality in large biological systems. Physical Review Letters, 126:118302, 2021. doi: 10.1103/PhysRevLett.126.118302.

53. Renart, A., De La Rocha, J., Bartho, P., Hollender, L., Parga, N., Reyes, A., and Harris, K. D. The asynchronous state in cortical circuits. science, 327(5965):587–590, 2010.

54. Manley, J., Lu, S., Barber, K., Demas, J., Kim, H., Meyer, D., Traub, F. M., and Vaziri, A. Simultaneous, cortex-wide dynamics of up to 1 million neurons reveal unbounded scaling of dimensionality with neuron number. Neuron, 2024.

55. Hoffmann, M., Henninger, J., Veith, J., Richter, L., and Judkewitz, B. Blazed oblique plane microscopy reveals scale-invariant inference of brain-wide population activity. Nature Communications, 14(1):8019, Dec. 2023. doi: 10.1038/s41467-023-43741-x.

56. Moosavi, S. A., Hindupur, S. S. R., and Shimazaki, H. Population coding under the scale-invariance of high-dimensional noise, Aug. 2024. URL 10.1101/2024.08.23.608710.

57. Mathis, A., Mamidanna, P., Cury, K. M., Abe, T., Murthy, V. N., Mathis, M. W., and Bethge, M. DeepLabCut: markerless pose estimation of user-defined body parts with deep learning. Nature Neuroscience, 21(9):1281–1289, Sept. 2018. doi: 10.1038/s41593-018-0209-y.

58. Tabor, K. M., Marquart, G. D., Hurt, C., Smith, T. S., Geoca, A. K., Bhandiwad, A. A., Subedi, A., Sinclair, J. L., Rose, H. M., Polys, N. F., and Burgess, H. A. Brain-wide cellular resolution imaging of Cre transgenic zebrafish lines for functional circuit-mapping. eLife, 8:e42687, Feb. 2019. doi: 10.7554/eLife.42687.

59. Studholme, C., Hill, D. L. G., and Hawkes, D. J. Automated three-dimensional registration of magnetic resonance and positron emission tomography brain images by multiresolution optimization of voxel similarity measures. Medical Physics, 24(1):25–35, 1997. doi: 10.1118/1.598130.

60. Rueckert, D., Sonoda, L., Hayes, C., Hill, D., Leach, M., and Hawkes, D. Nonrigid registration using free-form deformations: application to breast MR images. IEEE Transactions on Medical Imaging, 18(8):712–721, Aug. 1999. doi: 10.1109/42.796284.

61. Friedrich, J., Zhou, P., and Paninski, L. Fast online deconvolution of calcium imaging data. PLOS Computational Biology, 13 (3):e1005423, Mar. 2017. doi: 10.1371/journal.pcbi.1005423.

62. Gao, P. and Ganguli, S. On simplicity and complexity in the brave new world of large-scale neuroscience. Current opinion in neurobiology, 32:148–155, 2015.

63. Bordenave, C. Eigenvalues of Euclidean random matrices. Random Structures and Algorithms, 33(4):515–532, Dec. 2008. doi: 10.1002/rsa.20228.

64. Rudin, W. Fourier Analysis on Groups. Wiley, 1 edition, Jan. 1990. ISBN 978-0-470-74481-9 978-1-118-16562-1. doi: 10.1002/9781118165621. URL https://onlinelibrary.wiley.com/doi/book/10.1002/9781118165621.

65. Knapp, T. R. Canonical correlation analysis: a general parametric significance-testing system. Psychological Bulletin, 85(2): 410, 1978.

66. Bradde, S. and Bialek, W. PCA meets RG. Journal of Statistical Physics, 167(3):462–475, May 2017. doi: 10.1007/s10955-017-1770-6.

67. Meshulam, L., Gauthier, J. L., Brody, C. D., Tank, D. W., and Bialek, W. Coarse–graining and hints of scaling in a population of 1000+ neurons. arXiv preprint arXiv:1812.11904, 2018. doi: 10.48550/arXiv.1812.11904.

